# Transcriptional profiling reveals a subset of human breast tumors that retain wt *TP53* but display mutant p53-associated features

**DOI:** 10.1101/2020.04.30.070037

**Authors:** Gal Benor, Garold Fuks, Suet-Feung Chin, Oscar Rueda, Saptaparna Mukherjee, Sharathchandra Arandkar, Yael Aylon, Carlos Caldas, Eytan Domany, Moshe Oren

## Abstract

*TP53* gene mutations are very common in human cancer. While such mutations abrogate the tumor suppressive activities of the wild type (wt) p53 protein, some of them also endow the mutant protein with oncogenic gain-of-function (GOF), facilitating cancer progression. Yet, p53 may acquire altered functionality even without being mutated; in particular, experiments with cultured cells revealed that wt p53 can be rewired to adopt mutant-like features in response to growth factors or cancer-mimicking genetic manipulations. To assess whether such rewiring also occurs in human tumors, we interrogated gene expression profiles and pathway deregulation patterns in the METABRIC breast cancer (BC) dataset as a function of *TP53* gene mutation status. Harnessing the power of machine learning, we optimized a gene expression classifier for ER+Her2- patients that distinguishes tumors carrying *TP53* mutations from those retaining wt *TP53*. Interestingly, a small subset of wt *TP53* tumors displayed gene expression and pathway deregulation patterns markedly similar to those of *TP53*-mutated tumors. Moreover, similar to *TP53* mutated tumors, these “pseudomutant” cases displayed a signature for enhanced proliferation and had worse prognosis than typical wt p53 tumors. Notably, these tumors revealed upregulation of genes which, in BC cell lines, were reported to be positively regulated by p53 GOF mutants. Thus, such tumors may benefit from mutant p53-associated activities without having to accrue *TP53* mutations.

## 1 Introduction

One out of eight women is likely to develop breast cancer during her lifetime (DeSantis *et al*., 2017). BC accounts for 30% of all estimated new cancer cases in women and 25% of all estimated cancer-related women’s deaths. A major challenge in treating BC is its heterogeneity, comprising a large variety of subtypes (Sorlie *et al*., 2001). Understanding each individual patient’s disease better, by its molecular and genomic characterization, is a prerequisite for cancer precision medicine, increasing treatment efficacy while minimizing side effects, and eventually reducing BC mortality (Deng and Nakamura, 2017). With the advent of the big data era, harnessing machine learning towards the optimization of individualized treatments bears great promise for the future of cancer therapy.

The p53 protein, encoded by the *TP53* tumor suppressor gene, has a central role in safeguarding cells against cancer. Over half of all human cancers carry *TP53* mutations, which may be associated with poor prognosis, increased treatment resistance and relapse (Silwal-Pandit *et al*., 2014). The most obvious consequence of such mutations is loss of the tumor suppressive activity of the wt p53 protein. However, some *TP53* mutations may also facilitate cancer progression by endowing the mutant p53 protein with oncogenic GOF (Brosh and Rotter, 2009). Such GOF, manifested by an increase in proliferation, cell motility, therapy resistance and more, is driven mainly by interactions of the mutant p53 with a variety of other proteins, eventually altering gene expression (E. Kim *et al*., 2009; M. P. Kim and Lozano, 2018). Yet, about half of all human tumors retain wt p53 expression. How do such tumors evade p53’s tumor suppressive effects? In some cell types or biological contexts p53 might lose its tumor suppressive capabilities, alleviating the need to override its effects (E. Kim *et al*., 2009). Alternatively, p53 may not reach sufficient steady-state levels because of suppressed transcription of the *TP53* gene (Miller *et al*., 2005), reduced translation, or rapid protein turnover, e.g. via augmented MDM2-mediated degradation (Haupt *et al*., 1997; Kubbutat *et al*., 1997). Additionally, wt p53 may undergo aberrant post-translational modifications or be excluded from the cell nucleus, depriving it of its ability to act as a transcription factor. All these mechanisms may result in p53 loss of function (LOF), equivalent to genetic loss of both wt *TP53* alleles.

Nevertheless, wt p53 may sometimes not only lose its normal activity, but also acquire structural and functional properties resembling those of *bona fide* GOF mutant p53 proteins (Furth *et al*., 2015, 2018; Milner, 1995; Trinidad *et al*., 2013; Zhang and Deisseroth, 1994). Such “pseudomutant” (PM) wt p53 may become an active contributor to cancer (Furth *et al*., 2018). An early study by Miller et al (Miller *et al*., 2005), looking at data from 251 breast tumors, identified a group of wt p53 cases that were labeled as mutant on the basis of their gene expression patterns.

Of note, the study of Miller et al was performed on a mix of tumors of all breast cancer subtypes, analyzed together as one population. However, the relative abundance of *TP53* mutations varies greatly between subtypes; for example, ER+ tumors are predominantly wt *TP53*, whereas a large proportion of ER- tumors carry *TP53* mutations. Thus, part of the reported differences in transcription profiles between wt and mutant p53 were most probably due to their unequal representation in the different breast cancer subtypes. Indeed, the wt p53 transcriptional signature defined by Miller et al was enriched in estrogen-inducible genes (Miller *et al*., 2005), which might simply reflect the fact that the great majority of wt p53 tumors are ER+.

To avoid a bias introduced by the unequal frequency of *TP53* mutations in the different BC subtypes, we chose to compare the expression patterns of wt and mutant *TP53* tumors only within the same subtype. We believe that this approach eliminates the “noise” occurring from comparisons between different subclasses of disease and thus enables rigorous identification of wt *TP53* tumors that indeed behave in a mutant-like manner. We interrogated each breast cancer subtype separately for the existence of “pseudomutant” (PM) tumors that harbor wt *TP53* by sequence, but nevertheless exhibit features suggestive of the presence of a mutant-like p53 protein. To that end, we used the METABRIC BC dataset, in which both the transcriptome (Curtis *et al*., 2012; Rueda, 2019) and the p53 mutation status (Pereira *et al*., 2016; Silwal-Pandit *et al*., 2014) are described for a large number of samples. Using machine learning, we constructed a classifier that best distinguished wt p53 tumors from those carrying *bone fide TP53* mutations. Subsequently, we searched for tumors that were identified by our classifier as mutant, despite harboring wt *TP53*. We found among ER+Her2- BCs a subgroup of PM p53 tumors, with a transcriptome resembling that of true mutant p53 tumors. Moreover, like ER+Her2- BC with authentic p53 mutations, these PM tumors exhibited an enhanced proliferation signature and were associated with worse prognosis. Understanding the mechanisms that underlie the aberrant behavior of p53 in such tumors may provide clues towards individualized treatment, with improved patient outcome.

## 2 Methods

### 2.1 Aims and strategy. Aims

To identify PM tumor samples, and to characterize these tumors and patients. *Strategy:* Using Machine Learning (ML) on single probe expression data, we constructed a robust conservative classifier that assigns each sample a (learned) label of either wt or mutant (mut) p53. False Positive (FP) samples, which are labeled as mutant even though they are wt, are designated as PM.

### 2.2 Data

Probe-level expression data from the METABRIC dataset (Curtis *et al*., 2012) were filtered and preprocessed for 1928 tumor samples for which we had both expression and *TP53* status (see Appendix S1). The final expression table, of 34,363 (HT-12 v3 platform, Illumina_Human_WG-v3) probes representing 24,369 genes, for 1928 tumor and 144 normal samples, can be found in Supplemental file 1 (provided upon request). “Ground truth” *TP53* mutation status was from sequencing: from NGS when available (Pereira *et al*., 2016), or from Sanger Sequencing (SS) (Silwal-Pandit *et al*., 2014). For the learning process, samples without *TP53* mutations and with synonymous mutations were labeled wt, and everything else was labeled mut. In some analyses we separated missense from all other mutations (the latter are referred to as “null”). Samples that were suspected to harbor germline mutations (n=15) were excluded from the analysis (Table S1). Our *TP53* labels, the mutation types and breast cancer subtypes (ER+Her2-, ER- Her2- and Her2+, as determined by immunohistochemistry), are summarized in Table 1 and given in detail in Supplemental file 2, which contains also protein change, survival and treatment information and clinical parameters. Learning, classification, validation and subsequent analyses were done separately for each clinical subtype.

**Table 1.** Number of samples of each clinical breast cancer subtype and p53 mutation status.

| **Subtype\p53 status** | **p53 wt** | **p53 missense** | **p53 null** | **total** |
| --- | --- | --- | --- | --- |
| **ER+Her2-** | 1122 | 153 | 98 | 1373 |
| **ER-Her2-** | 63 | 144 | 108 | 315 |
| **Her2+** | 78 | 95 | 67 | 240 |
| **total** | 1263 | 392 | 273 | 1928 |

METABRIC data used for wt/mut p53 classification: see text for definitions of labels.

#### Focus on the ER+Her2- subtype

Since each BC subtype is a distinct disease with different molecular characteristics, we analyzed tumors from these three subtypes separately and independently. The relative abundance of *TP53* mutations varies greatly between these subtypes, which biases the analysis of the transcriptional effects of such mutations when all subtypes are combined together into a single group (Miller *et al*., 2005). We therefore chose to focus on ER+Her2- tumors: this group comprises by far the largest number of samples of the same subtype, ensuring best statistics. ER+Her2- tumors exhibit pronounced misbalance towards wt *TP53* (1122 wt and 251 mutant samples), whereas the Her2+ and ER-Her2- groups exhibit the opposite misbalance; this also motivated us to focus on the ER+Her2- subtype. Analyses that try to compensate the learning algorithm for imbalanced training sets introduce several uncontrolled parameters. To avoid this, we used learning processes that do not correct for imbalance. In the ER+Her2- group, lack of compensation for the prevalence of wt samples gives rise to a conservative classifier, with a bias towards a wt call. Hence, we have high confidence in the classification of those wt *TP53* samples whose transcriptome deviated so strongly from wt behavior, that a mut call was generated; confidence in identification of such “false positives” is essential for fulfilling the aim of this analysis. With this said, we also ran learning processes that did compensate for sample misbalance (see Appendix S2).

### 2.3 Constructing the wt/mut p53 classifier

The computational pipeline used to generate an optimal classifier/predictor of p53 status on the basis of expression data is presented in Fig. 1.

**Figure 1.**
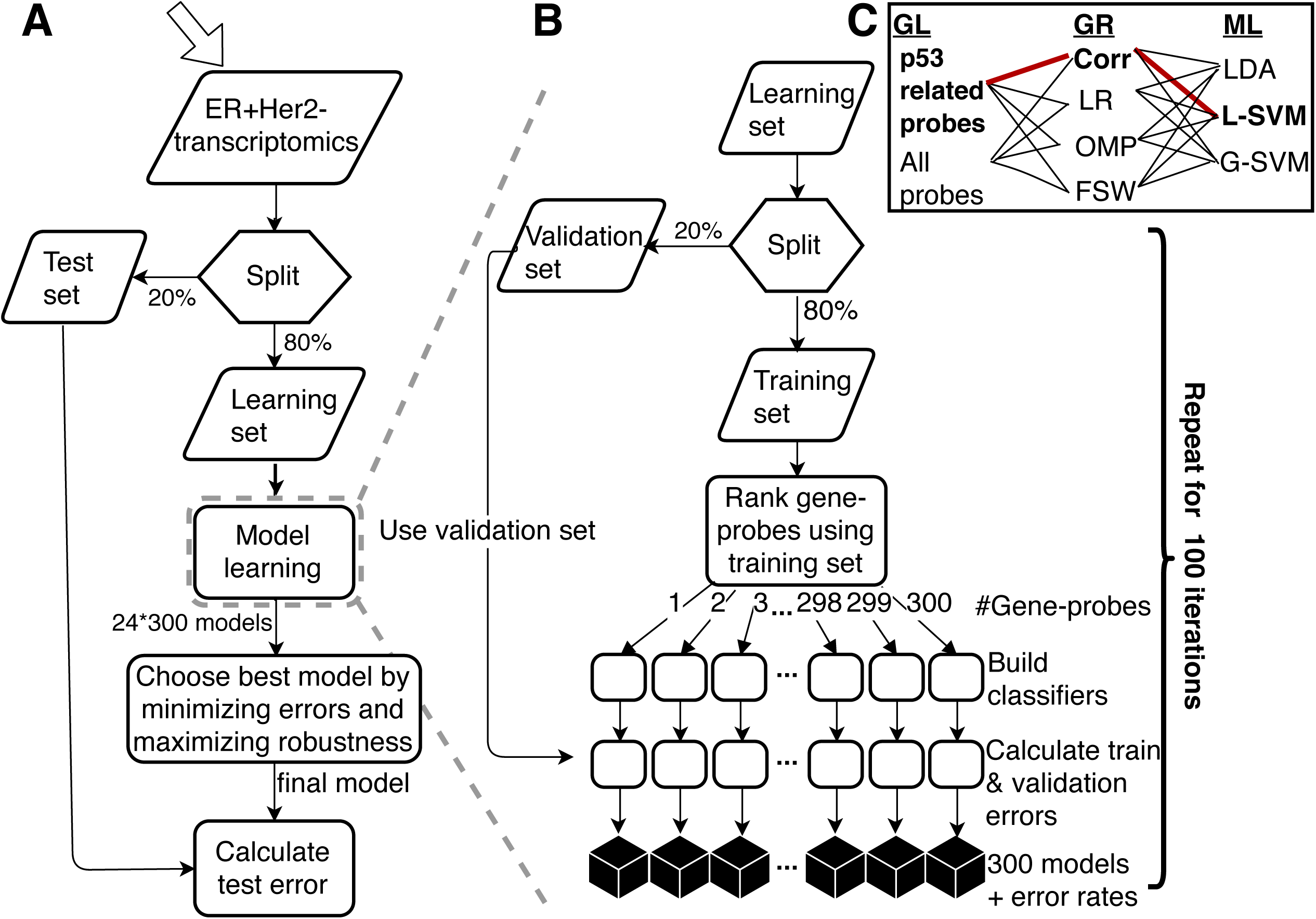
A. Flow chart of the learning process to create a classifier of wt versus mutant *TP53* samples. B. detailed description of the central learning module: for each one of the 100 iterations we ran the learning process for every combination of C. GL (gene-probe list), GR (gene-probe ranking) and ML (machine learning method). Each iteration uses a different learning set and corresponding ranked gene-probe list, generating a sub-classifier of k=1,2,…N≤ 300 genes. Optimal GL*, GR*, ML* and k* are selected on the basis of the validation error and robustness. The final model consists of the majority vote of C (70≤C≤100) sub-classifiers based on k* genes and the selected GL*, GR* and ML* methods. See text for detailed description.

*Step 1* is a 20-80% random split of the samples (separately for the METABRIC Discovery and Validation sets (Curtis *et al*., 2012)) to test and learning sets. The test set is used (only once!) to test the final classifier. The learning set is randomly split to validation (20%) and training (80%) sets. This is repeated 100 times (each split is referred to as an “iteration”).

S*tep* 2 is *feature selection*: since the number of samples available for learning is about one thousand, the number of variables (probes) used to fit this data must be reduced drastically to avoid overfitting. We started from two Gene-probe Lists (GL): (i) knowledge-based, of p53-related genes (Allen *et al*., 2014; Schaefer *et al*., 2009) (425 probes, corresponding to 272 genes) and (ii) using all 34,363 probes that correspond to 24,369 annotated genes. Next, four gene-probe ranking (GR) methods were used to create a ranked gene-probe list (see Fig 1B): Linear Regression (LR) (Mining, 2009), Correlation (Corr) – ranking by the absolute value of the Pearson correlation coefficient between the gene-probes’ expression and the *TP53* status of the learning samples; Forward Selection Wrapper (FSW) (John *et al*., 1994) and Orthogonal Matching Pursuit (OMP) (Pati *et al*., n.d.) (see Appendix S3). Only those *N* gene-probes that passed Benjamini-Hochberg FDR (Benjamini and Hochberg, 1995) at q=0.05 for the property used for ranking (e.g. correlation of expression with *TP53* status) were retained. For each iteration and each GL/GR combination, we generated a list of the top-ranked *N* ≤ *300* probes. Ranking features is a notoriously unstable process – two slightly different training sets may produce very different lists of top-ranked genes (Ein-Dor *et al*., 2005, 2006). Repeating the process 100 times stabilizes the results of feature selection by ranking; robustness was tested according to several criteria (see Appendix S4, Fig. S1 and Table S2).

*Step 3 is learning: for each iteration* and its associated ranked gene-probe lists we learned the p53 labels, using three machine learning (ML) methods: Linear Discriminant Analysis (LDA) (Duda *et al*., 2001), Linear Support Vector Machine (LSVM) and Gaussian Support Vector Machine (GSVM) (Mining, 2009).

*Step 4 - Model selection*: For every iteration, each learning process (GL, GR, ML combination) was applied using the top ranked *k=1,2,3…N* probes, adding one probe at a time. We call a classifier “valid” if all its *k* probes passed the threshold of FDR = 0.05 (e.g. p-value for correlation). For each *k* we required to have at least *C=70* valid classifiers, for which the classification error was calculated for the training set and for the validation set, otherwise we stop at *k-1*. Finally, the training and validation errors were averaged over the valid classifiers, and plotted as a function of the number of probes used. These plots, together with considerations of robustness, were then used to determine the optimal learning process (GL*, GR*, ML*) and *k\**, the number of probes used, to yield the optimal set of sub-classifiers.

Notably, the total number of classifiers that was constructed is on the order of 2×4×3×300×100 = 720,000 (in practice it is smaller; e.g. since the number of valid classifiers may be less than 100, the number of probes that passed various filters may be less than 300, and not all GL/GR/ML combinations were tested).

*Step 5: The final master classifier* uses, for a given sample, the expression levels of the *k\** probes of each of the *70*≤*C*≤*100* sub-classifiers to produce *C* predicted p53 labels; the majority vote of these constitutes the final deduced p53 status.

*Step 6: testing the master classifier*. The samples of the test set were submitted to the master classifier and the test error was calculated. Tumors identified as mut are referred to as “Positive”; the error is the ratio of (False Positives + False Negatives)/(Total number of classified samples).

*Step 7: identification of the PM samples:* False Positive samples from *the test set* were labeled PM. For *the learning set* we relied only on misclassification of the validation samples, using the following rule: identify the iterations in which a wt sample was assigned to the validation set; if in the majority of these iterations it was classified as mut – it is labeled PM.

### 2.4 Validating the classifier on an independent dataset – TCGA

The classifier derived on the basis of METABRIC data was tested on TCGA breast cancer data (Koboldt *et al*., 2012). The TCGA dataset contains 98 mutant and 381 wt *TP53* ER+Her2- samples, for which expression was measured using RNA-seq. The manner in which the TCGA data was handled to account for different representations of genes in TCGA and METABRIC is described in Appendix S5.

### 2.5 Dimensionality reduction

To demonstrate intermixing of the PM group with mut samples, Principal Component Analysis and tSNE (Van Der Maaten and Hinton, 2008) were used.

### 2.6. Characterizing the PM samples; Pathway-level analysis

The METABRIC samples were studied also on pathway level, using Pathifier (Drier *et al*., 2013; Livshits *et al*., 2015). Lists of genes that belong to various pathways were downloaded from KEGG (Ogata *et al*., 1999), BioCarta (Nishimura, 2001) and the NCI-Nature curated Pathway Interaction Database (Schaefer *et al*., 2009). All probes that appeared among the top 5000 varying ones (Drier *et al*., 2013) were used in the analysis; hence some genes were represented by more than a single probe. For each pathway *P* and sample *k* we derived *D(P,k)*, a Pathway Deregulation Score (PDS), as described (Drier *et al*., 2013). The samples and pathways were sorted using SPIN (Tsafrir *et al*., 2005), to place together groups of similarly deregulated pathways and samples with similar deregulation profiles. Deregulation of pathways in the PM samples was compared to the mutants and to the wt (by sequence *and* classification), by two-sided t-tests and Benjamini-Hochberg FDR correction for multiple hypotheses. All the genes belonging to the pathways that were significantly differentially deregulated between the PM group and mut p53 samples were collected. For each corresponding gene-probe expression we calculated a two-sided t-test and FDR correction between the PM and the mut p53 groups.

### 2.7 Clinical characterization

Kaplan-Meier analysis of survival data of the three tumor types was performed by Matlab routine “ecdf” for calculating the curve and “survdiff” for calculating the p-value of log-rank test.

## 3 Results

### 3.1 The Training Process

We generated classifiers that distinguish wt *TP53* tumors from mut *TP53* tumors, based on their gene expression profiles. We present here results only for ER+Her2- tumors; for the other subtypes see Appendix S6, Table S3 and Fig. S2. We assumed that if PM tumors indeed exist, they probably constitute a minority of the wt *TP53* cases, whereas most wt *TP53* cases retain a transcriptional profile distinct from that of true mut *TP53* tumors. For all tested combinations of gene lists (GL), gene ranking (GR) and machine learning (ML), 100 iterations were performed. Classifiers based on *k* gene probes were constructed by adding one probe at a time, according to their ranking. Note that for ER+Her2- we stopped at *k=138* probes; beyond this, the number of valid classifiers was less than 70. For each *k*, GL, GR and ML combination, the average and standard deviation of the error (training and validation) over the 100 iterations was stored. These error curves, together with considerations of their robustness (Appendix S4), served to select the optimal classification model (see Appendix S7 and Fig. S3). The selected model for ER+Her2- tumors is presented in Table 2; it uses the p53-related gene list as GL*, gene probes ranked by correlation to p53 status as GR*, and LSVM as the ML* method of choice. The fact that the p53-related gene list separates well wt *TP53* tumors from those that carry *TP53* mutations supports our assumption that the majority of the former tumors retain a wt p53-driven transcriptional program.

**Table 2.** Details of the final classifier for ER+Her2- tumors and the classification errors for METABRIC and TCGA data (see text).

| #Training samples | #Validation samples | #Test samples | #probes k*<br>(mean ±SEM genes) | Selected GL*<br>(GR*) | Selected ML*<br>method | Validation error mean ±SEM | Training error mean ±SEM | Test error (majority vote) | TCGA test error (majority vote) |
| --- | --- | --- | --- | --- | --- | --- | --- | --- | --- |
| 878 | 220 | 275 | 62 (49.3±0.01) | p53related genes (Corr) | Linear SVM | 0.116±0.002 | 0.0868 ±0.006 | 0.13 | 0.11 |

The mean errors (of 70≤*C*≤100 classifiers) plotted as a function of gene probe number *k*, and the mean Receiver Operating Characteristic (ROC) curves of the optimal classifiers, are presented in Fig 2A and Fig 2B, respectively. We decided to use *k*=62* probes, representing, on average, 49.3±0.1 genes (mean ±SEM); this yielded a mean±SEM validation error of 0.116±0.002 (Table 2). Shorter gene lists are more robust, and at 62 probes we had 100 valid classifiers. Hence, we preferred *k*=62*, even though the validation error was slightly higher than for 125 probes (Fig. 2A). The list of 62 probes used for each of the final 100 sub-classifiers is in Supplemental File 3. The ROC curve presented is the average (over all the 100 classifiers) of the ROC curves that were obtained separately by Matlab routine “perfcurve” for each classifier.

**Figure 2.**
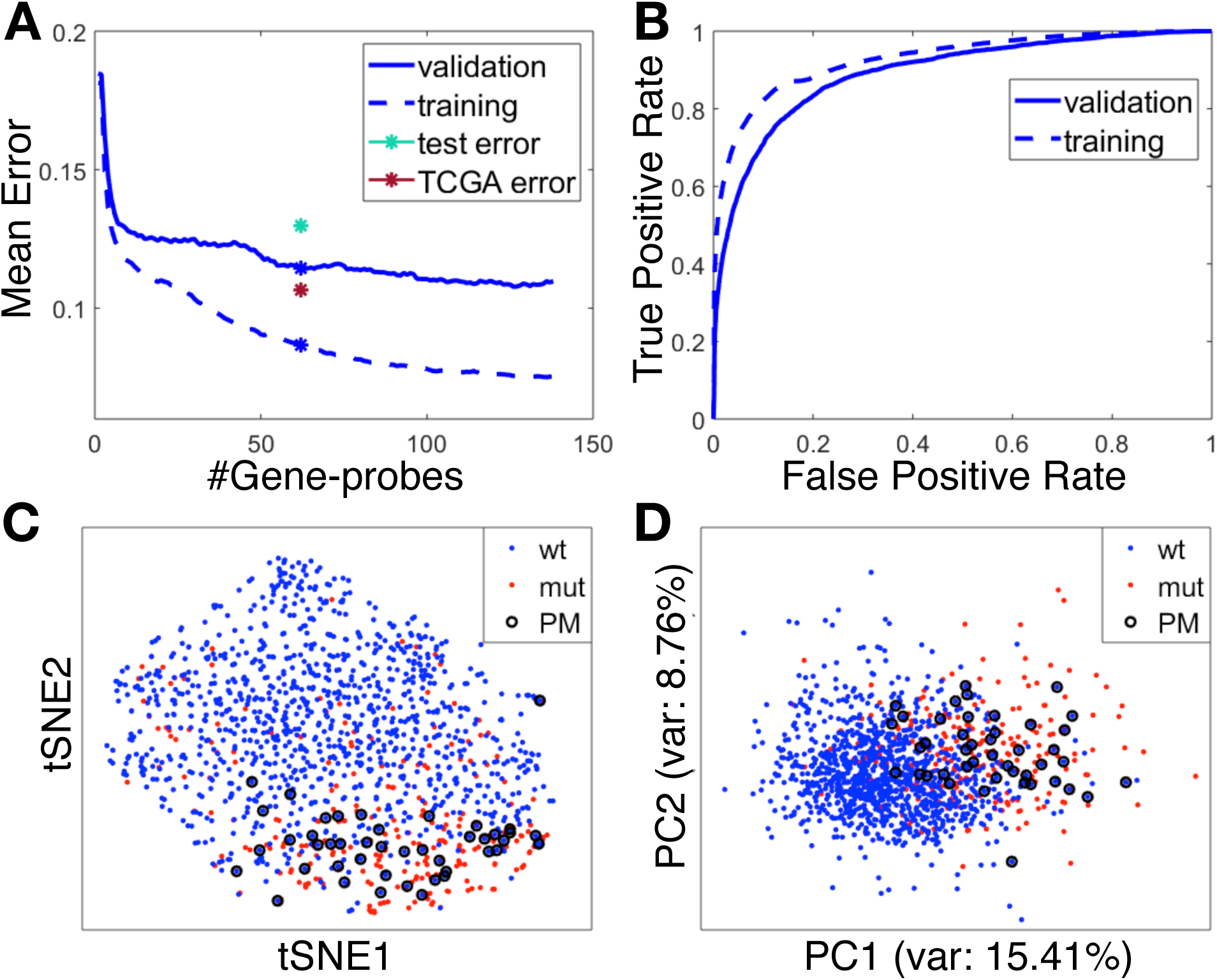
Accuracy and error rates for the chosen model for ER+Her2- tumors. A. Mean error rates of prediction of *TP53* status (mut/wt) for the validation and training sets, using the chosen ML, GR and GL, for 100 iterations, versus the number of top ranked gene probes used. The test (METABRIC) and the TCGA errors of the final model (with 62 gene probes) are marked by green and red asterisks, respectively. The mean validation and training errors of the selected model are marked by blue asterisks. B. Mean ROC Curves for the final model for both validation and training sets. The Area Under the Curve values for the validation curve and the training curve are 0.89 and 0.93, respectively. C. tSNE dimensional reduction from the space of the 62 most used gene probes (see text) to 2 dimensions. Positions of wt (blue), mut (red) and PM (circled) show that the PM tumors are intermixed with the mut tumors. D. Same as C, using PCA.

### 3.2 Identifying “pseudomutant” p53 tumors

The final wt/mut p53 call, for a previously unseen sample (e.g. from the test set), is obtained as a majority vote of the 100 sub-classifiers (Appendix S8, Table S4 and Fig. S4). The test error of this final classifier was 0.13 (Table 2): out of 224 wt *TP53* samples, 14 were misclassified as mutant, and were defined as PM. Of note, since we decided not to correct for the unbalanced learning data (see Methods), our classifier is strongly biased towards wt calls and against mutant calls; as a result, out of the 51 mutant *TP53* test samples, 22 were falsely classified as wt (false negative, FN). When the imbalance is corrected, this error rate is reduced to 12/51. We then returned to the training set, to identify additional PM tumors (see Methods, Step 7). Altogether, we found 30 such additional cases (out of 898 wt *TP53* training samples), so that in total 44 out of 1122 (about 4%) wt *TP53* ER+Her2- tumors were identified as PM (for patient IDs see Table S5). This fraction of 4% is very robust: a similar percentage was obtained independently of the GL, GR and ML methods used. Of note, our classifier is conservative, with a bias against mutant p53 calls; hence, the frequency of PM cases may actually be higher. Indeed, a larger number of samples (19 from the test set and 76 from the validation sets, Appendix S2) were identified as PM when we used a learning process (Table S6) that compensates for the unbalanced learning set (Appendix S2), bringing the percentage of PM cases up to about 8.5%.

Next, we ranked the gene probes according to the number of their appearances among the top 62 of the 100 iterations, and used the top ranked 62 probes to represent each sample. Dimensionality reduction to two dimensions (tSNE and PCA) was then applied (Fig. 2C and 2D). Remarkably, the great majority of PM samples projected onto “mutant territory” and were intermixed with authentic mut *TP53* tumors, further displaying the similarity between the transcriptional profiles of the two groups.

It is important to rule out the possibility that misclassification of wt as mut may have been due to questionable sequencing data. In that regard, we note that 34 of the 44 PM tumors were sequenced by *both* Sanger sequencing (SS) and NGS. Of those, 30 were called as wt *TP53* by both methods. Only for 4 was there a discrepancy; they were called wt by NGS but mutant by SS. Of the remainder 10 wt *TP53* cases, 8 were sequenced only by NGS and 2 only by SS (Table S7). The probability of mistakenly identifying a sample as wt by two independent methods is very low, giving good reason to trust that the PM tumors are not mis-sequenced *TP53* mutants. Deeper curation of the 4 discordant tumors indicated that 3 of these (one from the test set and two from the learning set, see Appendix S9) were complex frameshifts, seen in SS chromatograms but missed by the NGS indel mutation caller. A fourth case had a single indel read in NGS. Importantly, the potential misclassification of this very small number of tumors does not affect our conclusions.

In all but one of the PM cases, mutations were identified in other sequenced genes (Supplemental File 4). This strongly suggests that the cellularity and sequencing quality of those samples should have been sufficient to detect *TP53* coding region mutations, had they existed in those samples.

### 3.3 “Pseudomutants” in TCGA data

The TCGA ER+Her2- BC dataset contains 98 mutant and 381 wt *TP53* samples. Gene expression was measured by RNA-Seq, and some of the METABRIC genes we used for classification had no reported expression levels in TCGA. Consequently, only 42 out of the 100 sub-classifiers had TCGA expression data for all their genes, and the majority vote of these 42 sub-classifiers was used for wt/mut *TP53* calling. Nevertheless, the TCGA error rate of 0.11 is even lower than that of the METABRIC test error. Through this analysis, 15 PM samples were identified in TCGA, which again constitutes about 4% of all wt *TP53* samples. Thus, our classifier works for a different set of patients, with expression measured by a different platform; this constitutes strong evidence for the robustness of our method. The similar percentage of PM cases indicates that their identification is not an artifact of one particular dataset.

### 3.4 Pathway deregulation in the PM tumors

Moving from single gene-based transcriptomes to pathway level, we combined the expression profiles the 1373 ER+Her2- tumors with those of 144 normal breast samples and calculated the pathway deregulation score (PDS) (Drier *et al*., 2013; Livshits *et al*., 2015) of each sample for 379 pathways that passed a threshold of stability. The heatmap of these scores is presented in Fig 3A. Both samples and pathways are ordered in a manner that places in proximity objects with similar PDS profiles (Tsafrir *et al*., 2005). Again, most PM samples were assigned to a region preferentially occupied by mut *TP53* tumors, indicating that their pathway deregulation profiles resemble those of these tumors.

**Figure 3.**
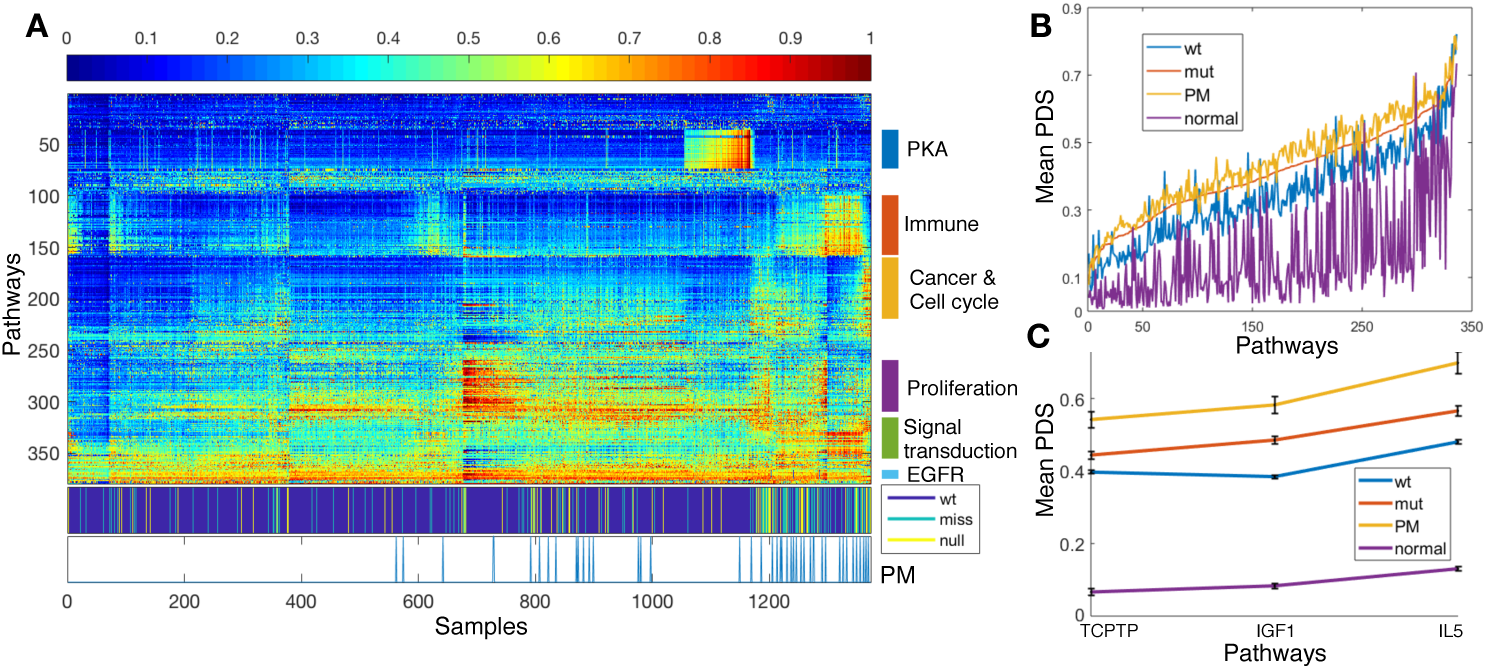
Pathway deregulation-based analysis of ER+Her2- tumors. A. Heatmap of the pathway deregulation scores (PDS) of 1373 tumors and 144 normal breast samples, encompassing 379 pathways. Samples and pathways were ordered by SPIN, placing in proximity objects of similar deregulation profiles. The second color bar from the bottom identifies *TP53* mutation status: wt, missense and null. The bar below it identifies the PM samples; remarkably, these samples cluster in the region dominated by mutants. Different pathway clusters are identified on the right side. B. Mean PDS of 336 pathways that were differentially deregulated in PM versus wt p53 tumors. Pathways were ordered by their mean PDS in the mutant samples. C. Mean PDS of normal breast tissue and of the wt p53, mut p53 and PM groups, for three pathways that are differentially deregulated between PM and mut p53 tumors.

Next, we looked for pathways displaying significantly different mean PDS in PM versus the rest of the wt *TP53* tumors. Out of 574 tested pathways, 336 passed as differentially deregulated in PM versus wt p53 at an FDR of 0.05. Ordering these pathways by the mean PDS of the mut *TP53* samples, we plotted the mean PDS of the PM, mut and wt p53 tumors and of normal mammary tissue samples (Fig 3B). Interestingly, the pathway deregulation scores of the wt p53 tumors were closest to normal tissue, while those of the mut p53 cases were higher. Remarkably, the PM tumors had the highest mean pathway deregulation: nearly all 336 pathways had a higher mean PDS for the PM than for the mut *TP53* cases, indicating an extremely significant overall deregulation difference (on a multi-pathway level). Thus, at least by PDS assessment, the PM tumors appear to behave as even “more mutant” than true mutants.

Deregulation of cancer-related pathways can cause wt p53 to adopt mutant-like behavior (Furth *et al*., 2015). We therefore looked for pathways that are preferentially deregulated in the PM samples, relative to mut p53 breast tumors, and therefore may potentially drive wt p53 protein into a PM state. Using an FDR threshold of 0.1, we found three such pathways (Fig. 3C): the IGF1 (BIOCARTA), IL5 and TCPTP pathways (NCI). Altogether, these 3 pathways comprise 67 genes. When each of the corresponding gene-probes was tested individually for differential expression between the PM and mut p53 groups, only two probes, both representing *GRB2*, were significantly differentially expressed. GRB2 (Growth Factor Receptor Bound Protein 2) is an adaptor protein that links growth factor receptors to the Ras signaling pathway. As seen in Fig 4, its mean expression, in both METABRIC and TCGA, was indeed higher in the PM tumors than in the mut or wt p53 tumors.

**Figure 4.**
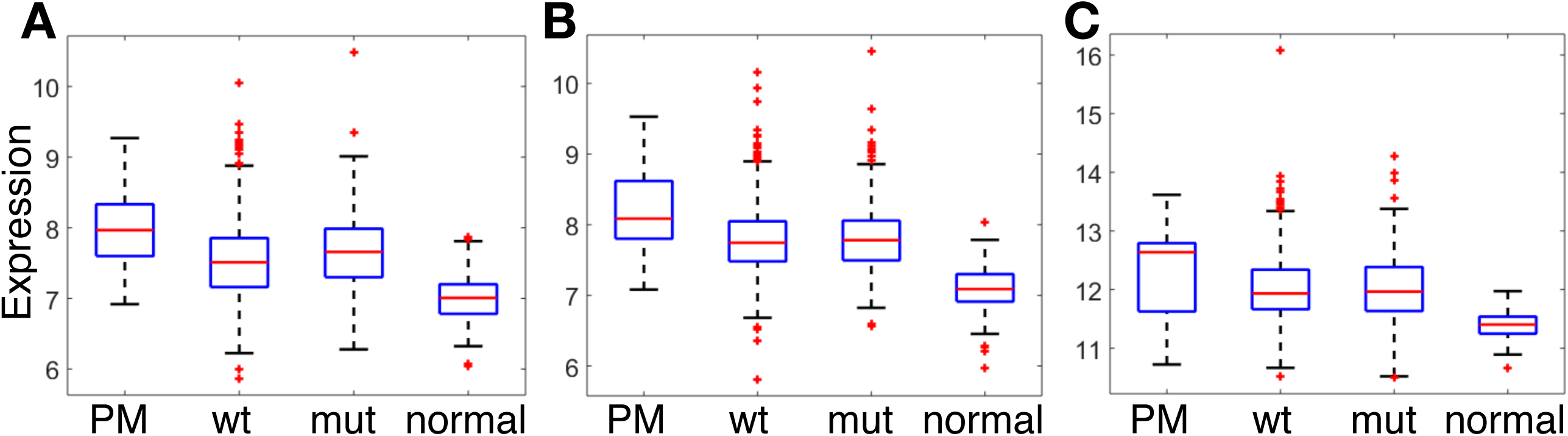
Mean (log) expression values of GRB2 for the different groups of samples. A. METABRIC, probe ILMN 1742597, PM vs mut; p = 0.00035; B. METABRIC, probe ILMN 1748721, PM vs mut; p = 0.00003; C. TCGA, PM (n=15 samples) vs mut; p=0.11.

### 3.5 Clinical features of the PM tumors

MKI67 mRNA levels, indicative of cell proliferation, were significantly higher (p=1.5×10^−5^) in the PM group than in the wt p53 group, resembling those of mut p53 samples (Fig. 5A). Hence, PM tumors possess higher average proliferation rates than the remainder of the wt *TP53* tumors (Fig. 5A). Moreover, the prognosis of BC patients with ER+Her2- PM tumors is significantly worse than that of patients with “conventional” wt p53 tumors, and similar to that of patients with authentic mut *TP53* tumors (Fig. 5B). Hence, PM tumors resemble *bona fide* mut *TP53* tumors not only in gene expression and pathway deregulation patterns, but also in clinical features.

**Figure 5.**
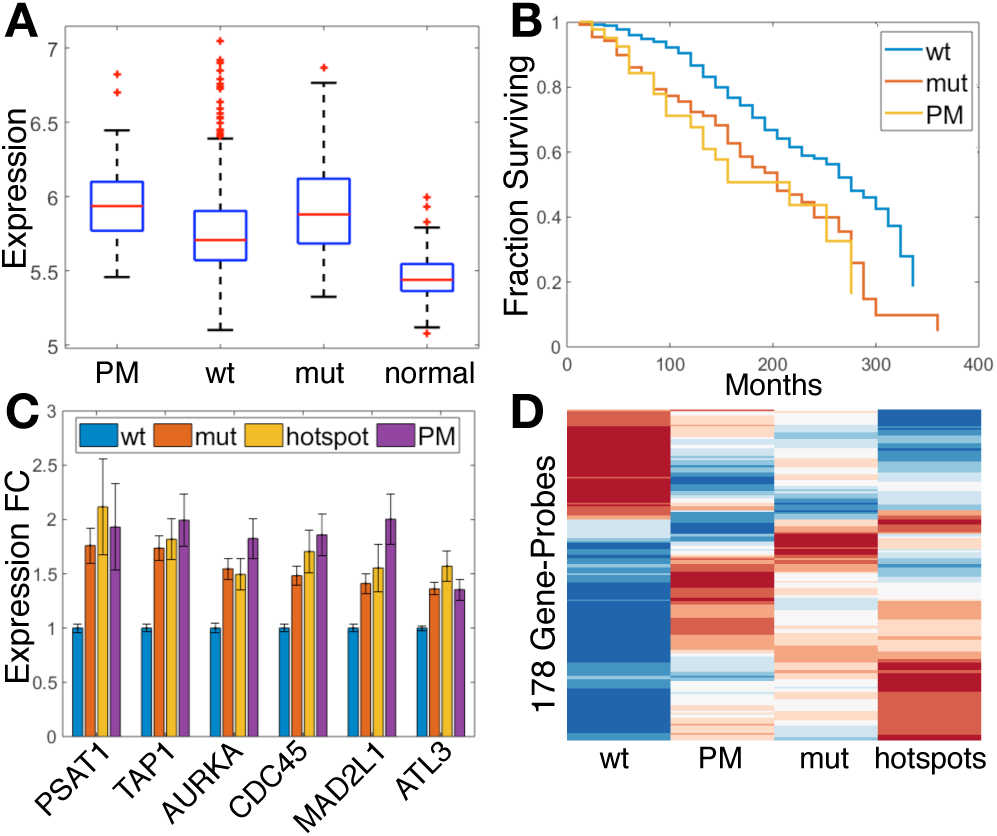
A. Expression levels of MKI67 in PM, wt, and mut p53 ER+Her2- breast tumors and in normal breast tissue samples. B. Kaplan-Meier survival curves for BC patients bearing PM, wt or mut p53 ER+Her2- tumors, with p-value = 0. 00276, calculated by log-rank test, between the survival of PM vs. wt. C. Expression fold change values in the indicated tumor groups, relative to wt p53 tumors, of six mutant p53- regulated genes. “hotspots” indicates tumors harboring one of the 10 most common *TP53* hotspot mutations, according to the IARC mutation database (Hainaut *et al*., 1998). D. Mean expression of 178 gene-probes corresponding to mutant p53-regulated genes (Walerych et al., 2016) that passed FDR<0.05 between wt and mut p53, displayed for the wt, PM, mut and p53 hotspots groups of samples. The gene-probes were ordered by hierarchical clustering, using the “clustergram” Matlab function.

Lastly, since common p53 missense mutants can exert oncogenic GOF effects, we asked whether PM p53 may also possess GOF activities in BC. To demonstrate p53 GOF formally, one needs to show that depletion of the tested p53 protein attenuates cancer-related features. Obviously, this is impractical for archived tumor specimens. Therefore, we took advantage of a study of *TP53*-mutated BC cell lines (Walerych *et al*., 2016). Silencing the endogenous mut p53, followed by RNA-Seq analysis, identified a core group of 205 genes that are specifically regulated by mut p53 and contribute to its GOF in BC (Walerych *et al*., 2016). These 205 genes are represented in METABRIC by 337 probes; 178 of these probes separate wt from mut *TP53* cases at FDR<0.05 (see Fig. 5D for heatmap of their expression for all samples), while 104 separate PM from the other wt cases (with 88 probes shared between these two lists). In Fig. 5C we display results for the six genes with the highest fold change between the mut p53 and the wt p53 ER+Her2- samples in the METABRIC dataset. As expected, all 6 genes were expressed more abundantly in the mut p53 tumors compared to wt tumors; in most cases, their expression was slightly higher in tumors with hotspot p53 mutations (10 most frequent *TP53* mutations in IARC mutation database (Hainaut *et al*., 1998)), consistent with their proposed contribution to mut p53 GOF. Notably, expression of those genes in the PM tumors was at least as high as in hotspot mut p53 tumors (Fig. 5C). This raises the intriguing possibility that in tumors where wt p53 acquires PM features, it not only loses its tumor suppressive activities but may even gain cancer-promoting activities.

## 4 Discussion

Breast cancer is a heterogeneous disease, with diverse subtypes, each driven by distinct molecular and genetic mechanisms. We developed classifiers, separately for each breast cancer subtype, which differentiate well between tumors that retain wt p53 expression and those that have undergone *TP53* mutations. To avoid the confounding effect of the marked differences in percentages of mut p53 in the different subtypes, each subtype must be analyzed as a separate group. The large number of samples in the METABRIC study, with available expression data (Curtis *et al*., 2012), *TP53* sequencing (Pereira *et al*., 2016; Silwal-Pandit *et al*., 2014) and clinical information enables statistical robustness of subtype-specific analysis. For the largest group, of ER+Her2- patients, even though 275 tumors were set aside as a test set, the number of tumors that remained available to be used for learning was large enough for reliable training and validation. By combining 100 repeats of the entire training process, each with its own random training/validation split, we overcame the lack of stability of ranked gene lists that has plagued most single-gene based classifiers (Ein-Dor *et al*., 2005). We derived a very robust expression-based classifier, which separated successfully mutant from wt *TP53* tumors also in the independent TCGA cohort of patients (Koboldt *et al*., 2012).

Using this approach, we identified a group of ER+Her2- patients whose tumors harbor wt *TP53* but display a mutant p53-associated transcriptional program. These “pseudomutant p53” tumors resemble authentic *TP53*-mutated ER+Her2- breast tumors not only in their gene expression and pathway deregulation profiles, but also in their highly proliferative nature and shorter patient survival.

What may cause a wt *TP53* tumor to acquire a PM transcriptional profile? Potentially, such an outcome might be obtained by complete loss of p53 expression, rendering these tumors practically p53-null (Miller *et al*., 2005). However, although we saw a slight reduction in p53 mRNA levels in our PM group relative to the rest of the wt p53 samples (Fig. S5), p53 expression was significantly higher in the PM tumors than in tumors carrying nonsense/frameshift mutations (p-value = 0.0022, two-sided t-test). Thus, additional mechanisms are likely to contribute to the PM behavior. These might involve deregulation of cancer-relevant signaling pathways; for example, it was previously shown that deregulation of the Hippo pathway can alter the functionality of wt p53 (Furth *et al*., 2015). Indeed, pathway-level comparisons (Drier *et al*., 2013) demonstrated that hundreds of biological pathways are significantly more deregulated in PM than in wt p53 tumors. Interestingly, elevated expression of *GRB2* was particularly symptomatic of PM tumors. GRB2 is involved in relaying signaling downstream to numerous growth factor receptors. Hence, although causality between high GRB2 and PM p53 behavior remains to be proven, it resonates well with the early observation that wt p53 protein can be shifted into a mutant-like conformation upon growth factor stimulation (Zhang and Deisseroth, 1994). Additionally, epigenetic changes may also contribute to acquisition of a PM p53 phenotype, as suggested by the mutant-like behavior of wt p53 in cancer-associated fibroblasts (Arandkar *et al*., 2018).

Elucidation of the molecular mechanisms that drive wt p53 into a pseudomutant state may identify potential treatments that can restore wt p53 functionality in tumors harboring PM p53. This may selectively benefit patients whose tumors display PM features in association with bad prognosis. Thus, further elaboration of the underlying mechanisms is highly desirable.

## Acknowledgements

This work was supported in part by grants from the Dr. Miriam and Sheldon G. Adelson Medical Research Foundation, the Rising Tide Foundation, and the DKFZ-MOST cooperation in cancer research.

## Author contributions

Author contributions: G.B., E.D. and M.O. designed research; G.B. performed research; G.F. provided computational advice, S-F.C, O.R. and C.C. provided insights about the data, S.M. and S.A. performed RNA quantification, E.D. and M.O. conceived the study and provided computational and biological guidance, respectively. G.B., E.D. and M.O. wrote the paper.

## Consent to publish

The authors are fully responsible for the contents of this manuscript, and the views and opinions described in the publication reflect solely those of the authors.

## Data availability

All data is available in supplementary files.

