## Supporting Information for "Transcriptional profiling reveals a subset of human breast tumors that retain wt *TP53* but display mutant p53-associated features"

**This PDF file includes:**

Supplementary text

Figs. S1 to S5

Tables S1 to S7

References for supplementary information

**Appendix S1. Pre-processing**

The METABRIC dataset consists of 2000 primary fresh frozen breast cancer samples, with 144 additional samples of matched normal tissues. Each sample has undergone transcriptional profiling by Illumina Human v3 microarray platform. The resultant gene expression data is at probe level, log scaled and normalized (1) (clinical information and TP53 mutation data were downloaded from the cBioPortal (2). TP53 mutation status was generated by Next Generation Sequencing with average sequencing depth of at least 112X in 80% of samples. Subtype information (ER+Her2-, ER+Her2- & Her2+) was generated by the METABRIC ER and Her2 IHC, which was also confirmed by gene expression data of ESR1 (for determining ER status) and ERBB2 (for determining Her2 status) genes.

Out of the samples for which transcriptomics and TP53 sequencing were available, 1904 had also TP53 sequencing by NGS from 2016 (3), and 39 additional samples had TP53 Sanger sequencing from 2014 (4). 15 samples with germline mutations were excluded from the analysis (see Table S7), owing to the fact that germline mutations are distinct from somatic mutations in their effects and clinical implications (5). In total, there were 1928 samples for which all the relevant information required for performing our analysis (expression, TP53 status & subtype) was available.

Associating probes with genes was done according to the METABRIC pipeline. Only probes representing known genes were converted into gene level, retaining for each gene all its corresponding probes (a gene with 5 different probes is represented by all 5 probes). Genes with bad quality annotation (according to illuminaHumanv3 information in R) were excluded from the analysis, similarly to the METABRIC pipeline. Each probe was standardized in log scale expression by subtracting its average value within each subtype (ER+Her2-, ER-Her2- and Her2+) and then dividing by the gene’s std in the subtype.

**Appendix S2. Imbalanced Learning**

Since the number of wt p53 and mut p53 samples in each of the subtypes is biased, the same learning process (Fig. 1) was repeated using a corrected score function for imbalanced data. For this aim, we updated the penalty of the cost function of the “fitcsvm” Matlab function as follows:

The penalty for classifying a sample from the majority class (e.g. wt p53 in ER+Her2-) as a minority (e.g. mutant p53 in ER+Her2-) for a specific iteration i was equal to $\frac{\#minority \mathrm{samples}_{i}}{\#majority \mathrm{samples}_{i}}$ , where the number of samples from each class is counted according to the randomly sampled training set for the current iteration i.

The penalty for classifying a minority class sample to the majority class was equal to 1.

A correct classification gets no penalty (0.0). We repeated the final constellation of the chosen models (p53 related genes for ER+Her2-). We also constructed a similar classifier which was based on all gene probes, but similarly to the classifiers without bias correction, it had a higher error rate than the original one based on p53-related genes. Correlation was used as the chosen Gene Ranking method, similar to the chosen GR of the original classifier. The validation errors of the imbalanced-corrected model resembled the original validation errors (0.127 for ER+Her2-), with Gaussian SVM having a lower error rate than linear SVM (See Table S5).

Altogether, 95 ER+Her2- wt p53 samples were classified as mut p53, according to the majority vote of the sub-classifiers; 76 during the learning phase (based on the number of iterations in which each of the samples appeared in the validation set) and 19 during the test phase (majority vote of all 100 sub-classifiers).

The IDs of these samples are displayed below.

76 wt-p53 samples from the learning phase had mut p53 classification according to the majority vote of the 100 sub-classifiers (with the corrected score function for imbalanced data) based on the validation sets:

'MB-0143' 'MB-0147' 'MB-0173' 'MB-0192' 'MB-0197' 'MB-0305' 'MB-0311' 'MB-0321' 'MB-0383' 'MB-0392' 'MB-0393' 'MB-0408' 'MB-0532' 'MB-0538' 'MB-0541' 'MB-0542' 'MB-0550' 'MB-0598' 'MB-0666' 'MB-2634' 'MB-2686' 'MB-2728' 'MB-2969' 'MB-3006' 'MB-3026' 'MB-3171' 'MB-3275' 'MB-3389' 'MB-3466' 'MB-3506' 'MB-3614' 'MB-3823' 'MB-4171' 'MB-4235' 'MB-4278' 'MB-4342' 'MB-4767' 'MB-4876' 'MB-4981' 'MB-4986' 'MB-4991' 'MB-5001' 'MB-5124' 'MB-5139' 'MB-5177' 'MB-5267' 'MB-5306' 'MB-5311' 'MB-5405' 'MB-5424' 'MB-5447' 'MB-5460' 'MB-5467' 'MB-5491' 'MB-5521' 'MB-5534' 'MB-5552' 'MB-5575' 'MB-5636' 'MB-5653' 'MB-6060' 'MB-6077' 'MB-6101' 'MB-6107' 'MB-6187' 'MB-6208' 'MB-6218' 'MB-6226' 'MB-6287' 'MB-6302' 'MB-6312' 'MB-7018' 'MB-7043' 'MB-7061' 'MB-7092' 'MB-7133'.

19 wt p53 samples from the test phase had mut p53 classification according to the majority vote of the 100 sub- classifiers (with the corrected score function for imbalanced data):

'MB-0066' 'MB-0195' 'MB-0131' 'MB-0283' 'MB-4004' 'MB-4937' 'MB-4148' 'MB-4834' 'MB-5350' 'MB-5366' 'MB-5188' 'MB-3600' 'MB-7011' 'MB-7161' 'MB-6118' 'MB-7299' 'MB-6284' 'MB-7292' 'MB-0543'.

Out of the 44 original PM samples (wt p53 classified as mut p53), 43 were also classified as mutants using the imbalanced corrected procedure (by majority vote of the sub-classifiers) among the 95 updated PM samples. Two sided t-test for each of the two probes of GRB2 between the PM group (discovered by the imbalanced-corrected classifier) and the rest of the wt p53 samples was calculated, with the following two p-values: for ILMN_1742521: $8.88e^{-8}$ and for ILMN_1748797: $5.67e^{-9}$. One can see that even after correcting for the imbalance classes in ER+Her2- the result still holds and GRB2 has a significantly differential expression between the wt p53 and the PM groups.

**Appendix S3. Forward Selection Wrapper and Orthogonal Matching Pursuit**

**A. Forward Selection Wrapper (FSW)** (6, 7)**.**

In this approach, the learning data was first divided randomly into training (80%) and validation (20%). The validation set was set aside and not accessible until the process of building the Forward Selection Wrapper (with SVM as the core ML method) was completed for the current iteration. In each step, the training data was further split into Real Training (RT, 80% of the training data), which was used for ranking the genes and constructing the core ML model (Gaussian or linear SVM) according to the pool of predictive variables (probes) and Mini Val (MV, 20% of the training data) that was used for deducing which of the available variables reduces the validation error the most. At each iteration, the Pearson correlation p-value of each probe with p53 binary mutation status was calculated, and corrected using Benjamini Hochberg FDR for inferring the q-values. Only probes with q-value below some threshold (0.05) were tested for inclusion into the probe set for the FSW. The probe chosen to be added to the model at a given step is that one which, when added to the pool of already selected probes, causes maximal decrease of the loss function. The loss function is the fraction of misclassified samples over the MV set, using the SVM model (that was constructed using the RT set).

$$Loss\left( mdl,g \right)= \frac{1}{N}\sum_{i=1}^{N} C(\hat{y_{i}},y_{i})$$

Where,

${{\hat{y_{i}}=sign[f}_{svm}(x}_{i},g)]$is the classification of $x_{i}$ by the SVM model $mdl$ created with the pool of already chosen probes with the additional probe g.

$N$is the number of samples in the $MV$set.

$C\left( \hat{y},y \right)$is the cost function of assigning a sample $x_{i}$ with true label y to class$\hat{y}.$

$C\left( \hat{y},y \right)=\left\{ \begin{aligned} 1 if \hat{y}\neq y \\ 0 if \hat{y}=y \end{aligned} \right.$.

Probes were added, one at a time, until 100 probes were in the pool of selected variables. This process was performed once using linear SVM as the core ML method and once using Gaussian SVM.

**B. Orthogonal Matching Pursuit** (OMP).

We used the OMP algorithm for finding a sparse representation for the expression of the probes that explains the p53 mutation status of ER+Her2- samples.

The Matching Pursuit (MP) algorithm was invented by Mallat and Zhang (8) and the Orthogonal Matching Pursuit (OMP) is a modification of the MP algorithm which also maintains full backward orthogonality of the residual (error) at each step, a property that improves convergence (9).

At each iteration of the algorithm the feature (probe) that is added to the pool of selected features is the one whose correlation with Ri (the residual) is the highest.

Where,

$Y_{i}$ is the orthogonal projection of the objective Y (the binary mutation status of the samples) onto the subspace spanned by the expression vectors of the already selected probes in the pool until iteration i (there are i-1 such probes).

$Ri=Y-Y_{i}$ is the orthogonal remainder of the objective’s orthogonal projection (un-explained by the previously chosen features) during iteration i. The stop condition was chosen to be 300 probes.

For further information see Mallat & Zhang (8) and Pati, Rezaiifar, & Krishnaprasad (9).

**Appendix S4. Robustness measurements for both genes and lists.**

Two robustness estimators were computed:

A. Lists’ robustness - Mean and std of the intersect of the final chosen number of top ranked genes (e.g. 62 for ER+Her2-) of every pair of lists obtained by the same GR method are presented in Table S1 (there are $\binom{100}{2}$ such pairs).

The ratio of mean intersection to number of probes is indicative of the robustness of the ranking list.

B. Genes’ robustness - Count the number of iterations in which each gene appeared among the 62 top selected ranked probes.

As one can see in Fig. S1, 91 genes appeared among the 100 sub-classifiers altogether, many of them appeared in all the 100 sub-classifiers, signifying a biological entity that is not determined by the training data, since it was repeated during the randomizations of the training sampling (that was used for the gene ranking in each iteration). We preferred having a classifier with less features (gene-probes) and with somewhat higher error rate, but more robust, than having a classifier that succeeding in having a bit lower error on the training set but with a risk of overfitting that is biased by noise of unstable gene ranking. Therefore, we trimmed the number of features as possible without harming the error rates extremely.

**Appendix S5. TCGA pre-processing**

The TCGA (10) BRCA expression RSEM normalized results file was used. p53 somatic mutations status and clinical information of the samples (i.e. ER, Her2 statuses) were downloaded from the Xena browser (11). For adjusting the TCGA expression data to be as similar as possible to the METABRIC expression, it was converted into log2 scale. Each gene was normalized by subtracting mean and dividing by its std, similarly to the METABRIC pre-processing.

The TCGA expression data was generated on an RNA-Seq based IlluminaGA system, which is quite different from the Illumina microarray method used for the METABRIC dataset. As a result, some of the genes used by the final classifier that was built based on the METABRIC dataset might not appear in the TCGA dataset. Moreover, the METABRIC analysis was done at probe level and the TCGA expression data is at gene level, meaning that for two distinct probes representing the same gene in METABRIC, in the TCGA we might have to use the same gene twice.

For each of the 100 final sub-classifiers we checked whether all their required probes are represented by a gene in the TCGA, if not, the sub-classifier is not usable. Then we collected all the 42 sub-classifiers which had all their required genes included in the TCGA expression data; Labeled each sample of ER+Her2- using the 42 sub-classifiers and finally calculated for each sample a majority vote on the 42 decisions. Consequently, each sample ended up with one classification of its p53 mutation status, based on the majority vote of 42 sub-classifiers.

**Appendix S6. Results of the learning process for ER-Her2- and for Her2+.**

The same learning process as in Fig. 1 was conducted also for ER-Her2- and Her2+ tumors (separately and independently). The error rates for the two final master classifiers are displayed below (Table S2 and Fig. S2) with the corresponding test errors for the METABRIC as well as for the TCGA.

**Appendix S7. Additional error curves for ER+Her2- tumors.**

For selecting one final constellation of GL, GR and ML method we compared the error curves of many combinations of the tunable parameters. After the final parameters were chosen (p53 related genes, Corr, Linear SVM) the number of features (probes) was chosen according to the following logic: we preferred having a more robust classifier than a classifier that uses many genes and could possibly overfit the training data (therefore we preferred to stop at 62 probes instead of a larger number with a slightly lower error rate).

In Fig. S3. One can see that (a) When using p53 related gene list the error rate is lower than for all genes (Fig. 3.A & 3.B). (b) Linear SVM returns the lower error rates than the ML methods (Fig. 3.A). (c) All genes list combined with FSW resulted in a similar error to p53 related genes combined with Corr GR, however since it was less robust, we selected p53 related genes combined with Corr GR.

**Appendix S8. Additional information about pseudomutant classification for ER+Her2- samples.**

The final master classifier consists of the majority vote of 100 sub-classifiers. Table S3 displays the number of FP (false positive) samples, meaning, all the wt p53 samples that were misclassified as mutants at least once (out of the labels of the 100 sub-classifiers). Fig. S4 and displays the classification of the 100 sub-classifiers independently and also the agreement between them, on which the final prediction is based (using majority vote), meaning, the fraction of the iterations in which wt p53 sample was classified as mutant out of the number of iterations the sample was chosen to be in the validation set.

A wt p53 sample from the learning set was labeled as pseudomutant only in case that in at least half of the iterations in which the sample was selected to be in the validation set it was classified as mutant. In case the sample was in the test set, it was called pseudomutant only if at least half of the sub-classifiers (>50) assigned a mutant classification to the sample.

**Appendix S9. Discrepancy between NGS and Sanger sequencing for TP53 mutations for the PM samples.**

After re-sequencing, three samples, out of the four samples classified as mut TP53 by Sanger but wt by NGS (see Table S6), were identified as carrying mutations in the TP53 gene: MB-0542, MB-4767 (from the learning set) and MB-7011 (from the test set).

*Table S1. Samples suspected to have germline mutations (n=15).*

| **chromosome** | **start** | **end** | **width** | **strand** | **ref** | **alt** | **depth** | **vaf** | **gene** | **exon** | **codon** | **class** | **location** | **oldAmino** | **newAmino** | **TSID** | **pubId** |
| --- | --- | --- | --- | --- | --- | --- | --- | --- | --- | --- | --- | --- | --- | --- | --- | --- | --- |
| chr17 | 7578388 | 7578388 | 1 | * | C | T | 174 | 0.793103 | TP53 | exon5 | 181 | nonsynonymous SNV | exonic | R | H | MTS-T0269 | MB-0005 |
| chr17 | 7576875 | 7576875 | 1 | * | T | C | 75 | 0.786667 | TP53 | exon9 | 324 | nonsynonymous SNV | exonic | D | G | MTS-T0294 | MB-0060 |
| chr17 | 7577103 | 7577103 | 1 | * | C | T | 167 | 0.431138 | TP53 | exon8 | 279 | nonsynonymous SNV | exonic | G | R | MTS-T0306 | MB-0093 |
| chr17 | 7576543 | 7576543 | 1 | * | C | T | 94 | 0.606383 | TP53 | NA | NA | NA | exonic | NA | NA | MTS-T0616 | MB-0180 |
| chr17 | 7577509 | 7577509 | 1 | * | C | T | 184 | 0.347826 | TP53 | exon7 | 258 | nonsynonymous SNV | exonic | E | K | MTS-T1759 | MB-0444 |
| chr17 | 7577575 | 7577575 | 1 | * | A | C | 175 | 0.262857 | TP53 | exon7 | 236 | nonsynonymous SNV | exonic | Y | D | MTS-T1035 | MB-0558 |
| chr17 | 7574026 | 7574026 | 1 | * | C | T | 110 | 0.627273 | TP53 | exon10 | 334 | nonsynonymous SNV | exonic | G | E | MTS-T1803 | MB-0590 |
| chr17 | 7579419 | 7579419 | 1 | * | A | AGG | 151 | 0.29801325 | TP53 | exon4 | 90 | frameshift substitution | exonic | NA | NA | MTS-T0128 | MB-7012 |
| chr17 | 7577509 | 7577509 | 1 | * | C | T | 201 | 0.218905 | TP53 | exon7 | 258 | nonsynonymous SNV | exonic | E | K | MTS-T0117 | MB-7035 |
| chr17 | 7577575 | 7577575 | 1 | * | A | C | 129 | 0.062016 | TP53 | exon7 | 236 | nonsynonymous SNV | exonic | Y | D | MTS-T0114 | MB-7081 |
| chr17 | 7576543 | 7576543 | 1 | * | C | T | 45 | 0.222222 | TP53 | NA | NA | NA | exonic | NA | NA | MTS-T1784 | MB-7253 |
| chr17 | 7576927 | 7576927 | 1 | * | C | T | 158 | 0.386076 | TP53 | exon9 | -1 | NA | splicing | NA | NA | MTS-T1100 | MB-7269 |
| chr17 | 7579320 | 7579322 | 3 | * | TCA | T | 155 | 0.16774194 | TP53 | exon4 | 122 | frameshift substitution | exonic | NA | NA | MTS-T0963 | MB-3556 |
| chr17 | 7576928 | 7576928 | 1 | * | T | C | 65 | 0.569231 | TP53 | exon9 | -1 | NA | splicing | NA | NA | MTS-T1409 | MB-5115 |
| chr17 | 7579320 | 7579322 | 3 | * | TCA | T | 207 | 0.09661836 | TP53 | exon4 | 122 | frameshift substitution | exonic | NA | NA | MTS-T0796 | MB-5323 |

*Table S2. Robustness of the final model’s gene lists (ER+Her2- using p53 related genes). The mean and std of the intersection and the total number of genes appearing among the 100 lists of top 62 ranked probes are displayed.*

| Final number of probes | Mean lists intersection | Std of lists intersection | #Total genes |
| --- | --- | --- | --- |
| 62 | 43.99 | 1.561 | 91 |

*Table S3. The final constellation of ML*, GR*, GL* and k* for ER-Her2- and Her2+ and the corresponding error rates.*

| Subtypes | #Training samples | #Validation samples | #Test samples | #Probes k* (mean ±SEM genes) | Selected  GR* (GL*) | Selected ML* method | Validation error mean ±SEM | Training error mean ±SEM | Test error | TCGA error (#valid sub-classifiers) |
| --- | --- | --- | --- | --- | --- | --- | --- | --- | --- | --- |
| TN | 202 | 50 | 63 | 19 (17.3±0.05) | Correlation  (all genes) | Linear SVM | 0.11±0.005 | 0.097±0.001 | 0.142 | 0.135 (80) |
| HER2+ | 154 | 38 | 48 | 27 (18.1±0.08) | Correlation  (all genes) | Gaussian SVM | 0.17± | 0.047± | 0.27 | 0.232 (20) |

*Table S4.* *The number of ER+Her2- samples that were falsely classified at least once during the training phase and the validation phase of the final classifiers. The numbers of mut and wt samples, and percentage of mutants at the training stages (without test data), are shown in the three right columns.*

| #FP train | #FP validation | #mut | #wt | %mut |
| --- | --- | --- | --- | --- |
| 62 | 81 | 200 | 898 | 18.21% |

*Table S5. PM samples (from ER+Her2-) that were recognized by the final classifier both METABRIC and TCGA datasets.*

| **METABRIC** | **TCGA** |
| --- | --- |
| 'MB-0054' | 'TCGA-A2-A0CY-01' |
| 'MB-0066' | 'TCGA-A2-A1FV-01' |
| 'MB-0131' | 'TCGA-A7-A0D9-01' |
| 'MB-0143' | 'TCGA-A8-A06N-01' |
| 'MB-0147' | 'TCGA-A8-A07G-01' |
| 'MB-0173' | 'TCGA-A8-A08F-01' |
| 'MB-0195' | 'TCGA-A8-A08L-01' |
| 'MB-0197' | 'TCGA-AN-A0AM-01' |
| 'MB-0311' | 'TCGA-AO-A03M-01' |
| 'MB-0393' | 'TCGA-AR-A0TW-01' |
| 'MB-0532' | 'TCGA-AR-A0TY-01' |
| 'MB-0542' | 'TCGA-AR-A24H-01' |
| 'MB-0543' | 'TCGA-BH-A0B1-01' |
| 'MB-0666' | 'TCGA-BH-A18P-01' |
| 'MB-2634' | 'TCGA-D8-A1J8-01' |
| 'MB-3389' |  |
| 'MB-3466' |  |
| 'MB-3600' |  |
| 'MB-4148' |  |
| 'MB-4235' |  |
| 'MB-4278' |  |
| 'MB-4767' |  |
| 'MB-4834' |  |
| 'MB-4876' |  |
| 'MB-4937' |  |
| 'MB-5001' |  |
| 'MB-5188' |  |
| 'MB-5350' |  |
| 'MB-5467' |  |
| 'MB-5521' |  |
| 'MB-5552' |  |
| 'MB-5575' |  |
| 'MB-5653' |  |
| 'MB-6060' |  |
| 'MB-6077' |  |
| 'MB-6208' |  |
| 'MB-6284' |  |
| 'MB-6287' |  |
| 'MB-7011' |  |
| 'MB-7043' |  |
| 'MB-7092' |  |
| 'MB-7133' |  |
| 'MB-7292' |  |
| 'MB-7299' |  |

*Table S6. Chosen parameters for the final imbalance-corrected model for ER+Her2- and its corresponding errors (training, validation, test and TCGA), with the number of models having all required genes in the TCGA (out of the 100 sub-classifiers).*

| **#Training samples** | **#Validation samples** | **#Probes k***  **(mean ±SEM genes)** | **Selected GL* (GR* )** | **Selected ML* method** | **Validation error**  **Mean**  **±SEM** | **Training error mean**  **±SEM** | **Test error (majority vote)** | **#Models with all genes in TCGA** | **TCGA test error (majority vote)** |
| --- | --- | --- | --- | --- | --- | --- | --- | --- | --- |
| 878 | 220 | 55  (43.83± 0.08) | p53 related genes (Corr) | Gaussian SVM | 0.125  ±0.0021 | 0.047  ±0.0005 | 0.1127 | 70 | 0.1086 |

*Table S7. Discrepancies between NGS and Sanger sequencing for the 44 pseudomutant (PM) samples.*

| SS\ NGS | wt | mut | Not sequenced | All |
| --- | --- | --- | --- | --- |
| wt | 30 | 0 | 2 | 32 |
| mut | 4 | 0 | 0 | 4 |
| Not sequenced | 8 | 0 | 0 | 8 |
| All | 42 | 0 | 2 | 44 |

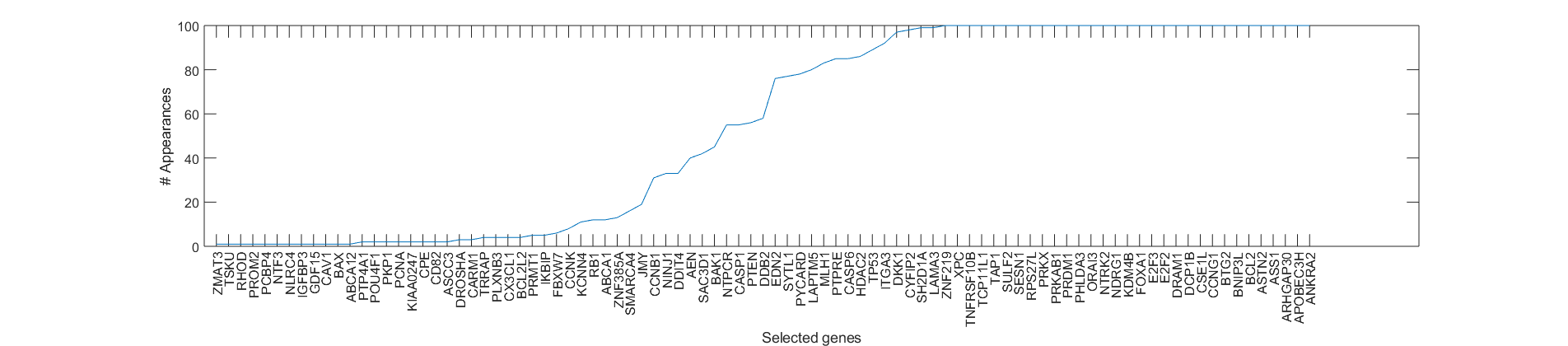

*Fig. S1. The number of appearance of each gene amongst the 62 top ranked probes (that were chosen to be used in at least one final sub-classifier) during the 100 iterations for ER+Her2- out of the p53 related genes list.*

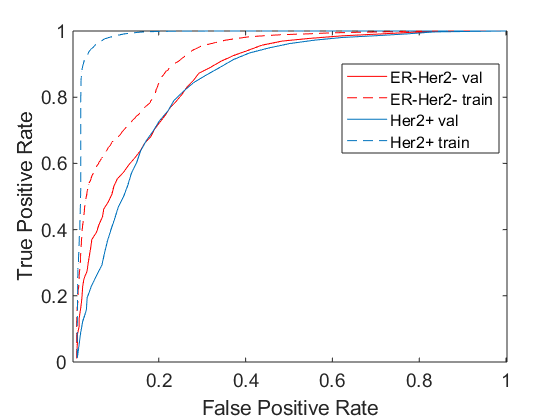

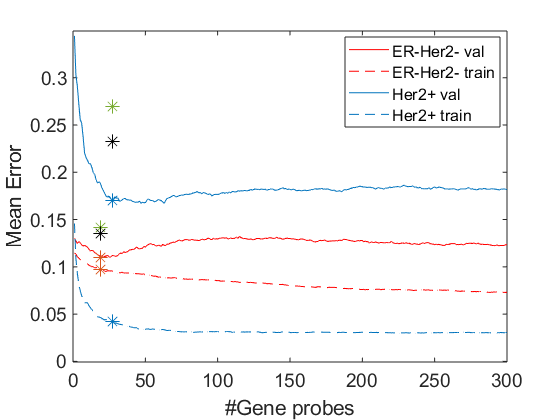

*Fig S2. Accuracy and error rates for the chosen models for ER-Her2- and Her2+ tumors.*

*A. Mean error rates of the predictions of p53 status (mut/wt) for the validation and training sets, using the chosen ML*, GR* and GL*, for 100 iterations, versus the number of top ranked gene-probes used. The test (METABRIC) and the TCGA errors of the final models are marked by green and black asterisks, respectively. The mean validation and training errors of the selected models are marked by blue and red asterisks.*

*B. Mean ROC Curves for the final models for ER-Her2- and Her2+ for both validation and training sets. The Area Under the Curve values for the validation curve and the training curve are 0.87 and 0.916 respectively for ER-Her2- and 0.85 and 0.99 respectively for Her2+.*

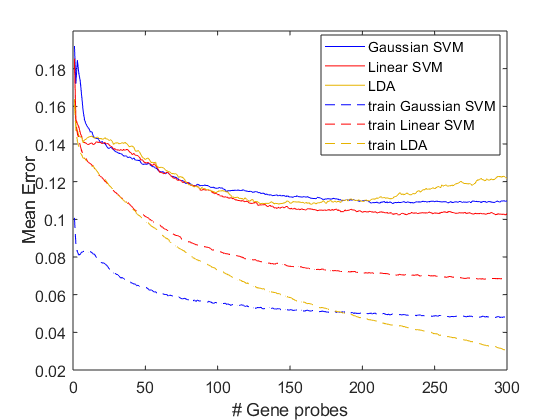

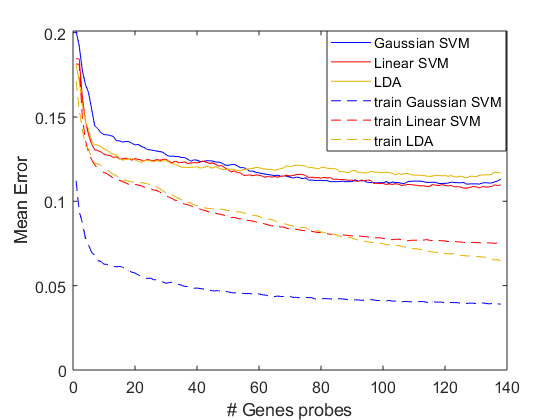

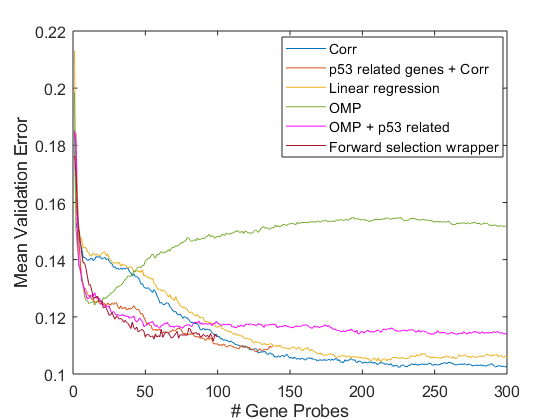

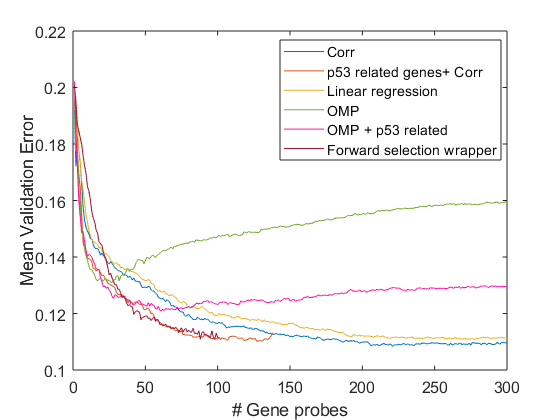

*Fig. S3. Additional error rates for ER+Her2- tumors, each based on 100 iterations (and therefore 100 error rates).*

*(A) Different ML methods using p53-related genes.*

*(B) Different ML methods using all genes.*

*(C) Different GR methods using linear SVM.*

*(D) Different GR methods using Gaussian SVM.*

**D**

**C**

**B**

**A**

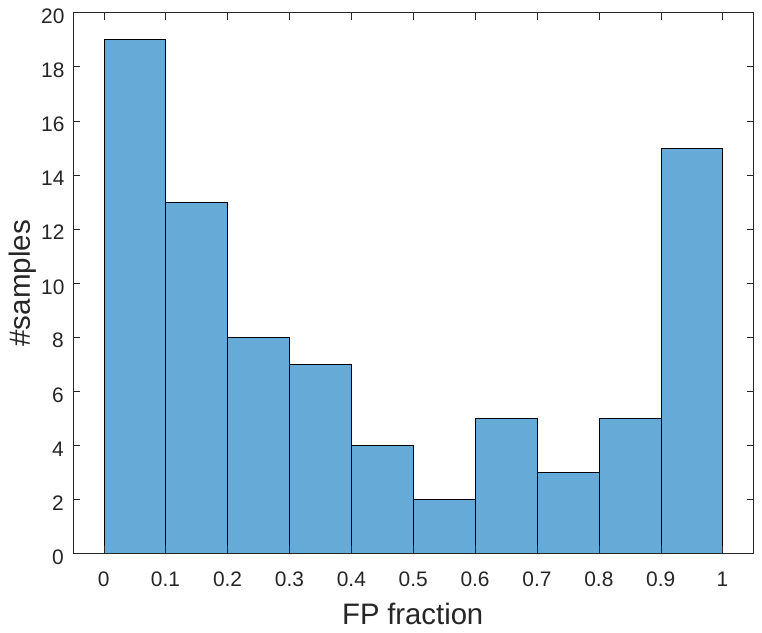

*Fig. S4. The number of wt p53 ER+Her2- samples from the validation set, which were misclassified as false positive (mut p53 classification) at least once (n=81). The horizontal axis shows the fraction of misclassifications out of the number of appearances of each sample in the validation set. For example, if an ER+Her2- wt p53 sample s was assigned to the validation set in 20 iterations, and was misclassified to be mutant 19 times, S will be in the rightmost column (0.9<%FP<1.0) of the left panel; the number of such samples is 15.*

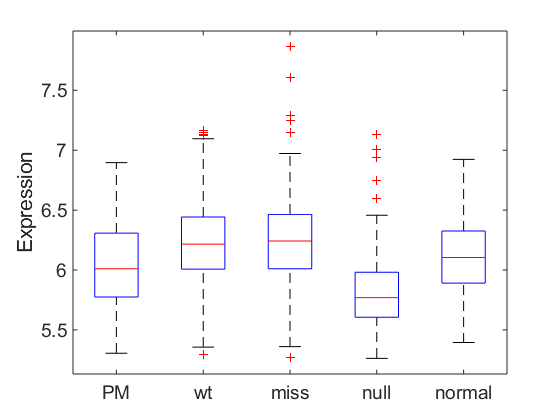

*Fig. S5. Comparison of TP53 mRNA expression in the different groups of ER+Her2- tumors and in normal human breast tissue.*

*PM = pseudomutant; miss = missense TP53 mutated; null = nonsense and frameshift TP53 mutations.*
