## Supplemental File 1 for "Transcriptional profiling reveals a subset of human breast tumors that retain wt *TP53* but display mutant p53-associated features"

Final expression table of 34,363 (HT-12 v3 platform, Illumina_Human_WG-v3) gene-probes representing 24,369 genes, for 1928 tumor and 144 normal samples.

The data file is too large to upload onto the website of the journal: it is available upon request from the authors.
