## Supplemental File 2 for "Transcriptional profiling reveals a subset of human breast tumors that retain wt *TP53* but display mutant p53-associated features"

TP53 labels, the mutation types and breast cancer subtypes (ER+Her2-, ER-Her2- and Her2+, as determined by Immunohistochemistry), protein change, survival and treatment information and clinical parameters.

| Row | MB_0362 | MB_0346 | MB_0386 | MB_0574 | MB_0185 | MB_0503 | MB_0641 | MB_0201 | MB_0218 | MB_0316 | MB_0189 | MB_0891 |
|---|---|---|---|---|---|---|---|---|---|---|---|---|
| p53_labels | WT | MUT | MUT | MUT | MUT | WT | WT | MUT | WT | MUT | WT | WT |
| mutation_type | WT | MISS | NULL | MISS | MISS | WT | WT | MISS | WT | MISS | WT | WT |
| protein_change | | R273C | E349Gfs*30 | Y236C | p.P278S | | | G245D | | P151S | | |
| subtype | ERpHER2n | HER2p | ERpHER2n | ERpHER2n | ERpHER2n | ERpHER2n | ERpHER2n | HER2p | ERpHER2n | ERnHER2n | ERpHER2n | ERpHER2n |
| survival_status | Died of Disease | Died of Disease | Living | Living | Died of Disease | Living | Living | Living | Living | Living | Died of Disease | Living |
| survival_months | 47.03333333 | 20.43333333 | 138.1333333 | 119.8 | 43.83333333 | 101.2333333 | 102.5666667 | 125.7 | 131.0666667 | 182.9 | 9.066666667 | 149.4 |
| chemotherapy | YES | NO | NO | NO | NO | NO | NO | YES | YES | NO | YES | NO |

| | | | | | | | | | | | | |
|---|---|---|---|---|---|---|---|---|---|---|---|---|
| MB_0658 | MB_0899 | MB_0605 | MB_0258 | MB_0506 | MB_0420 | MB_0223 | MB_0445 | MB_0199 | MB_0517 | MB_0155 | MB_0428 | MB_0117 |
| MUT | WT | WT | WT | MUT | MUT | WT | WT | WT | WT | WT | WT | WT |
| MISS | WT | WT | WT | MISS | NULL | WT | WT | WT | WT | WT | WT | WT |
| R248Q | | | | C135W | R306* | | | | | | | |
| ERnHER2n | ERpHER2n | ERpHER2n | ERpHER2n | ERnHER2n | ERnHER2n | ERpHER2n | ERpHER2n | ERpHER2n | ERpHER2n | ERpHER2n | ERpHER2n | ERpHER2n |
| Living | Living | Living | Living | Died of Disease | Living | Died of Disease | Living | Living | Living | Living | Living | Living |
| 97.26666667 | 175.8 | 114.6 | 86.13333333 | 45.8 | 76.73333333 | 81.33333333 | 132.5333333 | 144.4 | 61.9 | 59.8 | 55.93333333 | 2.4 |
| YES | YES | NO | NO | YES | YES | YES | YES | NO | NO | NO | NO | NO |

| | | | | | | | | | | |
|---|---|---|---|---|---|---|---|---|---|---|
| MB_0906 | MB_0249 | MB_0660 | MB_0497 | MB_0434 | MB_0143 | MB_0513 | MB_0541 | MB_0653 | MB_0455 | MB_0540 |
| MUT | MUT | MUT | WT | WT | WT | WT | WT | MUT | MUT | WT |
| MISS | MISS | MISS | WT | WT | WT | WT | WT | MISS | NULL | WT |
| L111P | Y205S | C141Y | | | | | | E285K | S166* | |
| ERnHER2n | ERnHER2n | ERnHER2n | ERpHER2n | HER2p | ERpHER2n | ERpHER2n | ERpHER2n | ERnHER2n | ERpHER2n | ERnHER2n |
| Died of Other Causes | Living | Died of Disease | Living | Died of Disease | Died of Other Causes | Living | Living | Died of Disease | Living | Living |
| 63.86666667 | 188.3333333 | 18.93333333 | 73.7 | 45.5 | 54.33333333 | 117.0333333 | 174.2666667 | 24.8 | 99.33333333 | 194.5333333 |
| NO | NO | YES | NO | YES | NO | NO | YES | NO | NO | NO |

| | | | | | | | | | | |
|---|---|---|---|---|---|---|---|---|---|---|
| MB_0384 | MB_0637 | MB_0157 | MB_0443 | MB_0584 | MB_0292 | MB_0322 | MB_0501 | MB_0401 | MB_0140 | MB_0606 |
| WT | MUT | MUT | WT | WT | MUT | WT | MUT | MUT | WT | WT |
| WT | MISS | MISS | WT | WT | NULL | WT | MISS | NULL | WT | WT |
| | V272G | Y163C | | | I195Yfs*14 | | P27L | D148Pfs*21 | | |
| ERpHER2n | ERpHER2n | ERnHER2n | ERpHER2n | ERpHER2n | ERnHER2n | ERpHER2n | ERpHER2n | ERnHER2n | ERpHER2n | ERpHER2n |
| Died of Disease | Living | Living | Living | Living | Died of Other Causes | Died of Other Causes | Living | Died of Disease | Living | Died of Disease |
| 61.1 | 80.66666667 | 114.7666667 | 45.03333333 | 118.2 | 49.43333333 | 50.53333333 | 71.5 | 26.26666667 | 147.9333333 | 27.46666667 |
| NO | NO | NO | NO | NO | NO | NO | YES | YES | NO | NO |

| | | | | | | | | | | | |
|---|---|---|---|---|---|---|---|---|---|---|---|
| MB_0666 | MB_0598 | MB_0453 | MB_0138 | MB_0579 | MB_0471 | MB_0347 | MB_0619 | MB_0171 | MB_0310 | MB_0621 | MB_0614 |
| WT | WT | MUT | WT | WT | WT | MUT | WT | WT | WT | WT | WT |
| WT | WT | NULL | WT | WT | WT | NULL | WT | WT | WT | WT | WT |
| | | Q165* | | | | p.E339X | | | | | |
| ERpHER2n | ERpHER2n | ERnHER2n | ERpHER2n | ERpHER2n | ERpHER2n | ERpHER2n | ERpHER2n | ERpHER2n | ERpHER2n | ERpHER2n | ERpHER2n |
| Died of Disease | Died of Disease | Died of Disease | Living | Died of Other Causes | Living | Died of Disease | Living | Living | Living | Living | Living |
| 24.33333333 | 89.36666667 | 43.13333333 | 150.5666667 | 92.76666667 | 107.4666667 | 205.5666667 | 94.23333333 | 5.433333333 | 185.1666667 | 33.56666667 | 119.3333333 |
| YES | NO | YES | NO | NO | NO | YES | YES | NO | NO | NO | YES |

| | | | | | | | | | | | |
|---|---|---|---|---|---|---|---|---|---|---|---|
| MB_0372 | MB_0374 | MB_0382 | MB_0327 | MB_0066 | MB_0144 | MB_0596 | MB_0164 | MB_0215 | MB_0146 | MB_0229 | MB_0505 |
| MUT | WT | MUT | WT | WT | WT | WT | MUT | WT | WT | WT | WT |
| NULL | WT | MISS | WT | WT | WT | WT | MISS | WT | WT | WT | WT |
| E285* | | R337L | | | | | R175H | | | | |
| ERnHER2n | ERpHER2n | ERpHER2n | ERpHER2n | ERpHER2n | ERpHER2n | ERpHER2n | ERnHER2n | ERpHER2n | ERpHER2n | ERpHER2n | ERpHER2n |
| Died of Disease | Living | Living | Living | Living | Died of Disease | Died of Other Causes | Living | Living | Died of Other Causes | Living | Living |
| 62.76666667 | 1.433333333 | 136.4666667 | 187.9333333 | 157.4333333 | 152.0666667 | 100.4666667 | 10.83333333 | 122.2 | 86 | 71.5 | 60.7 |
| YES | NO | YES | NO | NO | NO | NO | YES | NO | NO | NO | NO |

| | | | | | | | | | | |
|---|---|---|---|---|---|---|---|---|---|---|
| MB_0102 | MB_0569 | MB_0516 | MB_0272 | MB_0585 | MB_0494 | MB_0306 | MB_0463 | MB_0198 | MB_0203 | MB_0607 |
| MUT | WT | MUT | WT | MUT | MUT | MUT | WT | WT | WT | MUT |
| MISS | WT | MISS | WT | MISS | MISS | MISS | WT | WT | WT | MISS |
| R175H | | R175H | | R175H | P278S | G266E | | | | E285K |
| ERpHER2n | ERpHER2n | ERnHER2n | ERpHER2n | ERpHER2n | ERnHER2n | ERpHER2n | ERpHER2n | ERpHER2n | ERpHER2n | ERpHER2n |
| Died of Disease | Died of Disease | Living | Died of Disease | Died of Disease | Died of Disease | Died of Disease | Died of Disease | Living | Died of Other Causes | Living |
| 140.7666667 | 59.5 | 114.4666667 | 122 | 106.9666667 | 36.63333333 | 42.96666667 | 61.96666667 | 144.7666667 | 58.76666667 | 87.83333333 |
| YES | YES | YES | NO | NO | YES | NO | NO | YES | YES | YES |

| | | | | | | | | | | |
|---|---|---|---|---|---|---|---|---|---|---|
| MB_0631 | MB_0363 | MB_0427 | MB_0519 | MB_0371 | MB_0380 | MB_0221 | MB_0348 | MB_0261 | MB_0576 | MB_0385 |
| WT | WT | WT | MUT | WT | MUT | MUT | WT | WT | WT | WT |
| WT | WT | WT | NULL | WT | MISS | MISS | WT | WT | WT | WT |
| | | | X331_splice | | N239D | V197G | | | | |
| ERpHER2n | ERpHER2n | ERpHER2n | ERpHER2n | HER2p | ERpHER2n | ERnHER2n | ERpHER2n | ERpHER2n | ERpHER2n | ERpHER2n |
| Died of Disease | Died of Disease | Living | Living | Living | Died of Other Causes | Died of Disease | Died of Other Causes | Died of Disease | Died of Disease | Died of Disease |
| 47.43333333 | 89.9 | 116.1 | 110.9666667 | 131.1333333 | 69.33333333 | 20.2 | 122.7 | 90.23333333 | 31 | 36.43333333 |
| YES | NO | NO | YES | NO | NO | NO | NO | NO | NO | YES |

| | | | | | | | | | | |
|---|---|---|---|---|---|---|---|---|---|---|
| MB_0659 | MB_0270 | MB_0379 | MB_0527 | MB_0624 | MB_0273 | MB_0050 | MB_0460 | MB_0654 | MB_0454 | MB_0392 |
| WT | WT | WT | WT | WT | WT | WT | WT | WT | WT | WT |
| WT | WT | WT | WT | WT | WT | WT | WT | WT | WT | WT |
| ERnHER2n | ERpHER2n | ERpHER2n | ERpHER2n | ERpHER2n | ERpHER2n | ERpHER2n | ERpHER2n | ERpHER2n | ERpHER2n | ERpHER2n |
| Died of Disease | Living | Living | Died of Other Causes | Living | Living | Living | Died of Disease | Living | Died of Other Causes | Died of Disease |
| 23.33333333 | 337.0333333 | 135.6666667 | 109.8333333 | 64.03333333 | 186.5333333 | 75.33333333 | 114 | 69.4 | 46.83333333 | 38.13333333 |
| NO | NO | YES | NO | NO | NO | YES | NO | NO | NO | YES |

| | | | | | | | | | | |
|---|---|---|---|---|---|---|---|---|---|---|
| MB_0336 | MB_0467 | MB_0349 | MB_0378 | MB_0176 | MB_0429 | MB_0397 | MB_0571 | MB_0426 | MB_0135 | MB_0112 |
| WT | MUT | WT | MUT | WT | WT | WT | WT | WT | WT | WT |
| WT | NULL | WT | NULL | WT | WT | WT | WT | WT | WT | WT |
| | S90Pfs*33 | | S20Qfs*24 | | | | | | | |
| ERpHER2n | HER2p | ERpHER2n | HER2p | ERpHER2n | ERpHER2n | ERpHER2n | ERpHER2n | ERpHER2n | ERpHER2n | ERpHER2n |
| Died of Disease | Died of Disease | Died of Other Causes | Died of Disease | Living | Died of Other Causes | Living | Living | Living | Living | Died of Disease |
| 170.3 | 105.6 | 146.0333333 | 97.3 | 113.4333333 | 74.46666667 | 57.23333333 | 149.8666667 | 131.1 | 116.6333333 | 39.16666667 |
| NO | YES | NO | YES | YES | NO | NO | NO | YES | NO | NO |

| | | | | | | | | | | | |
|---|---|---|---|---|---|---|---|---|---|---|---|
| MB_0352 | MB_0644 | MB_0601 | MB_0568 | MB_0328 | MB_0325 | MB_0358 | MB_0413 | MB_0158 | MB_0636 | MB_0145 | MB_0195 |
| WT | WT | MUT | WT | WT | MUT | MUT | WT | MUT | MUT | WT | WT |
| WT | WT | MISS | WT | WT | NULL | NULL | WT | NULL | MISS | WT | WT |
| | | Y126C | | | R213Dfs*34 | X225_splice | | p.? | R273H | | |
| ERnHER2n | ERpHER2n | ERpHER2n | ERpHER2n | ERpHER2n | ERpHER2n | ERpHER2n | ERpHER2n | ERnHER2n | ERpHER2n | ERpHER2n | ERpHER2n |
| Died of Disease | Living | Died of Disease | Living | Died of Disease | Living | Living | Living | Living | Died of Disease | Living | Living |
| 55.2 | 93.3 | 29.66666667 | 181.4666667 | 125.6 | 177.5333333 | 32.86666667 | 140.0666667 | 192.3 | 32.63333333 | 147.6666667 | 146 |
| YES | NO | YES | NO | NO | NO | YES | YES | YES | NO | NO | NO |

| | | | | | | | | | |
|---|---|---|---|---|---|---|---|---|---|
| MB_0422 | MB_0483 | MB_0317 | MB_0486 | MB_0139 | MB_0257 | MB_0345 | MB_0375 | MB_0419 | MB_0480 |
| WT | WT | WT | WT | WT | WT | WT | MUT | WT | MUT |
| WT | WT | WT | WT | WT | WT | WT | NULL | WT | NULL |
| | | | | | | | A74Sfs*71 | | F338Rfs*8 |
| ERpHER2n | ERpHER2n | ERpHER2n | ERpHER2n | ERpHER2n | ERpHER2n | ERpHER2n | ERpHER2n | ERpHER2n | ERpHER2n |
| Died of Other Causes | Died of Disease | Died of Other Causes | Living | Living | Died of Disease | Living | Living | Died of Other Causes | Living |
| 33.8 | 74.46666667 | 151.6666667 | 89.96666667 | 109.2 | 91.73333333 | 72.66666667 | 143.1666667 | 104.3 | 94.56666667 |
| NO | YES | NO | YES | NO | YES | YES | NO | NO | NO |

| | | | | | | | | | | |
|---|---|---|---|---|---|---|---|---|---|---|
| MB_0311 | MB_0324 | MB_0368 | MB_0389 | MB_0248 | MB_0035 | MB_0667 | MB_0423 | MB_0904 | MB_0119 | MB_0650 |
| WT | WT | WT | WT | WT | WT | WT | WT | WT | WT | WT |
| WT | WT | WT | WT | WT | WT | WT | WT | WT | WT | WT |
| ERpHER2n | ERpHER2n | ERpHER2n | HER2p | ERpHER2n | ERpHER2n | HER2p | ERpHER2n | ERpHER2n | ERpHER2n | ERpHER2n |
| Died of Other Causes | Died of Disease | Died of Disease | Living | Living | Died of Disease | Died of Disease | Living | Died of Other Causes | Died of Disease | Living |
| 151.1666667 | 144.9666667 | 61.9 | 96.96666667 | 71 | 36.26666667 | 128 | 138.9333333 | 144.7 | 95.86666667 | 25.53333333 |
| NO | YES | YES | YES | YES | NO | NO | NO | NO | NO | NO |

| | | | | | | | | | | | | |
|---|---|---|---|---|---|---|---|---|---|---|---|---|
| MB_0204 | MB_0184 | MB_0600 | MB_0400 | MB_0511 | MB_0059 | MB_0500 | MB_0150 | MB_0895 | MB_0366 | MB_0173 | MB_0131 | MB_0206 |
| WT | WT | WT | MUT | WT | WT | MUT | MUT | MUT | WT | WT | WT | MUT |
| WT | WT | WT | MISS | WT | WT | NULL | MISS | MISS | WT | WT | WT | MISS |
| | | | R273C | | | R110Pfs*39 | K132N | G244S | | | | C135W |
| ERpHER2n | ERpHER2n | ERpHER2n | ERnHER2n | ERpHER2n | ERpHER2n | ERnHER2n | ERpHER2n | HER2p | ERpHER2n | ERpHER2n | ERpHER2n | ERnHER2n |
| Died of Disease | Living | Living | Died of Disease | Living | Living | Living | Living | Died of Other Causes | Living | Living | Died of Disease | Living |
| 24.3 | 109.0333333 | 111 | 22.46666667 | 61.7 | 160.9 | 67.46666667 | 111.2 | 43.1 | 64.23333333 | 3.766666667 | 66.63333333 | 141.1666667 |
| NO | NO | NO | YES | NO | NO | YES | NO | NO | NO | NO | YES | YES |

| | | | | | | | | | | |
|---|---|---|---|---|---|---|---|---|---|---|
| MB_0315 | MB_0361 | MB_0545 | MB_0370 | MB_0642 | MB_0431 | MB_0181 | MB_0603 | MB_0295 | MB_0618 | MB_0496 |
| MUT | MUT | MUT | WT | WT | WT | WT | WT | WT | WT | WT |
| NULL | MISS | MISS | WT | WT | WT | WT | WT | WT | WT | WT |
| P191Lfs*56 | R248Q | C242S | | | | | | | | |
| ERpHER2n | HER2p | ERpHER2n | ERpHER2n | ERpHER2n | ERpHER2n | ERpHER2n | ERpHER2n | ERpHER2n | ERpHER2n | ERpHER2n |
| Living | Died of Other Causes | Died of Other Causes | Died of Other Causes | Died of Other Causes | Living | Living | Living | Living | Living | Living |
| 186.2 | 15.06666667 | 41.16666667 | 93.5 | 84.2 | 85.13333333 | 85.96666667 | 121.9666667 | 164.5 | 101.9 | 130.9 |
| YES | NO | NO | YES | NO | NO | NO | NO | NO | NO | YES |

| | | | | | | | | | | | | |
|---|---|---|---|---|---|---|---|---|---|---|---|---|
| MB_0411 | MB_0285 | MB_0360 | MB_0359 | MB_0344 | MB_0583 | MB_0202 | MB_0485 | MB_0609 | MB_0538 | MB_0197 | MB_0410 | MB_0528 |
| MUT | WT | WT | WT | MUT | WT | MUT | MUT | MUT | WT | WT | WT | MUT |
| NULL | WT | WT | WT | MISS | WT | MISS | MISS | MISS | WT | WT | WT | NULL |
| G245del | | | | E285K | | D281H | C238R | R249S | | | | Q104* |
| ERpHER2n | ERpHER2n | ERpHER2n | ERpHER2n | ERpHER2n | ERpHER2n | ERpHER2n | ERpHER2n | ERpHER2n | ERpHER2n | ERpHER2n | ERpHER2n | ERpHER2n |
| Living | Living | Died of Disease | Living | Living | Living | Living | Died of Disease | Died of Disease | Died of Disease | Living | Living | Living |
| 139.1666667 | 3.366666667 | 132.5666667 | 21.6 | 152.9333333 | 62.63333333 | 128.3666667 | 36.76666667 | 98.7 | 24.1 | 70.73333333 | 63 | 193.9666667 |
| YES | NO | NO | NO | NO | NO | YES | YES | YES | YES | YES | NO | YES |

| | | | | | | | | | | | |
|---|---|---|---|---|---|---|---|---|---|---|---|
| MB_0165 | MB_0152 | MB_0148 | MB_0594 | MB_0521 | MB_0532 | MB_0536 | MB_0319 | MB_0491 | MB_0404 | MB_0243 | MB_0580 |
| MUT | MUT | MUT | WT | WT | WT | MUT | WT | WT | WT | WT | MUT |
| MISS | NULL | NULL | WT | WT | WT | MISS | WT | WT | WT | WT | NULL |
| H214R | G266Rfs*74 | Q192* | | | | H179R | | | | | H214Lfs*33 |
| HER2p | HER2p | HER2p | ERpHER2n | ERpHER2n | ERpHER2n | ERpHER2n | ERpHER2n | ERpHER2n | ERpHER2n | ERpHER2n | ERpHER2n |
| Died of Disease | Living | Living | Living | Living | Died of Disease | Died of Other Causes | Died of Disease | Living | Living | Living | Died of Disease |
| 47.63333333 | 63.03333333 | 1.766666667 | 44.23333333 | 212.2 | 143.1333333 | 147.3666667 | 40.7 | 125.8 | 63.5 | 149.7666667 | 79.36666667 |
| YES | NO | NO | NO | NO | YES | NO | NO | NO | NO | YES | YES |

| | | | | | | | | | | | |
|---|---|---|---|---|---|---|---|---|---|---|---|
| MB_0312 | MB_0635 | MB_0649 | MB_0643 | MB_0581 | MB_0383 | MB_0142 | MB_0320 | MB_0207 | MB_0167 | MB_0591 | MB_0613 |
| WT | WT | WT | MUT | MUT | WT | WT | WT | WT | MUT | WT | MUT |
| WT | WT | WT | NULL | MISS | WT | WT | WT | WT | MISS | WT | NULL |
| | | | T304Nfs*42 | L194R | | | | | R282W | | E287* |
| ERpHER2n | ERnHER2n | ERpHER2n | HER2p | ERnHER2n | ERpHER2n | ERpHER2n | ERpHER2n | ERpHER2n | ERpHER2n | ERpHER2n | ERnHER2n |
| Died of Disease | Living | Living | Died of Other Causes | Died of Disease | Died of Disease | Living | Living | Living | Died of Disease | Died of Disease | Living |
| 169 | 88 | 61.43333333 | 19.73333333 | 7.866666667 | 85.73333333 | 12.4 | 63.73333333 | 173.6333333 | 43.1 | 117.7666667 | 48.8 |
| NO | NO | NO | NO | NO | NO | NO | NO | NO | NO | NO | YES |

| | | | | | | | | | | | | |
|---|---|---|---|---|---|---|---|---|---|---|---|---|
| MB_0313 | MB_0599 | MB_0286 | MB_0575 | MB_0549 | MB_0214 | MB_0589 | MB_0406 | MB_0452 | MB_0524 | MB_0880 | MB_0573 | MB_0664 |
| MUT | WT | WT | WT | WT | MUT | WT | WT | MUT | MUT | WT | WT | MUT |
| MISS | WT | WT | WT | WT | MISS | WT | WT | MISS | MISS | WT | WT | NULL |
| Y220C | | | | | V157F | | | R248Q | C275Y | | | E271Gfs*74 |
| ERpHER2n | ERpHER2n | ERpHER2n | ERpHER2n | HER2p | ERnHER2n | HER2p | ERpHER2n | ERpHER2n | ERpHER2n | ERpHER2n | ERpHER2n | ERnHER2n |
| Living | Living | Living | Living | Living | Living | Living | Died of Disease | Living | Living | Living | Died of Other Causes | Died of Disease |
| 184.7666667 | 123.2666667 | 192.6 | 128.5666667 | 25.96666667 | 146.9 | 125.8 | 96.9 | 62.8 | 80.83333333 | 1.233333333 | 163.2 | 11.06666667 |
| NO | YES | NO | YES | NO | YES | NO | YES | NO | YES | NO | NO | YES |

| | | | | | | | | | | |
|---|---|---|---|---|---|---|---|---|---|---|
| MB_0446 | MB_0008 | MB_0656 | MB_0154 | MB_0597 | MB_0550 | MB_0616 | MB_0448 | MB_0412 | MB_0122 | MB_0425 |
| MUT | MUT | MUT | WT | WT | WT | WT | WT | MUT | WT | WT |
| MISS | MISS | MISS | WT | WT | WT | WT | WT | NULL | WT | WT |
| C141R | S241F | V216L | | | | | | T102Pfs*21 | | |
| ERnHER2n | ERpHER2n | HER2p | ERpHER2n | ERpHER2n | ERpHER2n | ERpHER2n | ERpHER2n | ERpHER2n | ERpHER2n | ERpHER2n |
| Living | Died of Disease | Died of Disease | Living | Died of Other Causes | Died of Disease | Died of Other Causes | Died of Disease | Living | Living | Died of Other Causes |
| 72.26666667 | 41.36666667 | 19.03333333 | 114.7666667 | 113.0666667 | 132.3333333 | 29.03333333 | 29.23333333 | 136.1666667 | 138.9 | 24.4 |
| YES | YES | YES | NO | YES | NO | NO | YES | NO | NO | NO |

| | | | | | | | | | | | |
|---|---|---|---|---|---|---|---|---|---|---|---|
| MB_0314 | MB_0356 | MB_0440 | MB_0398 | MB_0438 | MB_0449 | MB_0162 | MB_0593 | MB_0301 | MB_0628 | MB_0283 | MB_0106 |
| MUT | WT | WT | MUT | WT | WT | WT | WT | WT | WT | WT | WT |
| NULL | WT | WT | NULL | WT | WT | WT | WT | WT | WT | WT | WT |
| X224_splice | | | D148Hfs*31 | | | | | | | | |
| HER2p | ERpHER2n | ERpHER2n | ERpHER2n | HER2p | ERpHER2n | ERpHER2n | HER2p | ERpHER2n | ERpHER2n | ERpHER2n | ERpHER2n |
| Died of Disease | Living | Living | Died of Disease | Living | Living | Living | Died of Disease | Died of Other Causes | Living | Died of Disease | Living |
| 4.433333333 | 213.5 | 100.8333333 | 43.2 | 135.1666667 | 112.5666667 | 55.76666667 | 34.66666667 | 122.7 | 105.9666667 | 22.03333333 | 85.33333333 |
| NO | NO | YES | NO | YES | YES | NO | YES | NO | NO | NO | YES |

| | | | | | | | | | | | |
|---|---|---|---|---|---|---|---|---|---|---|---|
| MB_0341 | MB_0474 | MB_0394 | MB_0373 | MB_0247 | MB_0225 | MB_0424 | MB_0209 | MB_0115 | MB_0136 | MB_0542 | MB_0354 |
| WT | MUT | WT | MUT | WT | WT | WT | WT | MUT | WT | WT | MUT |
| WT | MISS | WT | MISS | WT | WT | WT | WT | MISS | WT | WT | MISS |
| | G245V | | Y234H | | | | | R175H | | | H178P |
| ERpHER2n | ERpHER2n | ERpHER2n | HER2p | ERpHER2n | HER2p | ERnHER2n | ERpHER2n | ERnHER2n | ERpHER2n | ERpHER2n | ERnHER2n |
| Died of Disease | Died of Disease | Living | Died of Disease | Living | Living | Died of Disease | Living | Died of Disease | Living | Died of Disease | Died of Disease |
| 170.0333333 | 111.1 | 64.23333333 | 27 | 185.6 | 212.2 | 9.833333333 | 82.36666667 | 66.73333333 | 88.2 | 48.6 | 11.6 |
| NO | YES | NO | NO | NO | NO | YES | NO | YES | NO | YES | NO |

| | | | | | | | | | | | |
|---|---|---|---|---|---|---|---|---|---|---|---|
| MB_0151 | MB_0608 | MB_0657 | MB_0559 | MB_0893 | MB_0514 | MB_0395 | MB_0294 | MB_0439 | MB_0481 | MB_0529 | MB_0224 |
| WT | MUT | WT | WT | WT | WT | MUT | WT | WT | MUT | WT | WT |
| WT | MISS | WT | WT | WT | WT | NULL | WT | WT | NULL | WT | WT |
| | R175H | | | | | H115Afs*34 | | | Y107* | | |
| ERpHER2n | ERnHER2n | ERpHER2n | ERpHER2n | ERnHER2n | ERpHER2n | HER2p | ERnHER2n | ERpHER2n | ERnHER2n | ERpHER2n | ERpHER2n |
| Died of Disease | Died of Other Causes | Living | Living | Living | Living | Died of Disease | Living | Living | Died of Disease | Died of Disease | Living |
| 49.23333333 | 63.96666667 | 30.43333333 | 161.7666667 | 175.6333333 | 13.4 | 29 | 72.9 | 112.1333333 | 36.4 | 48.53333333 | 176.7666667 |
| NO | NO | NO | YES | YES | YES | NO | YES | NO | YES | YES | NO |

| | | | | | | | | | | | |
|---|---|---|---|---|---|---|---|---|---|---|---|
| MB_0302 | MB_0126 | MB_0220 | MB_0192 | MB_0121 | MB_0239 | MB_0364 | MB_0232 | MB_0884 | MB_0238 | MB_0194 | MB_0882 |
| WT | WT | WT | WT | WT | WT | WT | WT | MUT | MUT | WT | MUT |
| WT | WT | WT | WT | WT | WT | WT | WT | NULL | MISS | WT | NULL |
| | | | | | | | | V218Hfs*5 | R248Q | | R213* |
| ERpHER2n | ERpHER2n | ERnHER2n | ERpHER2n | ERpHER2n | ERpHER2n | ERpHER2n | ERpHER2n | ERpHER2n | ERnHER2n | ERpHER2n | ERpHER2n |
| Died of Disease | Living | Died of Disease | Living | Living | Died of Disease | Living | Living | Died of Disease | Living | Living | Died of Disease |
| 84.2 | 127.6333333 | 39.83333333 | 144.6333333 | 152.2 | 85.56666667 | 176.6 | 207.6333333 | 93.66666667 | 193.1666667 | 91.6 | 42.36666667 |
| NO | NO | NO | NO | NO | NO | YES | NO | YES | YES | YES | NO |

| | | | | | | | | | | | |
|---|---|---|---|---|---|---|---|---|---|---|---|
| MB_0010 | MB_0236 | MB_0377 | MB_0123 | MB_0504 | MB_0564 | MB_0475 | MB_0321 | MB_0482 | MB_0101 | MB_0662 | MB_0291 |
| MUT | MUT | WT | WT | WT | WT | WT | WT | MUT | WT | MUT | WT |
| NULL | MISS | WT | WT | WT | WT | WT | WT | MISS | WT | MISS | WT |
| P67Qfs*56 | N239D | | | | | | | R337C | | D281G | |
| ERpHER2n | HER2p | ERpHER2n | ERpHER2n | ERpHER2n | HER2p | ERpHER2n | ERpHER2n | HER2p | ERpHER2n | HER2p | HER2p |
| Died of Disease | Living | Living | Died of Other Causes | Living | Died of Disease | Living | Living | Died of Disease | Living | Died of Disease | Died of Disease |
| 7.8 | 205.0333333 | 134.3666667 | 114.2333333 | 130.4666667 | 37.93333333 | 130.7 | 174.6333333 | 24.86666667 | 148.0333333 | 69.8 | 39.2 |
| NO | YES | NO | NO | NO | YES | NO | YES | YES | NO | NO | YES |

| | | | | | | | | | | | | |
|---|---|---|---|---|---|---|---|---|---|---|---|---|
| MB_0465 | MB_0630 | MB_0036 | MB_0166 | MB_0525 | MB_0307 | MB_0002 | MB_0408 | MB_0466 | MB_0632 | MB_0120 | MB_0507 | MB_0287 |
| MUT | WT | WT | WT | MUT | WT | MUT | WT | WT | WT | WT | WT | WT |
| MISS | WT | WT | WT | MISS | WT | MISS | WT | WT | WT | WT | WT | WT |
| R337C | | | | R110P | | H178P | | | | | | |
| HER2p | HER2p | ERpHER2n | ERpHER2n | ERnHER2n | HER2p | ERpHER2n | ERpHER2n | ERpHER2n | ERpHER2n | HER2p | ERpHER2n | ERpHER2n |
| Died of Disease | Living | Died of Disease | Living | Living | Living | Living | Living | Living | Living | Died of Disease | Living | Died of Other Causes |
| 28.6 | 86.8 | 132.0333333 | 104.4 | 65.86666667 | 177.3333333 | 84.63333333 | 101.8333333 | 43.86666667 | 56.5 | 29.06666667 | 23.8 | 94.73333333 |
| NO | YES | NO | YES | YES | YES | NO | YES | YES | NO | NO | NO | NO |

| | | | | | | | | | |
|---|---|---|---|---|---|---|---|---|---|
| MB_0869 | MB_4633 | MB_4627 | MB_4004 | MB_4708 | MB_4618 | MB_4641 | MB_4622 | MB_4634 | MB_4688 |
| MUT | WT | WT | WT | WT | WT | WT | MUT | WT | WT |
| NULL | WT | WT | WT | WT | WT | WT | MISS | WT | WT |
| R213* | | | | | | | R248Q | | |
| ERnHER2n | ERpHER2n | ERpHER2n | ERpHER2n | ERpHER2n | ERpHER2n | ERpHER2n | ERnHER2n | ERpHER2n | HER2p |
| Died of Other Causes | Living | Died of Other Causes | Died of Disease | Died of Other Causes | Living | Died of Other Causes | Died of Other Causes | Died of Disease | Died of Disease |
| 112.4 | 318.2 | 186.6333333 | 256.8666667 | 227.9 | 267.4 | 148.5666667 | 282.8333333 | 109.6 | 85.3 |
| YES | NO | NO | NO | NO | NO | NO | YES | NO | NO |

| | | | | | | | | | | |
|---|---|---|---|---|---|---|---|---|---|---|
| MB_4665 | MB_4666 | MB_4640 | MB_4691 | MB_4671 | MB_4667 | MB_4721 | MB_4723 | MB_4718 | MB_4716 | MB_4729 |
| WT | MUT | MUT | MUT | WT | MUT | WT | WT | WT | WT | MUT |
| WT | MISS | MISS | NULL | WT | MISS | WT | WT | WT | WT | MISS |
| | R248W | I195T | P128Lfs*42 | | E271K | | | | | R158L |
| ERpHER2n | ERpHER2n | ERnHER2n | ERpHER2n | ERpHER2n | ERnHER2n | ERpHER2n | ERpHER2n | ERpHER2n | ERpHER2n | HER2p |
| Living | Died of Disease | Living | Died of Disease | Died of Disease | Died of Other Causes | Died of Disease | Living | Died of Disease | Died of Disease | Died of Other Causes |
| 260.2 | 111.5666667 | 200.6 | 282.5666667 | 34.7 | 295.3333333 | 23.73333333 | 211.9666667 | 198.3333333 | 76.13333333 | 28 |
| NO | NO | NO | NO | NO | NO | NO | NO | NO | NO | NO |

| | | | | | | | | | | |
|---|---|---|---|---|---|---|---|---|---|---|
| MB_4717 | MB_4724 | MB_4770 | MB_4752 | MB_4762 | MB_5169 | MB_4796 | MB_4790 | MB_4797 | MB_4800 | MB_5541 |
| MUT | MUT | MUT | MUT | WT | WT | WT | WT | WT | WT | WT |
| NULL | NULL | MISS | NULL | WT | WT | WT | WT | WT | WT | WT |
| C124* | I50Mfs*73 | R248W | X307_splice | | | | | | | |
| ERnHER2n | HER2p | HER2p | ERpHER2n | ERpHER2n | ERpHER2n | HER2p | ERpHER2n | ERpHER2n | ERpHER2n | ERpHER2n |
| Died of Disease | Living | Died of Other Causes | Died of Disease | Living | Died of Disease | Living | Died of Disease | Died of Other Causes | Died of Disease | Living |
| 42.9 | 146.0666667 | 152.0666667 | 125.2666667 | 252.1 | 51.96666667 | 128.2666667 | 70.46666667 | 99.36666667 | 110.9333333 | 200.1333333 |
| NO | NO | NO | NO | NO | NO | NO | NO | NO | NO | NO |

| | | | | | | | | | | |
|---|---|---|---|---|---|---|---|---|---|---|
| MB_5549 | MB_5519 | MB_5495 | MB_4832 | MB_4745 | MB_4825 | MB_4814 | MB_4757 | MB_4694 | MB_4698 | MB_4715 |
| MUT | WT | WT | WT | MUT | WT | MUT | MUT | MUT | WT | MUT |
| MISS | WT | WT | WT | MISS | WT | MISS | NULL | NULL | WT | MISS |
| D281H | | | | R248W | | R273P | E349* | X125_splice | | R273C |
| HER2p | ERpHER2n | ERpHER2n | ERpHER2n | HER2p | ERpHER2n | ERpHER2n | ERnHER2n | ERpHER2n | ERpHER2n | ERnHER2n |
| Died of Disease | Died of Other Causes | Living | Died of Disease | Died of Disease | Died of Other Causes | Died of Disease | Died of Disease | Living | Died of Disease | Died of Disease |
| 88.93333333 | 169.2333333 | 228.8 | 141.7333333 | 42.66666667 | 214.4333333 | 73.46666667 | 27.4 | 198.3 | 170.6666667 | 35.23333333 |
| YES | NO | NO | NO | YES | NO | NO | YES | NO | NO | YES |

| | | | | | | | | |
|---|---|---|---|---|---|---|---|---|
| MB_4685 | MB_4712 | MB_4672 | MB_4704 | MB_4655 | MB_4651 | MB_4661 | MB_4649 | MB_4642 |
| WT | MUT | WT | WT | MUT | MUT | WT | MUT | WT |
| WT | MISS | WT | WT | MISS | NULL | WT | NULL | WT |
| | C176Y | | | G244C | H193Qfs*53 | | Q331* | |
| ERpHER2n | ERpHER2n | ERpHER2n | ERpHER2n | ERpHER2n | ERpHER2n | ERpHER2n | ERpHER2n | ERpHER2n |
| Died of Other Causes | Died of Other Causes | Died of Other Causes | Died of Disease | Died of Other Causes | Died of Disease | Living | Died of Other Causes | Died of Disease |
| 43.3 | 249.5333333 | 220.9333333 | 64 | 99 | 30.3 | 234.2333333 | 16.83333333 | 285.7 |
| NO | NO | NO | NO | NO | NO | NO | NO | NO |

| | | | | | | | | |
|---|---|---|---|---|---|---|---|---|
| MB_4674 | MB_4648 | MB_5197 | MB_5152 | MB_5201 | MB_5189 | MB_5281 | MB_5212 | MB_5172 |
| WT | WT | WT | WT | WT | MUT | WT | WT | WT |
| WT | WT | WT | WT | WT | MISS | WT | WT | WT |
| | | | | | P278R | | | |
| ERpHER2n | ERpHER2n | ERpHER2n | ERpHER2n | ERpHER2n | ERpHER2n | ERnHER2n | HER2p | HER2p |
| Died of Disease | Died of Other Causes | Died of Disease | Died of Other Causes | Died of Other Causes | Died of Disease | Living | Died of Other Causes | Living |
| 81.06666667 | 168.5333333 | 223.8333333 | 115.9333333 | 96.96666667 | 111.9666667 | 79.3 | 179.1 | 184.2666667 |
| NO | NO | NO | NO | NO | NO | NO | NO | YES |

| | | | | | | | | | | |
|---|---|---|---|---|---|---|---|---|---|---|
| MB_5183 | MB_5185 | MB_5022 | MB_5014 | MB_4994 | MB_5017 | MB_4982 | MB_5004 | MB_4986 | MB_5327 | MB_5341 |
| WT | WT | WT | WT | WT | MUT | MUT | MUT | WT | MUT | WT |
| WT | WT | WT | WT | WT | MISS | NULL | NULL | WT | NULL | WT |
| | | | | | G262V | S166Hfs*4 | I255del | | C141* | |
| ERpHER2n | ERpHER2n | HER2p | ERpHER2n | ERpHER2n | ERpHER2n | ERnHER2n | ERpHER2n | ERpHER2n | HER2p | ERpHER2n |
| Died of Other Causes | Died of Disease | Died of Disease | Living | Died of Other Causes | Living | Died of Disease | Living | Died of Disease | Living | Living |
| 118.5333333 | 77.46666667 | 118.6 | 213.3666667 | 174.1333333 | 144.9333333 | 19.16666667 | 216.7333333 | 79.8 | 163.5333333 | 250.8333333 |
| NO | NO | NO | NO | NO | NO | NO | YES | NO | NO | NO |

| | | | | | | | | | |
|---|---|---|---|---|---|---|---|---|---|
| MB_5328 | MB_5318 | MB_5287 | MB_5324 | MB_4682 | MB_4710 | MB_4701 | MB_4709 | MB_4686 | MB_4706 |
| WT | MUT | WT | WT | WT | WT | WT | WT | WT | WT |
| WT | NULL | WT | WT | WT | WT | WT | WT | WT | WT |
| | C229Yfs*10 | | | | | | | | |
| ERpHER2n | ERpHER2n | ERpHER2n | ERpHER2n | ERpHER2n | ERpHER2n | ERpHER2n | ERpHER2n | ERpHER2n | ERpHER2n |
| Died of Other Causes | Died of Disease | Died of Other Causes | Died of Other Causes | Died of Disease | Living | Living | Living | Died of Disease | Died of Other Causes |
| 14.2 | 14.16666667 | 124 | 190.1 | 139.5333333 | 57.66666667 | 127.5333333 | 146.7666667 | 48.43333333 | 55.66666667 |
| NO | NO | NO | NO | NO | NO | NO | NO | NO | NO |

| | | | | | | | | | | |
|---|---|---|---|---|---|---|---|---|---|---|
| MB_4719 | MB_4702 | MB_4908 | MB_4871 | MB_4906 | MB_4911 | MB_4866 | MB_4858 | MB_4862 | MB_4872 | MB_4887 |
| MUT | WT | MUT | WT | WT | MUT | WT | MUT | MUT | WT | MUT |
| MISS | WT | NULL | WT | WT | MISS | WT | NULL | NULL | WT | NULL |
| E11Q | | Q317* | | | C238Y | | R342* | P390Lfs*32 | | I255Nfs*9 |
| ERpHER2n | ERpHER2n | HER2p | HER2p | ERpHER2n | ERnHER2n | ERpHER2n | HER2p | ERpHER2n | ERpHER2n | ERpHER2n |
| Died of Other Causes | Living | Died of Disease | Died of Disease | Died of Other Causes | Died of Disease | Living | Died of Other Causes | Living | Living | Died of Disease |
| 81.56666667 | 300.8666667 | 47.9 | 39.8 | 233.8666667 | 31.43333333 | 224.3 | 129.3333333 | 187.3 | 157.1 | 35.4 |
| NO | NO | NO | NO | NO | YES | NO | NO | NO | NO | NO |

| | | | | | | | | | | |
|---|---|---|---|---|---|---|---|---|---|---|
| MB_4867 | MB_4888 | MB_4929 | MB_4945 | MB_4930 | MB_4894 | MB_4898 | MB_4670 | MB_5013 | MB_4977 | MB_4967 |
| WT | MUT | MUT | MUT | MUT | MUT | WT | WT | WT | WT | WT |
| WT | NULL | MISS | MISS | MISS | MISS | WT | WT | WT | WT | WT |
| | E258* | E285K | R273C | F134C | H179Y | | | | | |
| ERpHER2n | ERnHER2n | HER2p | ERnHER2n | HER2p | ERpHER2n | ERpHER2n | ERpHER2n | ERpHER2n | ERpHER2n | ERpHER2n |
| Living | Living | Died of Disease | Died of Disease | Living | Living | Died of Disease | Died of Disease | Died of Other Causes | Died of Disease | Died of Other Causes |
| 221.9 | 230.5 | 40 | 20.13333333 | 58.43333333 | 232.4 | 117.9 | 67.8 | 251.2 | 216.7333333 | 130.7 |
| NO | NO | NO | YES | NO | NO | NO | NO | NO | NO | NO |

| | | | | | | | | | | |
|---|---|---|---|---|---|---|---|---|---|---|
| MB_4981 | MB_4003 | MB_4968 | MB_5052 | MB_5049 | MB_5041 | MB_5044 | MB_5072 | MB_4171 | MB_5053 | MB_5045 |
| WT | MUT | WT | MUT | WT | MUT | WT | MUT | WT | WT | MUT |
| WT | NULL | WT | NULL | WT | MISS | WT | MISS | WT | WT | NULL |
| | M246* | | S183* | | R337C | | N239I | | | A74Pfs*49 |
| ERpHER2n | ERpHER2n | ERpHER2n | ERnHER2n | ERpHER2n | ERnHER2n | ERpHER2n | ERnHER2n | ERpHER2n | ERpHER2n | ERpHER2n |
| Died of Disease | Died of Disease | Living | Died of Other Causes | Living | Living | Living | Died of Disease | Died of Disease | Died of Other Causes | Died of Other Causes |
| 180.7666667 | 106.5666667 | 78.7 | 34.56666667 | 255.3 | 173.9333333 | 139.4333333 | 50.23333333 | 184.7 | 150.6 | 168.3 |
| NO | NO | NO | NO | NO | NO | NO | YES | NO | NO | NO |

| | | | | | | | | | |
|---|---|---|---|---|---|---|---|---|---|
| MB_5116 | MB_5120 | MB_5074 | MB_5119 | MB_5114 | MB_4230 | MB_4154 | MB_4737 | MB_4764 | MB_4735 |
| WT | MUT | WT | MUT | MUT | WT | MUT | WT | WT | WT |
| WT | NULL | WT | MISS | MISS | WT | NULL | WT | WT | WT |
| | P223Afs*2 | | H193R | R273L | | V217Gfs*29 | | | |
| ERpHER2n | HER2p | ERpHER2n | ERpHER2n | HER2p | ERpHER2n | ERpHER2n | ERpHER2n | ERpHER2n | ERpHER2n |
| Died of Disease | Died of Disease | Died of Other Causes | Living | Died of Disease | Died of Other Causes | Died of Other Causes | Living | Living | Living |
| 71.76666667 | 27.8 | 221.6 | 59.76666667 | 15.36666667 | 115.6 | 49.56666667 | 122.2 | 245.5 | 297.8 |
| NO | YES | NO | NO | NO | NO | NO | NO | NO | NO |

| | | | | | | | | | | |
|---|---|---|---|---|---|---|---|---|---|---|
| MB_4730 | MB_4733 | MB_4758 | MB_5033 | MB_4741 | MB_4732 | MB_5305 | MB_5256 | MB_5273 | MB_5236 | MB_5238 |
| WT | MUT | MUT | WT | WT | MUT | WT | WT | WT | MUT | WT |
| WT | MISS | MISS | WT | WT | MISS | WT | WT | WT | NULL | WT |
| | G154S | R248W | | | I195T | | | | C242Afs*5 | |
| ERpHER2n | ERnHER2n | ERnHER2n | ERpHER2n | ERpHER2n | ERnHER2n | ERpHER2n | ERpHER2n | ERpHER2n | ERnHER2n | HER2p |
| Died of Other Causes | Died of Disease | Living | Died of Disease | Died of Disease | Living | Died of Disease | Died of Disease | Died of Disease | Living | Died of Disease |
| 131.3 | 19.73333333 | 211.9333333 | 52.73333333 | 56.5 | 149.8666667 | 149.4333333 | 108.0666667 | 60.86666667 | 166.6666667 | 44.13333333 |
| NO | YES | NO | NO | NO | YES | NO | NO | NO | YES | YES |

| | | | | | | | | |
|---|---|---|---|---|---|---|---|---|
| MB_5233 | MB_5244 | MB_5253 | MB_5260 | MB_4998 | MB_4993 | MB_5001 | MB_5084 | MB_4969 |
| WT | WT | WT | WT | WT | MUT | WT | WT | WT |
| WT | WT | WT | WT | WT | NULL | WT | WT | WT |
| | | | | | R174Sfs*67 | | | |
| ERpHER2n | ERpHER2n | ERpHER2n | ERpHER2n | ERpHER2n | ERnHER2n | ERpHER2n | ERpHER2n | ERpHER2n |
| Died of Other Causes | Died of Disease | Died of Other Causes | Died of Other Causes | Died of Disease | Died of Disease | Died of Disease | Died of Disease | Died of Other Causes |
| 209.0333333 | 216.8666667 | 102.0666667 | 202.1 | 65.56666667 | 75.23333333 | 81.8 | 198.1 | 198.1 |
| NO | NO | NO | NO | NO | YES | NO | NO | NO |

| | | | | | | | | | |
|---|---|---|---|---|---|---|---|---|---|
| MB_4999 | MB_5011 | MB_4959 | MB_4599 | MB_4616 | MB_4623 | MB_4644 | MB_4869 | MB_4878 | MB_4851 |
| WT | WT | WT | WT | WT | MUT | MUT | WT | WT | WT |
| WT | WT | WT | WT | WT | MISS | NULL | WT | WT | WT |
| | | | | | A276P | *394T | | | |
| ERpHER2n | ERpHER2n | ERpHER2n | ERpHER2n | ERpHER2n | ERpHER2n | HER2p | ERpHER2n | HER2p | ERpHER2n |
| Living | Died of Other Causes | Died of Disease | Living | Died of Disease | Died of Other Causes | Died of Disease | Died of Disease | Died of Disease | Died of Disease |
| 187.9333333 | 80.5 | 117.5666667 | 191.1666667 | 90.13333333 | 292.0333333 | 90 | 29.3 | 22.46666667 | 90.26666667 |
| NO | NO | NO | NO | NO | NO | NO | NO | NO | NO |

| | | | | | | | | | | |
|---|---|---|---|---|---|---|---|---|---|---|
| MB_4233 | MB_4937 | MB_4934 | MB_4899 | MB_4912 | MB_4935 | MB_4933 | MB_4900 | MB_4941 | MB_5221 | MB_5139 |
| MUT | WT | WT | WT | WT | WT | WT | MUT | WT | WT | WT |
| NULL | WT | WT | WT | WT | WT | WT | MISS | WT | WT | WT |
| N268Lfs*75 | | | | | | | A159V | | | |
| ERpHER2n | ERpHER2n | ERpHER2n | ERpHER2n | ERpHER2n | HER2p | ERpHER2n | ERpHER2n | ERpHER2n | ERpHER2n | ERpHER2n |
| Died of Disease | Died of Disease | Living | Living | Died of Disease | Living | Living | Living | Died of Disease | Died of Other Causes | Died of Other Causes |
| 36.36666667 | 77.23333333 | 70.23333333 | 275.7333333 | 50 | 194.2 | 272.9 | 224.6 | 58.63333333 | 131.3 | 89.96666667 |
| NO | NO | NO | NO | NO | YES | NO | NO | NO | NO | NO |

| | | | | | | | | | |
|---|---|---|---|---|---|---|---|---|---|
| MB_5222 | MB_5097 | MB_5338 | MB_5315 | MB_5195 | MB_5226 | MB_5232 | MB_5160 | MB_5126 | MB_5124 |
| WT | MUT | MUT | MUT | MUT | WT | MUT | WT | MUT | WT |
| WT | MISS | MISS | MISS | MISS | WT | NULL | WT | MISS | WT |
| | R248W | S215G | N239D | K132E | | S314* | | M237I | |
| ERnHER2n | ERpHER2n | ERpHER2n | HER2p | HER2p | ERpHER2n | ERnHER2n | ERpHER2n | ERnHER2n | ERpHER2n |
| Died of Other Causes | Living | Died of Disease | Died of Disease | Died of Disease | Living | Died of Other Causes | Died of Disease | Died of Disease | Died of Disease |
| 111.8333333 | 61.6 | 51.66666667 | 43.03333333 | 196.8666667 | 200.6 | 211.2 | 87.73333333 | 27.3 | 124.2 |
| NO | NO | NO | YES | NO | NO | NO | NO | YES | NO |

| | | | | | | | | |
|---|---|---|---|---|---|---|---|---|
| MB_4855 | MB_5279 | MB_4173 | MB_5145 | MB_4944 | MB_4961 | MB_5121 | MB_4956 | MB_4886 |
| MUT | WT | WT | MUT | WT | WT | WT | WT | MUT |
| MISS | WT | WT | NULL | WT | WT | WT | WT | NULL |
| V272L | | | S185Tfs*62 | | | | | S260Pfs*85 |
| ERpHER2n | ERpHER2n | ERpHER2n | ERnHER2n | ERpHER2n | ERpHER2n | ERpHER2n | ERpHER2n | HER2p |
| Died of Other Causes | Died of Other Causes | Died of Other Causes | Died of Disease | Died of Other Causes | Living | Died of Disease | Died of Disease | Died of Disease |
| 41.46666667 | 68.76666667 | 160.3333333 | 159.2333333 | 237.2666667 | 219.4666667 | 114.9 | 69.3 | 48.16666667 |
| NO | NO | NO | NO | NO | NO | NO | NO | YES |

| | | | | | | | | | | |
|---|---|---|---|---|---|---|---|---|---|---|
| MB_4950 | MB_4965 | MB_4962 | MB_4952 | MB_5267 | MB_5266 | MB_5396 | MB_4938 | MB_5351 | MB_5347 | MB_5312 |
| WT | WT | WT | MUT | WT | MUT | MUT | WT | WT | WT | WT |
| WT | WT | WT | MISS | WT | MISS | NULL | WT | WT | WT | WT |
| | | | G266V | | R248W | Q192* | | | | |
| ERpHER2n | ERpHER2n | ERpHER2n | HER2p | ERpHER2n | HER2p | ERpHER2n | ERnHER2n | HER2p | ERpHER2n | HER2p |
| Died of Disease | Died of Other Causes | Died of Disease | Living | Died of Disease | Died of Other Causes | Living | Died of Disease | Living | Died of Disease | Died of Disease |
| 150.6 | 263.6 | 197.8333333 | 184.4 | 71.06666667 | 83.63333333 | 165.6666667 | 70.6 | 202.1 | 211.9 | 37.9 |
| NO | NO | NO | NO | NO | NO | NO | YES | NO | NO | YES |

| | | | | | | | | | | | |
|---|---|---|---|---|---|---|---|---|---|---|---|
| MB_5311 | MB_5284 | MB_4148 | MB_5078 | MB_5088 | MB_5064 | MB_5107 | MB_5066 | MB_5068 | MB_5061 | MB_5070 | MB_5059 |
| WT | WT | WT | MUT | WT | WT | WT | WT | WT | WT | WT | WT |
| WT | WT | WT | NULL | WT | WT | WT | WT | WT | WT | WT | WT |
| | | | X187_splice | | | | | | | | |
| ERpHER2n | ERpHER2n | ERpHER2n | HER2p | ERpHER2n | ERpHER2n | ERpHER2n | ERpHER2n | ERpHER2n | ERpHER2n | ERnHER2n | ERpHER2n |
| Died of Disease | Died of Other Causes | Died of Disease | Died of Disease | Died of Other Causes | Living | Died of Disease | Living | Living | Living | Living | Living |
| 57.66666667 | 65.33333333 | 63.03333333 | 87.1 | 199.3666667 | 91.5 | 82.63333333 | 94.7 | 182.5 | 39.43333333 | 213.9 | 257.6333333 |
| NO | NO | NO | NO | NO | NO | NO | NO | NO | NO | YES | NO |

| | | | | | | | | | | |
|---|---|---|---|---|---|---|---|---|---|---|
| MB_4266 | MB_4276 | MB_4771 | MB_4739 | MB_5331 | MB_4743 | MB_4785 | MB_4778 | MB_4763 | MB_4779 | MB_4849 |
| WT | MUT | WT | WT | WT | WT | WT | WT | MUT | WT | WT |
| WT | NULL | WT | WT | WT | WT | WT | WT | MISS | WT | WT |
| | H178Pfs*2 | | | | | | | R248Q | | |
| ERpHER2n | HER2p | ERpHER2n | ERpHER2n | HER2p | ERpHER2n | ERpHER2n | ERpHER2n | HER2p | ERpHER2n | ERpHER2n |
| Died of Disease | Died of Disease | Living | Died of Other Causes | Died of Disease | Living | Died of Disease | Died of Other Causes | Living | Living | Died of Disease |
| 33.9 | 132.2 | 297.2333333 | 161.6666667 | 124.1333333 | 230.1666667 | 121.6666667 | 206.1333333 | 102.1 | 101.5666667 | 45.33333333 |
| NO | NO | NO | NO | NO | NO | NO | NO | YES | NO | NO |

| | | | | | | | | | |
|---|---|---|---|---|---|---|---|---|---|
| MB_4787 | MB_4767 | MB_4784 | MB_5043 | MB_4992 | MB_5020 | MB_5019 | MB_5015 | MB_5048 | MB_5035 |
| WT | WT | WT | WT | WT | WT | MUT | WT | WT | WT |
| WT | WT | WT | WT | WT | WT | NULL | WT | WT | WT |
| | | | | | | S183* | | | |
| ERpHER2n | ERpHER2n | ERpHER2n | ERpHER2n | ERnHER2n | ERpHER2n | ERnHER2n | ERpHER2n | ERpHER2n | ERpHER2n |
| Died of Other Causes | Died of Disease | Died of Other Causes | Died of Disease | Living | Died of Disease | Died of Disease | Died of Other Causes | Died of Disease | Died of Disease |
| 81.06666667 | 47.13333333 | 145.7333333 | 128.3666667 | 165.3666667 | 84.83333333 | 37.8 | 252.3 | 86.36666667 | 213.2 |
| NO | NO | NO | NO | NO | NO | NO | NO | NO | NO |

| | | | | | | | | | | |
|---|---|---|---|---|---|---|---|---|---|---|
| MB_5398 | MB_5377 | MB_5291 | MB_5403 | MB_5224 | MB_5369 | MB_5404 | MB_5397 | MB_5378 | MB_5388 | MB_5395 |
| WT | WT | WT | WT | WT | WT | WT | MUT | MUT | WT | WT |
| WT | WT | WT | WT | WT | WT | WT | MISS | MISS | WT | WT |
| | | | | | | | Y163C | R248Q | | |
| ERpHER2n | ERpHER2n | ERpHER2n | ERpHER2n | ERpHER2n | ERpHER2n | ERpHER2n | ERpHER2n | ERpHER2n | ERpHER2n | ERpHER2n |
| Living | Died of Disease | Living | Died of Other Causes | Died of Disease | Died of Disease | Died of Disease | Died of Disease | Died of Disease | Died of Other Causes | Living |
| 213 | 102 | 153.5333333 | 95.73333333 | 99.4 | 39.3 | 74.73333333 | 172.8666667 | 12.26666667 | 146.9333333 | 165.1666667 |
| NO | YES | NO | NO | NO | NO | NO | NO | YES | NO | NO |

| | | | | | | | | | |
|---|---|---|---|---|---|---|---|---|---|
| MB_5365 | MB_5389 | MB_5393 | MB_5361 | MB_5392 | MB_5147 | MB_4140 | MB_5138 | MB_5123 | MB_5065 |
| WT | WT | WT | MUT | MUT | WT | WT | MUT | WT | MUT |
| WT | WT | WT | MISS | MISS | WT | WT | MISS | WT | MISS |
| | | | L194R | M246V | | | R273H | | C275F |
| ERpHER2n | ERpHER2n | ERpHER2n | ERpHER2n | ERnHER2n | HER2p | ERpHER2n | ERnHER2n | ERpHER2n | ERnHER2n |
| Died of Disease | Died of Other Causes | Died of Other Causes | Died of Disease | Living | Living | Died of Other Causes | Died of Disease | Died of Disease | Living |
| 23.03333333 | 124.7666667 | 154 | 15.36666667 | 237.2666667 | 20.26666667 | 114.5 | 37 | 90.3 | 184.8 |
| NO | NO | NO | NO | NO | NO | NO | YES | NO | NO |

| | | | | | | | | | |
|---|---|---|---|---|---|---|---|---|---|
| MB_4127 | MB_5058 | MB_5050 | MB_5027 | MB_5060 | MB_5062 | MB_5383 | MB_5101 | MB_5258 | MB_5268 |
| MUT | MUT | WT | WT | WT | WT | WT | MUT | MUT | WT |
| NULL | MISS | WT | WT | WT | WT | WT | MISS | MISS | WT |
| H214_E221del | L194P | | | | | | F134V | M246R | |
| HER2p | ERnHER2n | ERpHER2n | ERpHER2n | ERpHER2n | HER2p | ERpHER2n | ERpHER2n | ERnHER2n | ERpHER2n |
| Died of Other Causes | Died of Disease | Living | Living | Died of Other Causes | Living | Died of Disease | Died of Disease | Died of Other Causes | Living |
| 258.1666667 | 178.7333333 | 184.3333333 | 189.8666667 | 150.1 | 168.2666667 | 113.6666667 | 34.33333333 | 101.0666667 | 186.1333333 |
| YES | YES | NO | NO | NO | NO | NO | NO | NO | NO |

| | | | | | | | | | | |
|---|---|---|---|---|---|---|---|---|---|---|
| MB_5261 | MB_5272 | MB_5264 | MB_5259 | MB_5306 | MB_5270 | MB_5411 | MB_5490 | MB_5421 | MB_5348 | MB_5429 |
| WT | MUT | WT | MUT | WT | WT | MUT | WT | MUT | MUT | WT |
| WT | MISS | WT | MISS | WT | WT | MISS | WT | MISS | NULL | WT |
| | H193Y | | R175H | | | H179L | | C242Y | I195Sfs*52 | |
| ERpHER2n | ERpHER2n | ERpHER2n | HER2p | ERpHER2n | ERpHER2n | HER2p | ERpHER2n | ERnHER2n | ERnHER2n | ERpHER2n |
| Died of Other Causes | Died of Disease | Living | Died of Disease | Living | Died of Other Causes | Died of Disease | Died of Disease | Living | Living | Died of Other Causes |
| 116.4666667 | 201.9333333 | 199.3 | 46.16666667 | 159.7333333 | 187.8333333 | 26 | 51.2 | 194.5666667 | 169.6666667 | 30.7 |
| NO | NO | NO | YES | NO | NO | YES | NO | NO | YES | NO |

| | | | | | | | | | | |
|---|---|---|---|---|---|---|---|---|---|---|
| MB_5405 | MB_5414 | MB_5489 | MB_5417 | MB_5481 | MB_5409 | MB_5299 | MB_5408 | MB_5475 | MB_5505 | MB_5100 |
| WT | MUT | WT | MUT | MUT | MUT | MUT | WT | WT | WT | MUT |
| WT | MISS | WT | NULL | NULL | MISS | MISS | WT | WT | WT | NULL |
| | H193L | | F54Kfs*67 | E298* | K132E | C176S | | | | R306* |
| ERpHER2n | HER2p | ERpHER2n | HER2p | ERpHER2n | HER2p | ERnHER2n | ERnHER2n | ERpHER2n | ERpHER2n | ERnHER2n |
| Living | Died of Other Causes | Living | Living | Died of Disease | Living | Died of Disease | Living | Died of Other Causes | Died of Disease | Died of Disease |
| 117.6 | 140.6 | 216.9666667 | 107.0666667 | 46.36666667 | 198.1333333 | 16.7 | 101.4 | 2.533333333 | 164.6 | 19.9 |
| NO | NO | NO | YES | NO | YES | YES | NO | NO | NO | YES |

| | | | | | | | | | |
|---|---|---|---|---|---|---|---|---|---|
| MB_5502 | MB_5485 | MB_4293 | MB_4289 | MB_5293 | MB_5381 | MB_4834 | MB_5498 | MB_4749 | MB_4818 |
| MUT | WT | WT | WT | WT | MUT | WT | MUT | WT | WT |
| MISS | WT | WT | WT | WT | NULL | WT | NULL | WT | WT |
| A159D | | | | | E198* | | V197Gfs*50 | | |
| ERpHER2n | ERpHER2n | ERpHER2n | HER2p | ERpHER2n | HER2p | ERpHER2n | ERpHER2n | ERpHER2n | ERpHER2n |
| Living | Died of Other Causes | Died of Other Causes | Died of Disease | Died of Disease | Died of Disease | Died of Disease | Living | Died of Other Causes | Died of Other Causes |
| 189.7333333 | 31.8 | 219.1666667 | 96.83333333 | 208.9666667 | 201.8333333 | 36.16666667 | 225.1666667 | 51.46666667 | 76.23333333 |
| NO | NO | NO | NO | NO | YES | NO | NO | NO | NO |

| | | | | | | | | | | |
|---|---|---|---|---|---|---|---|---|---|---|
| MB_4805 | MB_4809 | MB_5418 | MB_5441 | MB_5424 | MB_5296 | MB_5446 | MB_5093 | MB_5350 | MB_5230 | MB_5206 |
| WT | MUT | WT | MUT | WT | MUT | WT | WT | WT | WT | WT |
| WT | NULL | WT | MISS | WT | NULL | WT | WT | WT | WT | WT |
| | Q144* | | C135F | | R213* | | | | | |
| ERpHER2n | ERpHER2n | HER2p | HER2p | ERpHER2n | HER2p | ERnHER2n | ERpHER2n | ERpHER2n | ERpHER2n | ERpHER2n |
| Died of Disease | Living | Living | Died of Other Causes | Died of Disease | Died of Disease | Died of Disease | Died of Other Causes | Died of Disease | Living | Living |
| 129.8 | 286.0666667 | 207.1666667 | 9.7 | 83.53333333 | 13.8 | 162.8333333 | 142.4333333 | 53.63333333 | 176.5 | 205.7333333 |
| NO | NO | NO | NO | NO | NO | YES | NO | NO | NO | NO |

| | | | | | | | | | |
|---|---|---|---|---|---|---|---|---|---|
| MB_4687 | MB_5433 | MB_5442 | MB_5427 | MB_5491 | MB_5493 | MB_5451 | MB_5454 | MB_5360 | MB_5229 |
| MUT | WT | MUT | MUT | WT | MUT | WT | MUT | WT | MUT |
| NULL | WT | MISS | MISS | WT | NULL | WT | MISS | WT | MISS |
| P318Afs*19 | | R249G | K120E | | G262Efs*9 | | R282W | | L111Q |
| ERpHER2n | ERpHER2n | ERnHER2n | ERnHER2n | ERpHER2n | ERpHER2n | ERpHER2n | ERpHER2n | ERpHER2n | HER2p |
| Died of Other Causes | Died of Disease | Living | Died of Other Causes | Died of Disease | Died of Disease | Living | Died of Disease | Died of Other Causes | Died of Disease |
| 74.1 | 176.3666667 | 205.8 | 155.7333333 | 65.83333333 | 57.23333333 | 189.9 | 50.66666667 | 161.1333333 | 33.96666667 |
| NO | NO | YES | NO | NO | NO | NO | NO | NO | YES |

| | | | | | | | |
|---|---|---|---|---|---|---|---|
| MB_4827 | MB_5455 | MB_5370 | MB_5290 | MB_5366 | MB_5382 | MB_5402 | MB_5288 |
| WT | WT | WT | WT | WT | WT | WT | MUT |
| WT | WT | WT | WT | WT | WT | WT | MISS |
| | | | | | | | E285K |
| ERpHER2n | ERpHER2n | ERpHER2n | ERpHER2n | ERpHER2n | ERpHER2n | ERpHER2n | ERpHER2n |
| Died of Other Causes | Died of Other Causes | Died of Other Causes | Died of Other Causes | Living | Died of Other Causes | Died of Other Causes | Died of Other Causes |
| 45.7 | 61.8 | 119.3 | 36.93333333 | 243.1666667 | 116.2333333 | 107.7666667 | 128.4 |
| NO | NO | NO | NO | NO | NO | NO | NO |

| | | | | | | | | | | |
|---|---|---|---|---|---|---|---|---|---|---|
| MB_5384 | MB_5358 | MB_4357 | MB_4991 | MB_5401 | MB_5390 | MB_5412 | MB_5193 | MB_5182 | MB_5157 | MB_5205 |
| WT | WT | MUT | WT | WT | MUT | WT | WT | WT | MUT | MUT |
| WT | WT | MISS | WT | WT | NULL | WT | WT | WT | MISS | MISS |
| | | K132N | | | L111Afs*11 | | | | V173L | K132E |
| ERpHER2n | ERpHER2n | ERpHER2n | ERpHER2n | ERpHER2n | ERnHER2n | ERpHER2n | ERpHER2n | ERpHER2n | ERpHER2n | ERpHER2n |
| Died of Other Causes | Died of Disease | Died of Disease | Living | Living | Died of Disease | Living | Died of Other Causes | Died of Other Causes | Died of Disease | Living |
| 71.16666667 | 28.83333333 | 114.0333333 | 126.8666667 | 70.16666667 | 30.93333333 | 220.2333333 | 54.26666667 | 134.2666667 | 18.23333333 | 185.3 |
| NO | NO | NO | NO | NO | YES | NO | NO | NO | NO | YES |

| | | | | | | | | | |
|---|---|---|---|---|---|---|---|---|---|
| MB_5161 | MB_5292 | MB_5211 | MB_5218 | MB_5209 | MB_5227 | MB_5213 | MB_5166 | MB_5163 | MB_5188 |
| WT | MUT | WT | WT | MUT | WT | MUT | MUT | WT | WT |
| WT | MISS | WT | WT | MISS | WT | MISS | MISS | WT | WT |
| | R175H | | | R110P | | N131I | R280K | | |
| ERpHER2n | HER2p | ERpHER2n | ERpHER2n | ERnHER2n | ERpHER2n | ERnHER2n | HER2p | ERpHER2n | ERpHER2n |
| Living | Died of Disease | Died of Other Causes | Living | Died of Other Causes | Died of Other Causes | Died of Disease | Died of Disease | Died of Other Causes | Died of Disease |
| 175.3333333 | 48.43333333 | 247.8333333 | 254.9666667 | 175.1 | 187.0333333 | 43.4 | 80.73333333 | 212.2 | 19.4 |
| NO | NO | NO | NO | NO | NO | YES | NO | NO | NO |

| | | | | | | | | |
|---|---|---|---|---|---|---|---|---|
| MB_5143 | MB_5329 | MB_5322 | MB_5310 | MB_4000 | MB_5240 | MB_5228 | MB_5117 | MB_5176 |
| WT | MUT | WT | WT | WT | WT | MUT | WT | WT |
| WT | MISS | WT | WT | WT | WT | NULL | WT | WT |
| | R280T | | | | | X225_splice | | |
| ERpHER2n | ERpHER2n | ERpHER2n | ERpHER2n | ERpHER2n | ERpHER2n | ERpHER2n | ERpHER2n | ERpHER2n |
| Died of Other Causes | Died of Disease | Died of Disease | Died of Other Causes | Died of Other Causes | Died of Other Causes | Died of Other Causes | Living | Died of Other Causes |
| 127.9333333 | 45.73333333 | 102.3 | 153.3 | 28.06666667 | 103.8333333 | 180.7333333 | 209.2666667 | 112.6333333 |
| NO | YES | NO | NO | NO | NO | NO | NO | NO |

| | | | | | | | | | |
|---|---|---|---|---|---|---|---|---|---|
| MB_5215 | MB_5223 | MB_5179 | MB_5330 | MB_5144 | MB_5196 | MB_5200 | MB_5583 | MB_5601 | MB_5632 |
| WT | MUT | MUT | WT | WT | WT | WT | WT | WT | WT |
| WT | MISS | NULL | WT | WT | WT | WT | WT | WT | WT |
| | Y234C | R213* | | | | | | | |
| ERpHER2n | ERnHER2n | ERpHER2n | ERpHER2n | ERpHER2n | ERpHER2n | ERpHER2n | ERpHER2n | ERpHER2n | ERpHER2n |
| Died of Disease | Died of Disease | Died of Other Causes | Died of Other Causes | Living | Died of Disease | Died of Disease | Living | Died of Other Causes | Died of Disease |
| 118.7 | 144.6666667 | 246.6 | 114.5333333 | 213.0333333 | 94.03333333 | 52.3 | 111.3666667 | 214.8 | 38.16666667 |
| NO | YES | NO | NO | NO | NO | NO | NO | NO | NO |

| | | | | | | | | | | |
|---|---|---|---|---|---|---|---|---|---|---|
| MB_5616 | MB_5604 | MB_5589 | MB_5582 | MB_5625 | MB_5571 | MB_5596 | MB_5590 | MB_5599 | MB_5629 | MB_5579 |
| MUT | WT | WT | WT | MUT | WT | WT | MUT | WT | WT | WT |
| MISS | WT | WT | WT | NULL | WT | WT | MISS | WT | WT | WT |
| Y220C | | | | p.E68X | | | S127F | | | |
| ERnHER2n | ERpHER2n | ERpHER2n | ERpHER2n | ERnHER2n | ERpHER2n | ERpHER2n | ERpHER2n | ERpHER2n | ERpHER2n | ERpHER2n |
| Living | Living | Living | Living | Died of Disease | Died of Disease | Died of Disease | Living | Living | Died of Other Causes | Died of Disease |
| 180.6333333 | 145.6333333 | 185.1333333 | 222.3333333 | 32.3 | 153.8666667 | 167.9333333 | 17.83333333 | 224.8666667 | 103.8 | 90.6 |
| YES | NO | NO | NO | YES | NO | NO | NO | NO | NO | NO |

| | | | | | | | |
|---|---|---|---|---|---|---|---|
| MB_5623 | MB_5654 | MB_5617 | MB_5638 | MB_5635 | MB_5642 | MB_5647 | MB_4018 |
| WT | WT | WT | WT | WT | WT | WT | WT |
| WT | WT | WT | WT | WT | WT | WT | WT |
| ERpHER2n | ERpHER2n | ERpHER2n | ERpHER2n | ERpHER2n | ERpHER2n | ERpHER2n | ERpHER2n |
| Died of Other Causes | Died of Other Causes | Died of Other Causes | Died of Other Causes | Died of Disease | Died of Other Causes | Died of Other Causes | Died of Disease |
| 137.8 | 173.0333333 | 128.2 | 195.7 | 21.06666667 | 156.8 | 30.8 | 105.6666667 |
| NO | NO | NO | NO | NO | NO | NO | NO |

| | | | | | | | | | |
|---|---|---|---|---|---|---|---|---|---|
| MB_5584 | MB_5626 | MB_5602 | MB_5653 | MB_5576 | MB_5575 | MB_5646 | MB_5634 | MB_5591 | MB_5567 |
| MUT | WT | MUT | WT | WT | WT | WT | MUT | WT | WT |
| NULL | WT | NULL | WT | WT | WT | WT | NULL | WT | WT |
| P301Qfs*44 | | W91* | | | | | N239* | | |
| ERpHER2n | ERpHER2n | ERnHER2n | ERpHER2n | ERpHER2n | ERpHER2n | ERpHER2n | ERnHER2n | ERpHER2n | ERpHER2n |
| Died of Other Causes | Living | Living | Living | Living | Died of Other Causes | Died of Other Causes | Died of Disease | Died of Other Causes | Died of Other Causes |
| 91.1 | 82.96666667 | 224.4333333 | 194.6 | 194 | 117.6666667 | 167.1 | 62.9 | 155.3666667 | 178.5666667 |
| NO | NO | YES | NO | NO | NO | NO | NO | NO | NO |

| | | | | | | | | | |
|---|---|---|---|---|---|---|---|---|---|
| MB_5651 | MB_5603 | MB_5622 | MB_5597 | MB_5613 | MB_5580 | MB_5645 | MB_5548 | MB_5526 | MB_5531 |
| MUT | WT | WT | WT | MUT | WT | WT | MUT | MUT | MUT |
| MISS | WT | WT | WT | NULL | WT | WT | MISS | MISS | NULL |
| I195T | | | | Y205_T211del | | | P278R | R273C | R333Vfs*12 |
| ERnHER2n | ERpHER2n | ERpHER2n | ERpHER2n | ERpHER2n | ERpHER2n | ERpHER2n | ERnHER2n | ERnHER2n | ERnHER2n |
| Died of Disease | Died of Other Causes | Living | Died of Other Causes | Died of Other Causes | Died of Other Causes | Died of Other Causes | Living | Living | Living |
| 21 | 171.6333333 | 182.9333333 | 32.73333333 | 212.3666667 | 71.63333333 | 22.23333333 | 25.33333333 | 151.0666667 | 183.9666667 |
| YES | NO | NO | NO | NO | NO | NO | NO | NO | NO |

| | | | | | | | | | | | |
|---|---|---|---|---|---|---|---|---|---|---|---|
| MB_5468 | MB_5474 | MB_5514 | MB_5302 | MB_5510 | MB_5473 | MB_5484 | MB_5432 | MB_5525 | MB_5562 | MB_5497 | MB_5551 |
| MUT | MUT | WT | WT | WT | WT | WT | WT | MUT | WT | WT | MUT |
| NULL | MISS | WT | WT | WT | WT | WT | WT | MISS | WT | WT | NULL |
| S106Rfs*41 | L111P | | | | | | | R248Q | | | R213* |
| HER2p | HER2p | ERpHER2n | ERpHER2n | ERpHER2n | ERpHER2n | ERpHER2n | ERpHER2n | ERpHER2n | ERpHER2n | ERpHER2n | ERnHER2n |
| Living | Living | Living | Living | Died of Disease | Died of Other Causes | Living | Died of Other Causes | Living | Died of Other Causes | Living | Living |
| 99.36666667 | 226.7 | 182.9 | 191.1 | 103.1 | 125.9 | 238.5 | 98.5 | 2 | 102.9666667 | 237.5 | 182.2333333 |
| YES | YES | NO | NO | NO | NO | NO | NO | NO | NO | NO | NO |

| | | | | | | | | | | |
|---|---|---|---|---|---|---|---|---|---|---|
| MB_5300 | MB_5560 | MB_5550 | MB_5520 | MB_5529 | MB_5554 | MB_5301 | MB_4421 | MB_4408 | MB_4146 | MB_5534 |
| WT | MUT | WT | MUT | MUT | WT | WT | WT | MUT | WT | WT |
| WT | MISS | WT | MISS | NULL | WT | WT | WT | NULL | WT | WT |
| | R175H | | C141Y | C275Lfs*70 | | | | R196* | | |
| ERpHER2n | ERnHER2n | ERpHER2n | ERpHER2n | ERnHER2n | ERpHER2n | ERpHER2n | ERpHER2n | ERnHER2n | ERnHER2n | ERpHER2n |
| Living | Living | Died of Other Causes | Died of Disease | Died of Disease | Living | Living | Died of Other Causes | Died of Other Causes | Died of Disease | Died of Disease |
| 190.1666667 | 186.6 | 171.3 | 16.3 | 14.8 | 229.8333333 | 241.2333333 | 264.7666667 | 206.5666667 | 15.63333333 | 79.1 |
| NO | NO | NO | NO | YES | NO | NO | NO | NO | YES | NO |

| | | | | | | | | | | | |
|---|---|---|---|---|---|---|---|---|---|---|---|
| MB_5521 | MB_5482 | MB_5483 | MB_5532 | MB_5556 | MB_5518 | MB_5486 | MB_5511 | MB_5422 | MB_5535 | MB_5477 | MB_5472 |
| WT | MUT | MUT | WT | WT | MUT | WT | MUT | WT | MUT | WT | WT |
| WT | MISS | MISS | WT | WT | NULL | WT | MISS | WT | NULL | WT | WT |
| | R337C | R248W | | | G374Afs*43 | | C238Y | | X261_splice | | |
| ERpHER2n | ERpHER2n | HER2p | ERpHER2n | ERpHER2n | ERpHER2n | ERpHER2n | ERnHER2n | ERpHER2n | HER2p | ERpHER2n | ERpHER2n |
| Living | Died of Disease | Died of Disease | Died of Disease | Living | Died of Disease | Died of Disease | Living | Died of Disease | Died of Disease | Living | Died of Disease |
| 124.8 | 17.66666667 | 42.43333333 | 79.86666667 | 225.4 | 30.86666667 | 123.3 | 158.9666667 | 46.06666667 | 120.1333333 | 110.4666667 | 19.56666667 |
| NO | YES | YES | NO | NO | NO | NO | NO | NO | YES | NO | NO |

| | | | | | | | | |
|---|---|---|---|---|---|---|---|---|
| MB_5540 | MB_5184 | MB_5294 | MB_5565 | MB_5040 | MB_4801 | MB_5499 | MB_5459 | MB_5243 |
| MUT | WT | MUT | MUT | WT | WT | WT | WT | MUT |
| MISS | WT | MISS | NULL | WT | WT | WT | WT | NULL |
| P278S | | R175H | N200Ifs*47 | | | | | G262_N263del |
| HER2p | ERpHER2n | ERnHER2n | ERnHER2n | ERpHER2n | ERpHER2n | ERpHER2n | HER2p | ERpHER2n |
| Died of Disease | Died of Other Causes | Living | Died of Other Causes | Died of Other Causes | Died of Other Causes | Living | Died of Disease | Died of Other Causes |
| 48.43333333 | 89.33333333 | 195.9333333 | 194.1 | 79.16666667 | 130.3666667 | 123.7333333 | 90.8 | 38.43333333 |
| NO | NO | YES | YES | NO | NO | NO | NO | NO |

| | | | | | | | | |
|---|---|---|---|---|---|---|---|---|
| MB_4120 | MB_4966 | MB_5457 | MB_4011 | MB_5478 | MB_4822 | MB_4017 | MB_5018 | MB_4791 |
| WT | WT | WT | MUT | WT | WT | WT | WT | MUT |
| WT | WT | WT | NULL | WT | WT | WT | WT | MISS |
| | | | X261_splice | | | | | E285K |
| ERpHER2n | ERpHER2n | ERpHER2n | ERpHER2n | ERpHER2n | ERpHER2n | ERpHER2n | ERpHER2n | ERpHER2n |
| Died of Other Causes | Died of Disease | Died of Disease | Living | Died of Other Causes | Died of Disease | Died of Other Causes | Died of Other Causes | Died of Other Causes |
| 217.1 | 49.3 | 164.5666667 | 193.1666667 | 34.3 | 91.63333333 | 139.3 | 108.6 | 173.8333333 |
| NO | NO | NO | NO | NO | NO | NO | NO | NO |

| | | | | | | | | | | |
|---|---|---|---|---|---|---|---|---|---|---|
| MB_5118 | MB_4806 | MB_4970 | MB_2964 | MB_2963 | MB_2957 | MB_2954 | MB_2916 | MB_2725 | MB_2711 | MB_2730 |
| WT | WT | WT | MUT | MUT | MUT | WT | WT | WT | WT | WT |
| WT | WT | WT | MISS | MISS | MISS | WT | WT | WT | WT | WT |
| | | | R248W | R267G | S241C | | | | | |
| ERpHER2n | ERpHER2n | ERpHER2n | ERpHER2n | ERnHER2n | ERnHER2n | ERpHER2n | ERpHER2n | ERpHER2n | ERpHER2n | ERpHER2n |
| Living | Died of Other Causes | Died of Disease | Died of Disease | Died of Disease | Living | Died of Disease | Living | Living | Living | Living |
| 211.7333333 | 201.7666667 | 88.8 | 9.6 | 227.8333333 | 262.1333333 | 5.5 | 266.9333333 | 236.9333333 | 201.4666667 | 133.7333333 |
| NO | NO | NO | NO | NO | YES | NO | NO | NO | NO | NO |

| | | | | | | | | | | | |
|---|---|---|---|---|---|---|---|---|---|---|---|
| MB_2669 | MB_2705 | MB_2728 | MB_2708 | MB_2735 | MB_3035 | MB_3064 | MB_3060 | MB_3049 | MB_3037 | MB_3063 | MB_3083 |
| WT | WT | WT | MUT | WT | WT | WT | WT | WT | WT | MUT | WT |
| WT | WT | WT | MISS | WT | WT | WT | WT | WT | WT | NULL | WT |
| | | | T155N | | | | | | | R213* | |
| ERpHER2n | ERpHER2n | ERpHER2n | ERpHER2n | ERpHER2n | ERpHER2n | ERpHER2n | ERpHER2n | ERpHER2n | ERpHER2n | ERnHER2n | ERpHER2n |
| Died of Disease | Living | Died of Disease | Died of Disease | Living | Living | Died of Disease | Died of Disease | Living | Living | Died of Disease | Died of Disease |
| 73.7 | 163.7333333 | 51.4 | 133.2333333 | 274.5 | 260.7333333 | 165.1666667 | 44.6 | 259.9666667 | 98.83333333 | 28.56666667 | 81.1 |
| NO | NO | YES | NO | YES | NO | NO | NO | NO | NO | YES | NO |

| | | | | | | | | | | | |
|---|---|---|---|---|---|---|---|---|---|---|---|
| MB_3046 | MB_3067 | MB_3351 | MB_3367 | MB_3361 | MB_3412 | MB_3371 | MB_3429 | MB_3360 | MB_3403 | MB_3350 | MB_3355 |
| MUT | MUT | WT | MUT | MUT | WT | WT | WT | MUT | WT | WT | MUT |
| NULL | MISS | WT | NULL | NULL | WT | WT | WT | MISS | WT | WT | NULL |
| L111Wfs*12 | R175H | | P191Lfs*53 | R333Vfs*12 | | | | P250L | | | X187_splice |
| ERnHER2n | ERnHER2n | ERpHER2n | ERpHER2n | HER2p | ERpHER2n | ERpHER2n | ERpHER2n | HER2p | ERpHER2n | ERpHER2n | HER2p |
| Died of Other Causes | Living | Living | Living | Died of Disease | Living | Living | Living | Died of Disease | Living | Died of Other Causes | Living |
| 73.83333333 | 259.5333333 | 241.8666667 | 179.4333333 | 68.2 | 228.8 | 241.2666667 | 243.5 | 26.76666667 | 138.3333333 | 227.8 | 109 |
| NO | YES | NO | NO | NO | NO | NO | NO | NO | NO | NO | YES |

| | | | | | | | | | |
|---|---|---|---|---|---|---|---|---|---|
| MB_3329 | MB_3467 | MB_3341 | MB_3298 | MB_3357 | MB_3388 | MB_3365 | MB_2749 | MB_2754 | MB_2752 |
| MUT | MUT | WT | WT | MUT | MUT | WT | WT | WT | WT |
| MISS | NULL | WT | WT | MISS | MISS | WT | WT | WT | WT |
| G245V | C229Yfs*10 | | | G245S | R249M | | | | |
| HER2p | HER2p | ERpHER2n | ERpHER2n | ERpHER2n | ERpHER2n | ERpHER2n | ERpHER2n | ERnHER2n | ERpHER2n |
| Died of Disease | Died of Disease | Died of Disease | Living | Died of Disease | Died of Other Causes | Living | Died of Other Causes | Died of Disease | Died of Disease |
| 37.96666667 | 38.8 | 148.7666667 | 242.5666667 | 38.03333333 | 228.6 | 241.6 | 97.76666667 | 203.5333333 | 81.93333333 |
| NO | YES | NO | NO | NO | NO | NO | NO | NO | NO |

| | | | | | | | | | | |
|---|---|---|---|---|---|---|---|---|---|---|
| MB_2745 | MB_2747 | MB_2760 | MB_2742 | MB_2750 | MB_2753 | MB_2966 | MB_2953 | MB_2969 | MB_3014 | MB_2947 |
| WT | WT | WT | MUT | WT | MUT | WT | MUT | WT | MUT | WT |
| WT | WT | WT | MISS | WT | NULL | WT | MISS | WT | NULL | WT |
| | | | L111P | | K139Nfs*9 | | Y234C | | E339Gfs*6 | |
| ERpHER2n | ERpHER2n | ERpHER2n | HER2p | ERpHER2n | ERnHER2n | ERpHER2n | ERpHER2n | ERpHER2n | ERnHER2n | ERpHER2n |
| Died of Disease | Died of Other Causes | Died of Disease | Died of Disease | Died of Other Causes | Living | Living | Died of Disease | Died of Other Causes | Living | Living |
| 168.9666667 | 234.3333333 | 118.0333333 | 24.9 | 145.4333333 | 274.4 | 265 | 30.13333333 | 172.9 | 262.6333333 | 112 |
| NO | NO | NO | YES | NO | YES | NO | NO | NO | YES | NO |

| | | | | | | | | | | |
|---|---|---|---|---|---|---|---|---|---|---|
| MB_2951 | MB_2960 | MB_2970 | MB_2971 | MB_2933 | MB_3479 | MB_3490 | MB_3487 | MB_3459 | MB_3476 | MB_3492 |
| WT | WT | MUT | WT | WT | WT | WT | WT | WT | MUT | WT |
| WT | WT | NULL | WT | WT | WT | WT | WT | WT | MISS | WT |
| | | C124Lfs*25 | | | | | | | C238G | |
| ERpHER2n | ERpHER2n | ERpHER2n | ERpHER2n | ERpHER2n | ERpHER2n | ERpHER2n | ERpHER2n | ERpHER2n | ERnHER2n | ERpHER2n |
| Living | Died of Other Causes | Died of Disease | Living | Died of Other Causes | Living | Died of Disease | Living | Living | Died of Other Causes | Died of Disease |
| 263.3666667 | 222.2 | 252.8666667 | 264.2333333 | 161.9333333 | 231.6666667 | 102.4 | 237.1333333 | 241.4 | 159.0666667 | 118.1333333 |
| NO | NO | NO | NO | NO | NO | NO | NO | NO | NO | NO |

| | | | | | | | | | | |
|---|---|---|---|---|---|---|---|---|---|---|
| MB_3452 | MB_3303 | MB_3328 | MB_3300 | MB_3277 | MB_3254 | MB_3275 | MB_3272 | MB_3271 | MB_3292 | MB_3840 |
| MUT | WT | WT | WT | MUT | WT | WT | MUT | MUT | MUT | MUT |
| NULL | WT | WT | WT | MISS | WT | WT | NULL | NULL | NULL | MISS |
| C229Yfs*10 | | | | R273C | | | I255Sfs*90 | D49Mfs*4 | R306* | R110H |
| ERpHER2n | ERpHER2n | ERpHER2n | ERpHER2n | ERnHER2n | ERpHER2n | ERpHER2n | HER2p | ERnHER2n | ERnHER2n | ERpHER2n |
| Died of Disease | Living | Died of Other Causes | Living | Died of Disease | Living | Died of Disease | Died of Disease | Died of Other Causes | Living | Living |
| 119.8666667 | 243.9 | 191.1333333 | 247 | 32.93333333 | 123.9 | 41.53333333 | 22.66666667 | 65.46666667 | 217.7666667 | 287.2333333 |
| NO | NO | NO | NO | YES | NO | NO | NO | NO | NO | NO |

| | | | | | | | | | | |
|---|---|---|---|---|---|---|---|---|---|---|
| MB_2536 | MB_2564 | MB_2513 | MB_2617 | MB_2614 | MB_2632 | MB_2556 | MB_2624 | MB_2867 | MB_2922 | MB_2917 |
| WT | WT | MUT | MUT | WT | MUT | WT | WT | WT | MUT | MUT |
| WT | WT | MISS | MISS | WT | MISS | WT | WT | WT | NULL | NULL |
| | | L344R | H179R | | G199V | | | | P152Rfs*18 | X126_splice |
| ERpHER2n | ERpHER2n | HER2p | ERpHER2n | ERpHER2n | HER2p | ERnHER2n | ERpHER2n | ERpHER2n | ERnHER2n | ERnHER2n |
| Died of Other Causes | Living | Died of Disease | Died of Disease | Died of Disease | Died of Disease | Died of Other Causes | Living | Living | Died of Disease | Died of Disease |
| 47.9 | 285.4333333 | 59.7 | 89.1 | 64.93333333 | 19.1 | 220.9 | 128.5333333 | 269.3333333 | 14.4 | 31.93333333 |
| NO | NO | NO | NO | YES | YES | NO | NO | NO | YES | YES |

| | | | | | | | | | | |
|---|---|---|---|---|---|---|---|---|---|---|
| MB_2858 | MB_2854 | MB_2853 | MB_3235 | MB_3165 | MB_3171 | MB_3167 | MB_3222 | MB_3706 | MB_3497 | MB_3488 |
| MUT | MUT | WT | MUT | MUT | WT | WT | WT | MUT | WT | MUT |
| MISS | MISS | WT | MISS | NULL | WT | WT | WT | NULL | WT | NULL |
| R267P | F134L | | R248W | R213* | | | | R213* | | Q52Lfs*68 |
| ERpHER2n | ERpHER2n | ERpHER2n | HER2p | ERpHER2n | ERpHER2n | ERpHER2n | ERpHER2n | ERnHER2n | HER2p | HER2p |
| Died of Other Causes | Living | Living | Living | Living | Died of Disease | Living | Died of Other Causes | Died of Disease | Died of Disease | Died of Disease |
| 141.5666667 | 270.4333333 | 269.6333333 | 236.1333333 | 221.2333333 | 219.6666667 | 136.9333333 | 23.93333333 | 38.8 | 15.5 | 27.46666667 |
| NO | NO | NO | NO | YES | NO | NO | NO | NO | YES | YES |

| | | | | | | | | | | | |
|---|---|---|---|---|---|---|---|---|---|---|---|
| MB_3711 | MB_3548 | MB_3506 | MB_3525 | MB_3606 | MB_3453 | MB_3545 | MB_3470 | MB_2765 | MB_2763 | MB_2767 | MB_2790 |
| WT | WT | WT | WT | MUT | MUT | MUT | MUT | WT | WT | WT | WT |
| WT | WT | WT | WT | NULL | MISS | NULL | MISS | WT | WT | WT | WT |
| | | | | Q331Rfs*14 | G262V | D393Tfs*29 | R248G | | | | |
| ERpHER2n | ERpHER2n | ERpHER2n | ERpHER2n | HER2p | ERnHER2n | ERpHER2n | HER2p | ERpHER2n | ERpHER2n | ERpHER2n | ERpHER2n |
| Living | Died of Other Causes | Living | Living | Living | Died of Disease | Died of Disease | Died of Disease | Died of Other Causes | Living | Living | Living |
| 222.1 | 234.5333333 | 240.8333333 | 236.7 | 110.6333333 | 32.83333333 | 232.7333333 | 62.53333333 | 227.8666667 | 275.6 | 271.3333333 | 116.9333333 |
| NO | NO | NO | NO | NO | YES | NO | YES | NO | NO | NO | NO |

| | | | | | | | | | | |
|---|---|---|---|---|---|---|---|---|---|---|
| MB_2769 | MB_2842 | MB_2772 | MB_2779 | MB_2774 | MB_2983 | MB_3005 | MB_2977 | MB_2996 | MB_2999 | MB_2994 |
| WT | MUT | WT | MUT | WT | MUT | WT | WT | WT | WT | WT |
| WT | MISS | WT | MISS | WT | NULL | WT | WT | WT | WT | WT |
| | L194R | | G245C | | Y107* | | | | | |
| ERpHER2n | ERnHER2n | ERpHER2n | ERpHER2n | ERpHER2n | HER2p | ERpHER2n | ERpHER2n | ERpHER2n | ERpHER2n | HER2p |
| Living | Died of Disease | Died of Other Causes | Living | Died of Other Causes | Died of Other Causes | Living | Died of Other Causes | Living | Living | Living |
| 160 | 19.73333333 | 150.7333333 | 275.2333333 | 131.6666667 | 180.8333333 | 253.0666667 | 110.6 | 191.2333333 | 262 | 254.5 |
| NO | YES | NO | NO | NO | NO | NO | NO | NO | NO | NO |

| | | | | | | | | | | | |
|---|---|---|---|---|---|---|---|---|---|---|---|
| MB_3025 | MB_2984 | MB_3002 | MB_2845 | MB_2912 | MB_2844 | MB_2847 | MB_2849 | MB_2851 | MB_2848 | MB_2863 | MB_2840 |
| MUT | MUT | WT | WT | MUT | MUT | MUT | MUT | MUT | WT | WT | WT |
| NULL | MISS | WT | WT | NULL | NULL | NULL | NULL | NULL | WT | WT | WT |
| N239* | N239S | | | V157Tfs*26 | I162delinsRL | X225_splice | T155Pfs*23 | E294* | | | |
| HER2p | HER2p | ERpHER2n | ERpHER2n | ERnHER2n | ERpHER2n | HER2p | ERnHER2n | ERpHER2n | ERpHER2n | ERpHER2n | ERpHER2n |
| Living | Living | Died of Other Causes | Living | Living | Living | Died of Disease | Died of Disease | Died of Disease | Died of Other Causes | Living | Living |
| 256 | 261.2 | 140.5666667 | 264.7666667 | 267.4 | 146.7333333 | 39.53333333 | 43.2 | 159 | 227.7333333 | 258.3333333 | 269 |
| YES | NO | NO | NO | YES | NO | NO | NO | NO | NO | NO | NO |

| | | | | | | | | | | |
|---|---|---|---|---|---|---|---|---|---|---|
| MB_2843 | MB_3013 | MB_3058 | MB_3016 | MB_3057 | MB_3008 | MB_3021 | MB_3032 | MB_3007 | MB_3033 | MB_3026 |
| MUT | WT | WT | MUT | MUT | WT | WT | WT | WT | WT | WT |
| MISS | WT | WT | NULL | NULL | WT | WT | WT | WT | WT | WT |
| S241C | | | R342Efs*3 | A138Pfs*32 | | | | | | |
| ERpHER2n | ERpHER2n | ERnHER2n | ERpHER2n | ERnHER2n | ERpHER2n | ERpHER2n | ERpHER2n | ERpHER2n | ERpHER2n | ERpHER2n |
| Died of Other Causes | Living | Living | Living | Died of Disease | Living | Died of Other Causes | Living | Died of Disease | Died of Disease | Living |
| 231.5333333 | 238.3666667 | 254.2666667 | 258.8666667 | 32.03333333 | 149.6 | 217.5666667 | 250.1333333 | 114.9 | 70.4 | 261.2 |
| NO | NO | YES | NO | YES | NO | NO | NO | NO | NO | NO |

| | | | | | | | | | | |
|---|---|---|---|---|---|---|---|---|---|---|
| MB_3006 | MB_3439 | MB_3437 | MB_3395 | MB_3386 | MB_3382 | MB_3379 | MB_3430 | MB_3389 | MB_3378 | MB_3110 |
| WT | WT | WT | MUT | MUT | WT | WT | WT | WT | WT | WT |
| WT | WT | WT | MISS | MISS | WT | WT | WT | WT | WT | WT |
| | | | Y220C | R248Q | | | | | | |
| ERpHER2n | ERpHER2n | ERpHER2n | ERnHER2n | HER2p | HER2p | HER2p | ERpHER2n | ERpHER2n | ERpHER2n | ERpHER2n |
| Died of Disease | Living | Living | Living | Died of Disease | Living | Died of Other Causes | Died of Other Causes | Living | Died of Other Causes | Living |
| 15.7 | 188.7333333 | 108.7666667 | 243.7666667 | 55.63333333 | 136.2333333 | 113.8333333 | 219.7666667 | 240.7 | 87.53333333 | 100.7333333 |
| NO | NO | NO | NO | YES | NO | NO | NO | NO | NO | NO |

| | | | | | | | | | | |
|---|---|---|---|---|---|---|---|---|---|---|
| MB_3079 | MB_3103 | MB_3102 | MB_3085 | MB_3122 | MB_3104 | MB_3123 | MB_3121 | MB_2835 | MB_2838 | MB_2801 |
| WT | WT | WT | WT | MUT | WT | MUT | WT | WT | WT | MUT |
| WT | WT | WT | WT | MISS | WT | MISS | WT | WT | WT | NULL |
| | | | | P278T | | S241Y | | | | R213* |
| ERpHER2n | HER2p | ERpHER2n | ERpHER2n | ERpHER2n | HER2p | ERnHER2n | ERpHER2n | ERpHER2n | ERpHER2n | ERpHER2n |
| Living | Died of Disease | Living | Died of Disease | Living | Died of Disease | Living | Died of Other Causes | Died of Other Causes | Living | Living |
| 149.6 | 28.73333333 | 258.3333333 | 210.9666667 | 257.7666667 | 75.36666667 | 167.4333333 | 131.3333333 | 55.03333333 | 270.3 | 271.2666667 |
| NO | NO | NO | NO | NO | NO | YES | NO | NO | NO | NO |

| | | | | | | | | | | |
|---|---|---|---|---|---|---|---|---|---|---|
| MB_2834 | MB_2796 | MB_2857 | MB_2819 | MB_2781 | MB_2793 | MB_2827 | MB_2814 | MB_2791 | MB_2803 | MB_2797 |
| MUT | MUT | MUT | WT | WT | WT | MUT | WT | WT | WT | WT |
| MISS | MISS | NULL | WT | WT | WT | NULL | WT | WT | WT | WT |
| I232T | Y220S | R306* | | | | X224_splice | | | | |
| ERnHER2n | ERpHER2n | ERnHER2n | ERpHER2n | ERpHER2n | ERpHER2n | ERnHER2n | ERpHER2n | ERpHER2n | ERpHER2n | ERpHER2n |
| Died of Other Causes | Died of Disease | Living | Living | Died of Other Causes | Living | Living | Died of Other Causes | Living | Died of Disease | Living |
| 153.8333333 | 87.7 | 250.6666667 | 270.5666667 | 40.43333333 | 267.2333333 | 235.6666667 | 231.0333333 | 271.9333333 | 85.86666667 | 252.9666667 |
| NO | NO | YES | NO | NO | NO | YES | NO | NO | NO | NO |

| | | | | | | | | | | | | |
|---|---|---|---|---|---|---|---|---|---|---|---|---|
| MB_2795 | MB_2786 | MB_3850 | MB_3838 | MB_3707 | MB_3600 | MB_3824 | MB_3823 | MB_3781 | MB_2642 | MB_2686 | MB_2616 | MB_2634 |
| WT | WT | WT | WT | MUT | WT | MUT | WT | WT | WT | WT | MUT | WT |
| WT | WT | WT | WT | MISS | WT | NULL | WT | WT | WT | WT | MISS | WT |
| | | | | E285K | | I162Tfs*14 | | | | | R248W | |
| ERpHER2n | HER2p | HER2p | ERpHER2n | ERpHER2n | ERpHER2n | ERpHER2n | ERpHER2n | ERpHER2n | ERpHER2n | ERpHER2n | ERpHER2n | ERpHER2n |
| Living | Died of Disease | Living | Died of Disease | Died of Disease | Living | Died of Disease | Living | Living | Died of Disease | Living | Died of Disease | Living |
| 272.1 | 68.26666667 | 72.3 | 134.4666667 | 228.7666667 | 214.7 | 126.4666667 | 222.5 | 224.5666667 | 45.16666667 | 177.2666667 | 219.1666667 | 274.0333333 |
| NO | NO | NO | NO | NO | NO | NO | NO | NO | YES | NO | NO | NO |

| | | | | | | | | | | | | |
|---|---|---|---|---|---|---|---|---|---|---|---|---|
| MB_2613 | MB_3842 | MB_3866 | MB_3874 | MB_3871 | MB_3702 | MB_3105 | MB_3852 | MB_3854 | MB_3865 | MB_3228 | MB_3028 | MB_3502 |
| WT | WT | MUT | MUT | WT | MUT | WT | MUT | WT | WT | WT | MUT | MUT |
| WT | WT | NULL | MISS | WT | MISS | WT | MISS | WT | WT | WT | MISS | MISS |
| | | R196* | R110H | | L257Q | | L194R | | | | R248Q | R273H |
| ERpHER2n | ERpHER2n | HER2p | ERpHER2n | ERpHER2n | ERnHER2n | ERpHER2n | HER2p | ERpHER2n | ERpHER2n | ERpHER2n | HER2p | ERnHER2n |
| Living | Living | Living | Living | Died of Other Causes | Living | Living | Living | Living | Living | Living | Died of Disease | Living |
| 163.4 | 226.0666667 | 225.5 | 186.4333333 | 172.8 | 230.4666667 | 252.3333333 | 225.5 | 99.23333333 | 199.9666667 | 252 | 42.56666667 | 229.3333333 |
| NO | NO | NO | NO | NO | YES | NO | NO | NO | NO | NO | NO | YES |

| | | | | | | | | | | | | |
|---|---|---|---|---|---|---|---|---|---|---|---|---|
| MB_3567 | MB_3752 | MB_3450 | MB_0476 | MB_0610 | MB_0451 | MB_0133 | MB_0048 | MB_0083 | MB_0053 | MB_0056 | MB_0068 | MB_0079 |
| MUT | MUT | WT | MUT | WT | WT | WT | WT | WT | WT | WT | WT | MUT |
| MISS | NULL | WT | MISS | WT | WT | WT | WT | WT | WT | WT | WT | MISS |
| Y126N | R196* | | R175H | | | | | | | | | R273C |
| ERnHER2n | ERnHER2n | ERpHER2n | ERpHER2n | ERpHER2n | ERpHER2n | ERpHER2n | HER2p | ERpHER2n | ERpHER2n | ERpHER2n | ERpHER2n | ERnHER2n |
| Living | Died of Disease | Living | Living | Living | Living | Living | Living | Died of Disease | Living | Living | Living | Died of Disease |
| 236.0333333 | 75.33333333 | 240.4666667 | 130.4333333 | 76.7 | 50.06666667 | 151 | 103.8333333 | 86.06666667 | 161.0666667 | 62.86666667 | 103.1333333 | 28.5 |
| YES | NO | NO | YES | NO | NO | YES | YES | NO | NO | NO | NO | YES |

| | | | | | | | | | | | |
|---|---|---|---|---|---|---|---|---|---|---|---|
| MB_0108 | MB_0006 | MB_0014 | MB_0022 | MB_0039 | MB_0062 | MB_0054 | MB_0081 | MB_0071 | MB_0099 | MB_0064 | MB_0107 |
| MUT | WT | WT | WT | WT | MUT | WT | WT | WT | WT | WT | MUT |
| MISS | WT | WT | WT | WT | NULL | WT | WT | WT | WT | WT | MISS |
| G245S | | | | | S303Efs*3 | | | | | | P177R |
| ERpHER2n | ERpHER2n | ERpHER2n | ERpHER2n | ERpHER2n | ERnHER2n | ERpHER2n | ERpHER2n | ERpHER2n | ERpHER2n | ERpHER2n | ERpHER2n |
| Died of Disease | Living | Living | Died of Other Causes | Living | Living | Living | Living | Died of Other Causes | Died of Disease | Living | Living |
| 42.7 | 164.9333333 | 164.3333333 | 99.53333333 | 163.5333333 | 153.9666667 | 160.3 | 69.5 | 131 | 132.1 | 108.9333333 | 158.0333333 |
| YES | YES | YES | NO | NO | YES | NO | NO | NO | YES | NO | NO |

| | | | | | | | | | |
|---|---|---|---|---|---|---|---|---|---|
| MB_4681 | MB_4626 | MB_4639 | MB_4673 | MB_4711 | MB_4607 | MB_4669 | MB_4015 | MB_2931 | MB_2927 |
| WT | WT | WT | WT | MUT | WT | WT | MUT | MUT | WT |
| WT | WT | WT | WT | NULL | WT | WT | NULL | NULL | WT |
| | | | | W91* | | | E180* | H178Pfs*3 | |
| ERpHER2n | HER2p | ERpHER2n | ERpHER2n | HER2p | ERpHER2n | ERpHER2n | ERnHER2n | ERpHER2n | ERpHER2n |
| Died of Disease | Died of Disease | Died of Disease | Died of Disease | Died of Disease | Living | Died of Other Causes | Died of Disease | Died of Other Causes | Died of Disease |
| 251.8 | 143 | 119 | 87 | 31.46666667 | 121.7333333 | 119.4666667 | 11.3 | 228.1 | 263.7 |
| NO | NO | NO | NO | YES | NO | NO | NO | NO | NO |

| | | | | | | | | | |
|---|---|---|---|---|---|---|---|---|---|
| MB_2919 | MB_2944 | MB_2929 | MB_2932 | MB_2724 | MB_2712 | MB_3062 | MB_3050 | MB_4722 | MB_4731 |
| WT | MUT | MUT | WT | WT | WT | MUT | MUT | WT | MUT |
| WT | MISS | MISS | WT | WT | WT | NULL | NULL | WT | MISS |
| | S127F | R181C | | | | N239_S240insT | Q192* | | R282W |
| ERpHER2n | ERpHER2n | ERnHER2n | ERpHER2n | ERnHER2n | ERpHER2n | ERnHER2n | ERpHER2n | ERpHER2n | HER2p |
| Died of Other Causes | Died of Disease | Died of Other Causes | Living | Died of Disease | Died of Other Causes | Living | Died of Other Causes | Living | Died of Disease |
| 240.0333333 | 27.2 | 262.8666667 | 108.4333333 | 108.0666667 | 234.4333333 | 146.3666667 | 255.1 | 86.1 | 44.86666667 |
| NO | NO | YES | NO | NO | NO | NO | NO | NO | YES |

| | | | | | | | | | | | |
|---|---|---|---|---|---|---|---|---|---|---|---|
| MB_4725 | MB_4707 | MB_4643 | MB_3363 | MB_3383 | MB_3344 | MB_3381 | MB_3297 | MB_3301 | MB_3088 | MB_2764 | MB_2744 |
| MUT | MUT | MUT | WT | MUT | WT | WT | MUT | WT | WT | WT | WT |
| MISS | MISS | MISS | WT | MISS | WT | WT | MISS | WT | WT | WT | WT |
| R273G | T125K | R342P | | R175H | | | R342P | | | | |
| HER2p | ERnHER2n | HER2p | ERpHER2n | ERnHER2n | ERpHER2n | ERpHER2n | ERnHER2n | ERpHER2n | ERpHER2n | ERpHER2n | ERpHER2n |
| Died of Disease | Living | Died of Disease | Died of Disease | Died of Disease | Living | Living | Living | Living | Living | Died of Disease | Living |
| 98.76666667 | 221.2 | 41.46666667 | 10.86666667 | 23.2 | 241.2666667 | 236.6333333 | 236.0666667 | 248.7666667 | 256.5 | 270.1333333 | 275.6333333 |
| NO | NO | NO | NO | YES | NO | NO | YES | NO | NO | NO | NO |

| | | | | | | | | | | |
|---|---|---|---|---|---|---|---|---|---|---|
| MB_7114 | MB_7118 | MB_7113 | MB_7130 | MB_7149 | MB_7140 | MB_7148 | MB_7208 | MB_7170 | MB_7174 | MB_7252 |
| MUT | WT | WT | WT | WT | WT | WT | WT | WT | MUT | MUT |
| NULL | WT | WT | WT | WT | WT | WT | WT | WT | MISS | NULL |
| C242Afs*5 | | | | | | | | | Y220C | F113del |
| ERnHER2n | ERpHER2n | ERpHER2n | ERpHER2n | ERpHER2n | ERpHER2n | ERpHER2n | ERnHER2n | ERpHER2n | ERpHER2n | ERnHER2n |
| Living | Living | Died of Other Causes | Died of Disease | Died of Other Causes | Died of Disease | Died of Disease | Died of Disease | Living | Living | Died of Other Causes |
| 123.2666667 | 140.5666667 | 136.7 | 102.9666667 | 44.63333333 | 90.8 | 19 | 45.93333333 | 155.4 | 140.7666667 | 35.7 |
| NO | NO | NO | NO | NO | NO | YES | YES | NO | NO | NO |

| | | | | | | | | | |
|---|---|---|---|---|---|---|---|---|---|
| MB_7244 | MB_7187 | MB_7251 | MB_7173 | MB_3001 | MB_2952 | MB_7097 | MB_7104 | MB_7099 | MB_7069 |
| WT | MUT | WT | WT | MUT | WT | WT | MUT | WT | MUT |
| WT | MISS | WT | WT | MISS | WT | WT | MISS | WT | NULL |
| | C242S | | | R273H | | | R175H | | P153Afs*28 |
| ERpHER2n | HER2p | HER2p | ERpHER2n | ERnHER2n | ERpHER2n | ERpHER2n | ERpHER2n | ERpHER2n | HER2p |
| Living | Died of Disease | Died of Disease | Died of Other Causes | Living | Died of Other Causes | Died of Disease | Died of Other Causes | Died of Other Causes | Living |
| 184.1666667 | 146.8333333 | 21.56666667 | 63.53333333 | 263.2333333 | 189.1 | 65.16666667 | 122.8333333 | 163.1666667 | 114.6 |
| NO | YES | NO | NO | YES | NO | YES | NO | NO | NO |

| | | | | | | | | | | |
|---|---|---|---|---|---|---|---|---|---|---|
| MB_7066 | MB_7100 | MB_7067 | MB_7073 | MB_7063 | MB_7102 | MB_7068 | MB_7075 | MB_7053 | MB_7074 | MB_7057 |
| WT | WT | MUT | MUT | WT | WT | MUT | WT | MUT | WT | MUT |
| WT | WT | MISS | NULL | WT | WT | MISS | WT | NULL | WT | MISS |
| | | L194R | G302Rfs*4 | | | R273C | | R306* | | S241F |
| ERnHER2n | ERpHER2n | HER2p | HER2p | ERpHER2n | ERpHER2n | HER2p | ERpHER2n | ERpHER2n | ERpHER2n | ERnHER2n |
| Living | Died of Other Causes | Living | Died of Other Causes | Living | Died of Other Causes | Living | Living | Living | Living | Died of Disease |
| 89.76666667 | 156.3333333 | 105 | 137.9333333 | 86.6 | 104.1 | 98.46666667 | 111.6333333 | 122.1333333 | 108.1666667 | 43.83333333 |
| YES | NO | NO | YES | NO | NO | NO | NO | NO | NO | YES |

| | | | | | | | | | | |
|---|---|---|---|---|---|---|---|---|---|---|
| MB_7109 | MB_7051 | MB_7071 | MB_7106 | MB_7059 | MB_7107 | MB_7076 | MB_7111 | MB_7015 | MB_7026 | MB_7013 |
| WT | MUT | WT | WT | MUT | WT | WT | MUT | WT | WT | WT |
| WT | MISS | WT | WT | NULL | WT | WT | MISS | WT | WT | WT |
| | P278S | | | E298* | | | R273H | | | |
| ERpHER2n | ERpHER2n | ERpHER2n | ERpHER2n | HER2p | ERpHER2n | ERpHER2n | ERpHER2n | ERpHER2n | ERpHER2n | ERpHER2n |
| Living | Died of Disease | Living | Died of Disease | Living | Died of Disease | Died of Other Causes | Died of Disease | Living | Died of Other Causes | Living |
| 117.5333333 | 85.93333333 | 102.0666667 | 37.5 | 119.3333333 | 56.93333333 | 142.6666667 | 30.16666667 | 78.16666667 | 111.7 | 46.43333333 |
| NO | NO | NO | NO | YES | NO | NO | NO | NO | NO | NO |

| | | | | | | | | | | |
|---|---|---|---|---|---|---|---|---|---|---|
| MB_7023 | MB_7024 | MB_7014 | MB_7017 | MB_7019 | MB_3510 | MB_3500 | MB_3396 | MB_3435 | MB_7186 | MB_7142 |
| MUT | WT | MUT | MUT | WT | WT | MUT | MUT | WT | WT | WT |
| MISS | WT | NULL | MISS | WT | WT | NULL | NULL | WT | WT | WT |
| H193R | | M44Cfs*79 | A159V | | | W53* | A69Vfs*54 | | | |
| ERnHER2n | ERpHER2n | ERpHER2n | ERnHER2n | ERpHER2n | ERpHER2n | ERnHER2n | ERnHER2n | HER2p | ERpHER2n | ERpHER2n |
| Died of Disease | Living | Living | Living | Died of Other Causes | Died of Other Causes | Living | Living | Died of Disease | Living | Died of Other Causes |
| 27.96666667 | 70.26666667 | 68.7 | 135.3333333 | 80.73333333 | 59.6 | 239.3 | 226.7333333 | 25.43333333 | 4.866666667 | 175.9 |
| YES | NO | NO | YES | NO | NO | NO | YES | YES | NO | NO |

| | | | | | | | | | | |
|---|---|---|---|---|---|---|---|---|---|---|
| MB_7196 | MB_7171 | MB_7145 | MB_7132 | MB_7189 | MB_7137 | MB_7181 | MB_7155 | MB_7138 | MB_3295 | MB_3218 |
| MUT | WT | MUT | WT | MUT | WT | WT | MUT | WT | MUT | MUT |
| NULL | WT | NULL | WT | NULL | WT | WT | NULL | WT | MISS | MISS |
| X224_splice | | E51* | | P177_C182del | | | C242Afs*5 | | P278R | L137Q |
| ERpHER2n | ERpHER2n | ERnHER2n | ERpHER2n | ERpHER2n | ERpHER2n | ERpHER2n | ERnHER2n | ERpHER2n | ERpHER2n | ERnHER2n |
| Died of Disease | Died of Other Causes | Living | Living | Died of Disease | Living | Living | Living | Died of Other Causes | Died of Other Causes | Living |
| 107.1 | 160.4 | 138.3333333 | 131.6 | 143.6 | 179.8 | 157.5333333 | 149.7666667 | 123 | 158.5333333 | 248.7666667 |
| NO | NO | YES | NO | YES | NO | NO | YES | NO | NO | YES |

| | | | | | | | | | |
|---|---|---|---|---|---|---|---|---|---|
| MB_7232 | MB_7281 | MB_7270 | MB_7231 | MB_7220 | MB_7277 | MB_7280 | MB_7227 | MB_7278 | MB_2610 |
| WT | MUT | MUT | WT | WT | WT | WT | WT | WT | WT |
| WT | MISS | NULL | WT | WT | WT | WT | WT | WT | WT |
| | R175G | N131del | | | | | | | |
| ERpHER2n | HER2p | ERnHER2n | ERpHER2n | ERpHER2n | ERpHER2n | ERpHER2n | ERpHER2n | ERpHER2n | ERpHER2n |
| Living | Died of Disease | Living | Died of Other Causes | Died of Other Causes | Living | Living | Died of Other Causes | Died of Other Causes | Living |
| 177.6 | 49.53333333 | 175.1666667 | 199.0333333 | 183.2 | 187.0333333 | 195.5333333 | 63.2 | 241.3 | 280.7 |
| NO | YES | YES | NO | NO | NO | NO | NO | NO | NO |

| | | | | | | | | | |
|---|---|---|---|---|---|---|---|---|---|
| MB_2517 | MB_2896 | MB_2901 | MB_2895 | MB_2900 | MB_2939 | MB_7008 | MB_7002 | MB_7000 | MB_7006 |
| MUT | WT | WT | MUT | MUT | WT | WT | WT | WT | WT |
| MISS | WT | WT | NULL | MISS | WT | WT | WT | WT | WT |
| L111P | | | R306* | K132N | | | | | |
| HER2p | ERpHER2n | ERpHER2n | HER2p | ERpHER2n | HER2p | ERnHER2n | ERpHER2n | ERpHER2n | ERpHER2n |
| Died of Other Causes | Died of Other Causes | Died of Disease | Died of Disease | Died of Other Causes | Died of Other Causes | Died of Other Causes | Living | Living | Living |
| 268.4333333 | 237.3666667 | 218.3 | 196.5666667 | 20.66666667 | 101.2666667 | 50.9 | 57.93333333 | 97.4 | 109.9333333 |
| NO | NO | NO | NO | NO | NO | NO | NO | NO | NO |

| | | | | | | | | | | | |
|---|---|---|---|---|---|---|---|---|---|---|---|
| MB_7007 | MB_7011 | MB_7001 | MB_7009 | MB_7003 | MB_7005 | MB_7010 | MB_3153 | MB_3211 | MB_3181 | MB_3252 | MB_3266 |
| MUT | WT | WT | WT | WT | WT | WT | MUT | WT | MUT | WT | MUT |
| MISS | WT | WT | WT | WT | WT | WT | MISS | WT | NULL | WT | MISS |
| E224D | | | | | | | S241F | | R342* | | Y220C |
| ERnHER2n | ERpHER2n | HER2p | ERnHER2n | ERpHER2n | ERpHER2n | ERpHER2n | ERnHER2n | ERnHER2n | ERpHER2n | ERpHER2n | ERpHER2n |
| Living | Died of Other Causes | Living | Died of Disease | Died of Other Causes | Living | Living | Living | Living | Died of Disease | Living | Died of Disease |
| 58.6 | 70.06666667 | 102.7 | 54.76666667 | 91.23333333 | 45.33333333 | 74.03333333 | 227.9333333 | 145.5 | 178.1666667 | 99.7 | 51.2 |
| YES | NO | NO | YES | NO | NO | NO | YES | NO | NO | NO | NO |

| | | | | | | | | | | | |
|---|---|---|---|---|---|---|---|---|---|---|---|
| MB_7082 | MB_7093 | MB_7083 | MB_7091 | MB_7065 | MB_7085 | MB_7096 | MB_7018 | MB_7088 | MB_7055 | MB_7080 | MB_7161 |
| MUT | WT | MUT | MUT | WT | WT | WT | WT | MUT | MUT | WT | WT |
| MISS | WT | MISS | MISS | WT | WT | WT | WT | MISS | NULL | WT | WT |
| C135W | | G154V | Y236C | | | | | M246V | R306* | | |
| HER2p | ERpHER2n | ERpHER2n | ERpHER2n | ERpHER2n | ERpHER2n | ERpHER2n | ERpHER2n | HER2p | ERnHER2n | ERpHER2n | ERpHER2n |
| Living | Died of Other Causes | Living | Living | Living | Living | Living | Died of Disease | Living | Died of Disease | Living | Living |
| 98.26666667 | 125.8333333 | 105.4666667 | 110.2666667 | 107.8666667 | 110.1 | 124 | 86.53333333 | 123.5333333 | 9.133333333 | 91.23333333 | 121.5333333 |
| YES | YES | YES | NO | NO | NO | NO | YES | NO | NO | NO | NO |

| | | | | | | | | | | |
|---|---|---|---|---|---|---|---|---|---|---|
| MB_7115 | MB_7123 | MB_7176 | MB_7121 | MB_7172 | MB_7164 | MB_7127 | MB_7128 | MB_7157 | MB_7092 | MB_7135 |
| WT | WT | WT | WT | WT | WT | WT | WT | WT | WT | WT |
| WT | WT | WT | WT | WT | WT | WT | WT | WT | WT | WT |
| HER2p | ERpHER2n | ERpHER2n | ERnHER2n | ERpHER2n | ERpHER2n | ERpHER2n | HER2p | ERpHER2n | ERpHER2n | HER2p |
| Died of Other Causes | Living | Died of Other Causes | Died of Other Causes | Living | Living | Living | Died of Other Causes | Living | Living | Died of Disease |
| 141.7333333 | 137.6666667 | 194.4 | 88.83333333 | 149.3 | 211.4 | 135.3333333 | 60.26666667 | 219.6333333 | 122.4 | 26.73333333 |
| NO | NO | NO | YES | NO | NO | NO | NO | NO | NO | YES |

| | | | | | | | | | |
|---|---|---|---|---|---|---|---|---|---|
| MB_3754 | MB_3536 | MB_3748 | MB_3530 | MB_3582 | MB_3528 | MB_3576 | MB_2770 | MB_2758 | MB_2771 |
| WT | MUT | MUT | WT | WT | WT | MUT | WT | MUT | MUT |
| WT | NULL | NULL | WT | WT | WT | NULL | WT | MISS | NULL |
| | Q192* | R306* | | | | M340Ifs*7 | | V197E | L111Ffs*40 |
| ERpHER2n | ERpHER2n | ERpHER2n | ERpHER2n | ERpHER2n | HER2p | ERpHER2n | ERpHER2n | HER2p | ERnHER2n |
| Died of Disease | Died of Disease | Living | Died of Other Causes | Died of Other Causes | Living | Died of Disease | Living | Died of Disease | Died of Disease |
| 92.73333333 | 86.23333333 | 228 | 112.9666667 | 229.3666667 | 113.6666667 | 24.86666667 | 274.3666667 | 35 | 44.83333333 |
| NO | NO | NO | NO | NO | NO | NO | NO | YES | NO |

| | | | | | | | | |
|---|---|---|---|---|---|---|---|---|
| MB_7152 | MB_7143 | MB_7197 | MB_7153 | MB_7144 | MB_7185 | MB_7199 | MB_7158 | MB_7163 |
| WT | WT | WT | WT | MUT | WT | WT | MUT | WT |
| WT | WT | WT | WT | MISS | WT | WT | MISS | WT |
| | | | | R282W | | | R175H | |
| ERpHER2n | HER2p | ERpHER2n | ERpHER2n | ERpHER2n | ERpHER2n | ERpHER2n | ERnHER2n | ERpHER2n |
| Died of Other Causes | Living | Died of Other Causes | Living | Died of Other Causes | Died of Disease | Died of Other Causes | Died of Disease | Died of Disease |
| 195.8 | 186.3666667 | 187.8666667 | 142.8 | 102.6333333 | 125.0333333 | 76 | 46.43333333 | 52.63333333 |
| NO | NO | NO | NO | NO | NO | NO | YES | YES |

| | | | | | | | | |
|---|---|---|---|---|---|---|---|---|
| MB_7215 | MB_7200 | MB_7193 | MB_7188 | MB_7198 | MB_7256 | MB_7258 | MB_7194 | MB_7254 |
| WT | WT | WT | WT | MUT | MUT | MUT | WT | WT |
| WT | WT | WT | WT | NULL | MISS | NULL | WT | WT |
| | | | | K291* | F134C | L194Pfs*13 | | |
| ERpHER2n | ERpHER2n | ERpHER2n | ERpHER2n | ERpHER2n | HER2p | ERnHER2n | ERpHER2n | ERpHER2n |
| Died of Other Causes | Died of Other Causes | Died of Other Causes | Died of Other Causes | Died of Disease | Died of Disease | Died of Disease | Died of Disease | Living |
| 208.9666667 | 97.6 | 87.46666667 | 188.1666667 | 16.73333333 | 34.7 | 15.06666667 | 46.66666667 | 192.2 |
| NO | NO | NO | NO | NO | YES | YES | YES | NO |

| | | | | | | | | | | |
|---|---|---|---|---|---|---|---|---|---|---|
| MB_7262 | MB_2990 | MB_2993 | MB_3031 | MB_2904 | MB_2846 | MB_7154 | MB_7229 | MB_7151 | MB_7228 | MB_7218 |
| MUT | WT | WT | WT | MUT | MUT | MUT | WT | MUT | WT | WT |
| MISS | WT | WT | WT | NULL | MISS | MISS | WT | NULL | WT | WT |
| G262V | | | | R110Lfs*13 | Y234N | R273H | | R196* | | |
| ERpHER2n | ERpHER2n | ERnHER2n | HER2p | ERnHER2n | ERpHER2n | ERnHER2n | ERpHER2n | ERnHER2n | ERpHER2n | ERpHER2n |
| Died of Other Causes | Living | Living | Died of Disease | Died of Disease | Died of Disease | Died of Other Causes | Died of Other Causes | Living | Living | Living |
| 104.0333333 | 260 | 187.0333333 | 35.63333333 | 125.6 | 149.7 | 108.4666667 | 111.6 | 144.4333333 | 239.1666667 | 165.4333333 |
| NO | NO | NO | YES | YES | NO | NO | NO | NO | NO | NO |

| | | | | | | | | | | |
|---|---|---|---|---|---|---|---|---|---|---|
| MB_7147 | MB_7219 | MB_7165 | MB_7150 | MB_3436 | MB_3462 | MB_3417 | MB_2721 | MB_3092 | MB_2821 | MB_2850 |
| WT | WT | MUT | MUT | WT | WT | WT | WT | WT | WT | WT |
| WT | WT | MISS | MISS | WT | WT | WT | WT | WT | WT | WT |
| | | R175H | R175H | | | | | | | |
| ERpHER2n | ERpHER2n | ERnHER2n | ERpHER2n | ERpHER2n | ERpHER2n | ERpHER2n | ERpHER2n | ERpHER2n | ERnHER2n | ERnHER2n |
| Died of Other Causes | Living | Living | Living | Died of Disease | Living | Living | Living | Died of Other Causes | Died of Other Causes | Died of Disease |
| 168.6 | 165.2 | 147.1666667 | 128.9666667 | 50.76666667 | 234.2333333 | 103.4666667 | 251.6333333 | 152.3333333 | 136.0666667 | 9.433333333 |
| NO | NO | YES | NO | YES | NO | NO | NO | NO | NO | YES |

| | | | | | | | | | | |
|---|---|---|---|---|---|---|---|---|---|---|
| MB_2820 | MB_2823 | MB_2792 | MB_7273 | MB_7283 | MB_7284 | MB_7279 | MB_7233 | MB_7287 | MB_7250 | MB_7238 |
| WT | WT | WT | MUT | MUT | WT | MUT | WT | MUT | MUT | MUT |
| WT | WT | WT | MISS | NULL | WT | MISS | WT | MISS | NULL | NULL |
| | | | E258Q | E349* | | L330R | | R273H | A83Gfs*66 | X224_splice |
| ERpHER2n | HER2p | ERpHER2n | HER2p | ERpHER2n | ERpHER2n | ERpHER2n | ERpHER2n | HER2p | HER2p | ERpHER2n |
| Living | Living | Died of Disease | Living | Died of Disease | Died of Other Causes | Died of Disease | Died of Other Causes | Died of Disease | Died of Disease | Died of Disease |
| 254.6333333 | 269.9 | 110.7666667 | 208.2 | 28.86666667 | 203 | 21.3 | 222.0333333 | 96.96666667 | 51 | 23.26666667 |
| NO | NO | NO | YES | YES | NO | NO | NO | NO | NO | NO |

| | | | | | | | | | |
|---|---|---|---|---|---|---|---|---|---|
| MB_7285 | MB_7260 | MB_7264 | MB_7268 | MB_7276 | MB_7212 | MB_7267 | MB_7266 | MB_7216 | MB_7195 |
| MUT | MUT | MUT | WT | WT | WT | MUT | WT | WT | WT |
| MISS | NULL | NULL | WT | WT | WT | NULL | WT | WT | WT |
| T211A | G325* | X125_splice | | | | C229Yfs*10 | | | |
| ERpHER2n | HER2p | ERpHER2n | ERpHER2n | ERpHER2n | ERpHER2n | ERnHER2n | ERpHER2n | ERpHER2n | ERpHER2n |
| Died of Disease | Living | Died of Disease | Living | Living | Died of Other Causes | Died of Other Causes | Died of Disease | Died of Other Causes | Died of Other Causes |
| 109 | 31.16666667 | 42.96666667 | 227.4666667 | 185.3333333 | 52.5 | 195.3666667 | 68.16666667 | 207.4666667 | 80.43333333 |
| NO | YES | NO | NO | NO | NO | YES | NO | NO | NO |

| | | | | | | | | | | | |
|---|---|---|---|---|---|---|---|---|---|---|---|
| MB_7201 | MB_7217 | MB_2778 | MB_2815 | MB_2833 | MB_3614 | MB_7052 | MB_7124 | MB_7122 | MB_7133 | MB_7084 | MB_7101 |
| MUT | WT | WT | WT | MUT | WT | MUT | WT | MUT | WT | MUT | WT |
| MISS | WT | WT | WT | MISS | WT | NULL | WT | MISS | WT | NULL | WT |
| K132N | | | | R175H | | P177_C182del | | R248W | | I50* | |
| ERnHER2n | ERpHER2n | ERpHER2n | ERpHER2n | ERnHER2n | ERpHER2n | ERnHER2n | ERpHER2n | ERpHER2n | ERpHER2n | HER2p | ERpHER2n |
| Living | Died of Other Causes | Died of Disease | Living | Living | Living | Living | Living | Died of Other Causes | Died of Disease | Living | Living |
| 163.4333333 | 90.4 | 181.2333333 | 257.5666667 | 259.7666667 | 235.4 | 106.1333333 | 139.6 | 29.9 | 23.83333333 | 116.5333333 | 124.2666667 |
| NO | NO | NO | NO | YES | NO | NO | NO | NO | NO | NO | NO |

| | | | | | | | | | | | |
|---|---|---|---|---|---|---|---|---|---|---|---|
| MB_7116 | MB_7131 | MB_7090 | MB_7095 | MB_7094 | MB_7087 | MB_7077 | MB_7072 | MB_7041 | MB_7044 | MB_7022 | MB_7070 |
| WT | WT | MUT | WT | WT | MUT | WT | MUT | WT | WT | WT | MUT |
| WT | WT | NULL | WT | WT | NULL | WT | NULL | WT | WT | WT | NULL |
| | | E294* | | | X261_splice | | E349* | | | | N239_S240del |
| ERpHER2n | ERpHER2n | ERnHER2n | ERpHER2n | ERpHER2n | ERnHER2n | ERpHER2n | ERpHER2n | ERpHER2n | ERpHER2n | ERpHER2n | ERpHER2n |
| Died of Disease | Living | Living | Living | Living | Died of Other Causes | Living | Died of Disease | Died of Other Causes | Living | Living | Living |
| 33.36666667 | 134.7333333 | 109.7666667 | 115.6333333 | 113.3666667 | 5.066666667 | 101.2666667 | 52.33333333 | 101.9666667 | 86.83333333 | 85.5 | 108.0666667 |
| NO | NO | YES | NO | NO | YES | NO | NO | NO | NO | NO | NO |

| | | | | | | | | | | |
|---|---|---|---|---|---|---|---|---|---|---|
| MB_7079 | MB_7025 | MB_7062 | MB_7036 | MB_7046 | MB_4839 | MB_5559 | MB_5463 | MB_4744 | MB_4769 | MB_4484 |
| WT | MUT | WT | MUT | WT | WT | MUT | WT | WT | WT | WT |
| WT | NULL | WT | MISS | WT | WT | MISS | WT | WT | WT | WT |
| | Q165* | | Y236S | | | R280G | | | | |
| ERpHER2n | ERnHER2n | ERpHER2n | ERnHER2n | HER2p | ERpHER2n | ERpHER2n | ERpHER2n | ERpHER2n | ERnHER2n | ERpHER2n |
| Living | Living | Living | Living | Died of Other Causes | Living | Died of Disease | Died of Other Causes | Died of Disease | Died of Disease | Died of Disease |
| 91.06666667 | 80.23333333 | 95.86666667 | 120.4333333 | 146.4 | 299.4 | 18.83333333 | 58.13333333 | 153 | 21.7 | 52.3 |
| NO | NO | NO | YES | NO | NO | YES | NO | NO | NO | NO |

| | | | | | | | | | |
|---|---|---|---|---|---|---|---|---|---|
| MB_5558 | MB_4750 | MB_4793 | MB_4843 | MB_5563 | MB_5546 | MB_5513 | MB_5656 | MB_5530 | MB_5553 |
| MUT | WT | MUT | WT | WT | WT | MUT | WT | WT | WT |
| MISS | WT | NULL | WT | WT | WT | NULL | WT | WT | WT |
| G266V | | I195Tfs*52 | | | | R209Kfs*6 | | | |
| HER2p | ERpHER2n | ERnHER2n | ERpHER2n | ERpHER2n | ERpHER2n | ERpHER2n | ERpHER2n | ERpHER2n | ERpHER2n |
| Living | Died of Disease | Died of Other Causes | Living | Died of Other Causes | Living | Died of Disease | Died of Other Causes | Living | Died of Other Causes |
| 25.03333333 | 37.73333333 | 83.36666667 | 265.1333333 | 117.6666667 | 233.2666667 | 36.63333333 | 126.4 | 185.9333333 | 154.5 |
| YES | NO | NO | NO | NO | NO | NO | NO | NO | NO |

| | | | | | | | | | |
|---|---|---|---|---|---|---|---|---|---|
| MB_4836 | MB_4828 | MB_4697 | MB_4679 | MB_4033 | MB_4692 | MB_4675 | MB_4653 | MB_4663 | MB_4660 |
| WT | WT | WT | MUT | MUT | WT | WT | MUT | MUT | MUT |
| WT | WT | WT | NULL | MISS | WT | WT | MISS | MISS | MISS |
| | | | Y107* | E271K | | | L257P | Y220C | I255F |
| ERpHER2n | ERpHER2n | ERpHER2n | ERnHER2n | ERpHER2n | ERpHER2n | ERpHER2n | ERpHER2n | HER2p | ERnHER2n |
| Died of Disease | Died of Disease | Died of Other Causes | Died of Other Causes | Died of Disease | Died of Other Causes | Living | Died of Disease | Living | Died of Disease |
| 23.43333333 | 42.5 | 118 | 292.6666667 | 15.2 | 107.2666667 | 263.0333333 | 48.13333333 | 239.3333333 | 39.86666667 |
| NO | NO | NO | NO | NO | NO | NO | NO | NO | YES |

| | | | | | | | | | | |
|---|---|---|---|---|---|---|---|---|---|---|
| MB_5275 | MB_4654 | MB_2618 | MB_2718 | MB_2643 | MB_2629 | MB_2645 | MB_2626 | MB_6263 | MB_6305 | MB_6185 |
| WT | WT | WT | MUT | WT | WT | MUT | WT | WT | MUT | WT |
| WT | WT | WT | NULL | WT | WT | MISS | WT | WT | NULL | WT |
| | | | R342* | | | R175H | | | X126_splice | |
| ERpHER2n | ERpHER2n | ERpHER2n | ERnHER2n | ERnHER2n | ERpHER2n | ERpHER2n | HER2p | ERpHER2n | ERnHER2n | ERpHER2n |
| Died of Other Causes | Died of Other Causes | Died of Disease | Living | Died of Disease | Living | Died of Disease | Living | Living | Died of Disease | Died of Other Causes |
| 75.96666667 | 64.3 | 89.1 | 278.2666667 | 52.96666667 | 279.1 | 44.76666667 | 193.7 | 75.86666667 | 89.53333333 | 197.7333333 |
| NO | NO | NO | YES | NO | NO | NO | NO | NO | YES | NO |

| | | | | | | | | |
|---|---|---|---|---|---|---|---|---|
| MB_6187 | MB_6248 | MB_6330 | MB_6184 | MB_6329 | MB_6283 | MB_6359 | MB_6337 | MB_6256 |
| WT | MUT | WT | MUT | MUT | WT | MUT | MUT | WT |
| WT | MISS | WT | MISS | MISS | WT | MISS | MISS | WT |
| | R248W | | R280K | R337H | | Q317R | G266V | |
| ERpHER2n | ERnHER2n | HER2p | ERpHER2n | ERpHER2n | ERpHER2n | ERpHER2n | HER2p | ERpHER2n |
| Died of Other Causes | Living | Living | Died of Disease | Died of Other Causes | Died of Other Causes | Died of Disease | Died of Other Causes | Died of Other Causes |
| 37.76666667 | 182.8333333 | 140.3333333 | 185.7 | 81.5 | 92.4 | 85.03333333 | 110.8666667 | 127.8333333 |
| NO | YES | YES | YES | NO | NO | NO | YES | NO |

| | | | | | | | | |
|---|---|---|---|---|---|---|---|---|
| MB_6201 | MB_6207 | MB_6194 | MB_6281 | MB_6322 | MB_6223 | MB_6190 | MB_6306 | MB_6188 |
| WT | WT | MUT | MUT | WT | MUT | WT | WT | MUT |
| WT | WT | MISS | NULL | WT | MISS | WT | WT | MISS |
| | | R280K | L265Gfs*34 | | G245D | | | A159P |
| ERpHER2n | ERpHER2n | ERnHER2n | ERpHER2n | ERpHER2n | ERnHER2n | ERpHER2n | ERpHER2n | ERnHER2n |
| Died of Other Causes | Died of Other Causes | Died of Disease | Died of Other Causes | Died of Other Causes | Living | Died of Other Causes | Died of Other Causes | Living |
| 222.7 | 118.3 | 52.46666667 | 3.5 | 80.36666667 | 221.9333333 | 24.63333333 | 64.6 | 23.76666667 |
| NO | NO | YES | NO | NO | YES | NO | NO | NO |

| | | | | | | | | | |
|---|---|---|---|---|---|---|---|---|---|
| MB_6318 | MB_6228 | MB_6226 | MB_6225 | MB_6280 | MB_6253 | MB_6098 | MB_6017 | MB_6019 | MB_6163 |
| MUT | MUT | WT | MUT | MUT | WT | MUT | WT | MUT | WT |
| NULL | MISS | WT | NULL | NULL | WT | NULL | WT | MISS | WT |
| Q331* | R249S | | I254Sfs*91 | X126_splice | | S215Cfs*6 | | R248Q | |
| ERnHER2n | ERnHER2n | ERpHER2n | ERpHER2n | ERnHER2n | ERpHER2n | ERnHER2n | ERpHER2n | ERpHER2n | ERpHER2n |
| Died of Other Causes | Living | Died of Other Causes | Living | Died of Disease | Living | Died of Disease | Died of Other Causes | Died of Other Causes | Died of Other Causes |
| 109.2333333 | 200.1 | 35.36666667 | 117.8666667 | 27.86666667 | 194.3666667 | 10.63333333 | 70.9 | 13.53333333 | 88.66666667 |
| NO | NO | NO | NO | YES | NO | YES | NO | NO | NO |

| | | | | | | | | |
|---|---|---|---|---|---|---|---|---|
| MB_6157 | MB_6179 | MB_6105 | MB_6047 | MB_6001 | MB_6050 | MB_6169 | MB_4548 | MB_4564 |
| MUT | MUT | WT | MUT | MUT | WT | MUT | MUT | WT |
| MISS | MISS | WT | MISS | MISS | WT | MISS | MISS | WT |
| R248W | G245S | | C242F | F270L | | H193Y | C238R | |
| HER2p | ERpHER2n | HER2p | ERpHER2n | ERpHER2n | ERpHER2n | HER2p | HER2p | ERpHER2n |
| Died of Disease | Died of Disease | Died of Other Causes | Died of Disease | Died of Other Causes | Living | Died of Other Causes | Living | Died of Other Causes |
| 15.86666667 | 58.46666667 | 192.2 | 37.86666667 | 135.3 | 192.2 | 84.73333333 | 322.8333333 | 210.4333333 |
| NO | NO | NO | NO | NO | NO | NO | YES | NO |

| | | | | | | | | | |
|---|---|---|---|---|---|---|---|---|---|
| MB_4591 | MB_4557 | MB_4601 | MB_4598 | MB_4578 | MB_4593 | MB_5186 | MB_4974 | MB_4978 | MB_5086 |
| WT | WT | MUT | WT | WT | WT | MUT | MUT | WT | WT |
| WT | WT | NULL | WT | WT | WT | NULL | NULL | WT | WT |
| | | X261_splice | | | | R196* | G108Vfs*15 | | |
| ERpHER2n | ERpHER2n | HER2p | ERpHER2n | ERpHER2n | ERpHER2n | HER2p | ERnHER2n | ERpHER2n | ERpHER2n |
| Died of Other Causes | Living | Died of Disease | Died of Disease | Died of Other Causes | Living | Living | Died of Other Causes | Living | Died of Other Causes |
| 41.16666667 | 279.8 | 56.26666667 | 119.3666667 | 197.3333333 | 140.2333333 | 203.2666667 | 176.1 | 198.4333333 | 84.23333333 |
| NO | NO | YES | NO | NO | NO | NO | NO | NO | NO |

| | | | | | | | | |
|---|---|---|---|---|---|---|---|---|
| MB_5150 | MB_5345 | MB_4696 | MB_4695 | MB_4714 | MB_4705 | MB_4882 | MB_4879 | MB_4881 |
| WT | WT | MUT | WT | MUT | WT | WT | MUT | MUT |
| WT | WT | NULL | WT | MISS | WT | WT | MISS | MISS |
| | | X307_splice | | R248Q | | | P278S | C242Y |
| ERpHER2n | ERpHER2n | ERnHER2n | ERpHER2n | ERnHER2n | ERpHER2n | ERpHER2n | HER2p | ERnHER2n |
| Died of Disease | Died of Disease | Living | Died of Other Causes | Died of Other Causes | Died of Other Causes | Died of Other Causes | Died of Other Causes | Living |
| 110.8333333 | 216.0333333 | 255.2666667 | 21.3 | 83.66666667 | 213.1 | 266.1 | 79.96666667 | 274.2 |
| NO | NO | NO | NO | YES | NO | NO | NO | NO |

| | | | | | | | | | | |
|---|---|---|---|---|---|---|---|---|---|---|
| MB_4846 | MB_4870 | MB_4224 | MB_4845 | MB_4904 | MB_4865 | MB_4876 | MB_4860 | MB_5081 | MB_4897 | MB_4925 |
| WT | WT | WT | WT | WT | MUT | WT | MUT | MUT | WT | WT |
| WT | WT | WT | WT | WT | MISS | WT | MISS | MISS | WT | WT |
| | | | | | R175H | | R175H | V173M | | |
| HER2p | ERpHER2n | ERpHER2n | ERpHER2n | ERnHER2n | ERnHER2n | ERpHER2n | ERpHER2n | ERpHER2n | ERpHER2n | ERpHER2n |
| Died of Disease | Living | Died of Disease | Died of Disease | Died of Other Causes | Living | Died of Disease | Living | Died of Disease | Living | Died of Other Causes |
| 101.6333333 | 229.0666667 | 65.5 | 25.46666667 | 129.2333333 | 229.9 | 150.6 | 234.6 | 58 | 197.4333333 | 102.7666667 |
| NO | NO | NO | NO | YES | NO | NO | NO | NO | NO | NO |

| | | | | | | | | | |
|---|---|---|---|---|---|---|---|---|---|
| MB_4883 | MB_4896 | MB_4987 | MB_4416 | MB_4374 | MB_4407 | MB_5039 | MB_4119 | MB_4091 | MB_5079 |
| WT | MUT | WT | MUT | WT | MUT | WT | WT | WT | WT |
| WT | MISS | WT | NULL | WT | MISS | WT | WT | WT | WT |
| | G245V | | V122Cfs*27 | | R248Q | | | | |
| ERpHER2n | HER2p | ERpHER2n | ERnHER2n | ERpHER2n | ERnHER2n | HER2p | ERpHER2n | ERpHER2n | ERpHER2n |
| Living | Died of Disease | Died of Other Causes | Died of Disease | Died of Other Causes | Died of Disease | Died of Disease | Died of Disease | Died of Other Causes | Living |
| 268.9 | 64.86666667 | 221.7666667 | 241.6 | 42.56666667 | 83.36666667 | 62.13333333 | 67.8 | 281.3666667 | 199.9333333 |
| NO | NO | NO | YES | NO | NO | NO | NO | NO | NO |

| | | | | | | | | | |
|---|---|---|---|---|---|---|---|---|---|
| MB_5073 | MB_5105 | MB_4254 | MB_5174 | MB_4742 | MB_5239 | MB_5251 | MB_4996 | MB_4282 | MB_4603 |
| WT | WT | MUT | MUT | WT | WT | WT | WT | MUT | WT |
| WT | WT | NULL | MISS | WT | WT | WT | WT | NULL | WT |
| | | R282Gfs*63 | G245V | | | | | Q317* | |
| ERpHER2n | ERpHER2n | ERnHER2n | HER2p | ERpHER2n | ERpHER2n | ERpHER2n | ERpHER2n | ERpHER2n | ERpHER2n |
| Died of Other Causes | Died of Disease | Living | Died of Other Causes | Died of Other Causes | Living | Died of Disease | Died of Other Causes | Living | Living |
| 194.1666667 | 204.4333333 | 178.4 | 151.2 | 83.13333333 | 192.1333333 | 112.4666667 | 99.76666667 | 34.1 | 239.2333333 |
| NO | NO | NO | NO | NO | NO | NO | NO | NO | NO |

| | | | | | | | | | | |
|---|---|---|---|---|---|---|---|---|---|---|
| MB_4621 | MB_4853 | MB_4362 | MB_4602 | MB_4222 | MB_4893 | MB_4859 | MB_4212 | MB_4873 | MB_4942 | MB_4949 |
| MUT | WT | WT | WT | WT | MUT | WT | WT | WT | MUT | MUT |
| NULL | WT | WT | WT | WT | NULL | WT | WT | WT | NULL | MISS |
| R213* | | | | | R213* | | | | X126_splice | R175H |
| ERnHER2n | ERpHER2n | ERpHER2n | ERpHER2n | ERpHER2n | ERnHER2n | ERnHER2n | ERpHER2n | ERpHER2n | ERnHER2n | ERpHER2n |
| Living | Living | Living | Died of Disease | Died of Disease | Living | Died of Other Causes | Living | Died of Disease | Died of Other Causes | Living |
| 271.8666667 | 265.9333333 | 267.2666667 | 122.8 | 55.23333333 | 176.7 | 157.8 | 330.3666667 | 117.6666667 | 124.1333333 | 216.9666667 |
| YES | NO | NO | NO | NO | YES | NO | NO | NO | NO | NO |

| | | | | | | | | |
|---|---|---|---|---|---|---|---|---|
| MB_4928 | MB_5098 | MB_5313 | MB_4145 | MB_5122 | MB_5278 | MB_4213 | MB_4126 | MB_4957 |
| MUT | WT | MUT | MUT | WT | WT | WT | MUT | WT |
| NULL | WT | MISS | MISS | WT | WT | WT | MISS | WT |
| V157_R158del | | R273G | H179Q | | | | Y220C | |
| ERnHER2n | ERpHER2n | ERpHER2n | ERpHER2n | ERpHER2n | ERpHER2n | ERpHER2n | ERpHER2n | ERpHER2n |
| Living | Died of Other Causes | Died of Disease | Died of Other Causes | Died of Other Causes | Died of Disease | Died of Disease | Died of Other Causes | Died of Disease |
| 220.0333333 | 186.4 | 46.53333333 | 217.6333333 | 125.7 | 99.96666667 | 84.5 | 174.8 | 197.6666667 |
| NO | NO | NO | NO | NO | NO | NO | NO | NO |

| | | | | | | | | | |
|---|---|---|---|---|---|---|---|---|---|
| MB_5470 | MB_5317 | MB_5308 | MB_4976 | MB_5277 | MB_5110 | MB_4046 | MB_4012 | MB_4141 | MB_4001 |
| WT | WT | WT | WT | WT | WT | WT | WT | WT | WT |
| WT | WT | WT | WT | WT | WT | WT | WT | WT | WT |
| ERpHER2n | ERpHER2n | ERpHER2n | ERpHER2n | ERpHER2n | ERpHER2n | ERpHER2n | HER2p | ERpHER2n | ERpHER2n |
| Died of Disease | Died of Disease | Died of Disease | Died of Other Causes | Died of Disease | Died of Disease | Died of Other Causes | Died of Other Causes | Died of Disease | Died of Disease |
| 42.06666667 | 53.63333333 | 104.7666667 | 224.1 | 221.0666667 | 21.9 | 298.0333333 | 277.3666667 | 147.8333333 | 49.66666667 |
| NO | NO | NO | NO | NO | NO | NO | NO | NO | NO |

| | | | | | | | | | |
|---|---|---|---|---|---|---|---|---|---|
| MB_4792 | MB_4738 | MB_4264 | MB_4794 | MB_4782 | MB_5008 | MB_4426 | MB_4390 | MB_4434 | MB_4368 |
| MUT | WT | WT | WT | WT | MUT | MUT | WT | WT | WT |
| MISS | WT | WT | WT | WT | NULL | NULL | WT | WT | WT |
| G245C | | | | | X225_splice | P153Afs*28 | | | |
| ERnHER2n | ERpHER2n | ERpHER2n | ERpHER2n | ERpHER2n | ERnHER2n | ERpHER2n | ERpHER2n | ERpHER2n | ERpHER2n |
| Died of Disease | Living | Died of Other Causes | Died of Other Causes | Living | Living | Died of Other Causes | Died of Other Causes | Died of Disease | Died of Disease |
| 40.63333333 | 176.0666667 | 163.2 | 45.5 | 251.0666667 | 223.3 | 170.9 | 164.3333333 | 148.1 | 119.4666667 |
| YES | NO | NO | NO | NO | YES | NO | NO | NO | NO |

| | | | | | | | | | | |
|---|---|---|---|---|---|---|---|---|---|---|
| MB_4442 | MB_4418 | MB_4381 | MB_4395 | MB_4630 | MB_6010 | MB_6021 | MB_6059 | MB_6154 | MB_6097 | MB_6107 |
| WT | WT | WT | WT | WT | MUT | WT | MUT | WT | WT | WT |
| WT | WT | WT | WT | WT | MISS | WT | MISS | WT | WT | WT |
| | | | | | R175H | | R175G | | | |
| ERpHER2n | ERpHER2n | ERpHER2n | ERpHER2n | ERpHER2n | ERpHER2n | ERpHER2n | ERpHER2n | ERpHER2n | ERpHER2n | ERpHER2n |
| Living | Living | Died of Disease | Died of Disease | Died of Other Causes | Living | Living | Living | Living | Died of Other Causes | Died of Other Causes |
| 296.8666667 | 307.6333333 | 61.46666667 | 139.6 | 95.83333333 | 218.2333333 | 301.2333333 | 278.4666667 | 195.3 | 159.7 | 123.3333333 |
| NO | NO | NO | NO | NO | NO | NO | YES | NO | NO | NO |

| | | | | | | | | | |
|---|---|---|---|---|---|---|---|---|---|
| MB_6023 | MB_6152 | MB_6018 | MB_6082 | MB_6029 | MB_5295 | MB_5386 | MB_5092 | MB_6068 | MB_6039 |
| WT | MUT | WT | WT | WT | MUT | WT | WT | MUT | MUT |
| WT | MISS | WT | WT | WT | MISS | WT | WT | NULL | NULL |
| | P278S | | | | G245D | | | E294* | X307_splice |
| ERpHER2n | ERnHER2n | ERpHER2n | HER2p | ERpHER2n | ERnHER2n | ERpHER2n | ERpHER2n | ERnHER2n | ERpHER2n |
| Living | Living | Died of Disease | Died of Disease | Died of Disease | Died of Other Causes | Living | Died of Other Causes | Died of Other Causes | Died of Other Causes |
| 232.9666667 | 228.9 | 143.1666667 | 102.5 | 278.3666667 | 75.7 | 192.2 | 174.5666667 | 182.6 | 73.13333333 |
| NO | NO | NO | YES | NO | NO | NO | NO | NO | NO |

| | | | | | | | | | |
|---|---|---|---|---|---|---|---|---|---|
| MB_6022 | MB_6125 | MB_6053 | MB_6135 | MB_6042 | MB_6058 | MB_6077 | MB_6052 | MB_5368 | MB_4189 |
| WT | WT | MUT | MUT | WT | MUT | WT | WT | WT | MUT |
| WT | WT | NULL | NULL | WT | MISS | WT | WT | WT | MISS |
| | | C135Afs*35 | K305Sfs*40 | | Y220C | | | | G245S |
| ERpHER2n | ERpHER2n | ERpHER2n | ERpHER2n | ERpHER2n | ERnHER2n | ERpHER2n | ERnHER2n | ERpHER2n | ERpHER2n |
| Died of Disease | Died of Other Causes | Living | Died of Other Causes | Living | Died of Disease | Died of Other Causes | Living | Died of Other Causes | Died of Other Causes |
| 52.06666667 | 153.9 | 250.8 | 78.6 | 263.4333333 | 22.66666667 | 170.2666667 | 255.3666667 | 100.1333333 | 355.2 |
| NO | NO | NO | NO | NO | YES | NO | NO | NO | NO |

| | | | | | | | | | | | |
|---|---|---|---|---|---|---|---|---|---|---|---|
| MB_5335 | MB_4008 | MB_5134 | MB_4139 | MB_5130 | MB_5135 | MB_4880 | MB_5054 | MB_5057 | MB_4098 | MB_4760 | MB_5102 |
| MUT | WT | WT | WT | MUT | MUT | MUT | MUT | MUT | WT | WT | MUT |
| NULL | WT | WT | WT | MISS | NULL | NULL | MISS | NULL | WT | WT | MISS |
| T125Vfs*23 | | | | R280T | G108Vfs*13 | Y107* | R248Q | N239* | | | Y220C |
| ERnHER2n | ERpHER2n | ERpHER2n | ERpHER2n | ERpHER2n | ERnHER2n | ERnHER2n | HER2p | ERnHER2n | ERpHER2n | ERpHER2n | ERnHER2n |
| Died of Disease | Died of Disease | Living | Died of Other Causes | NA | Died of Disease | Living | Living | Died of Disease | Living | Died of Disease | Died of Disease |
| 25.23333333 | 44.73333333 | 196.5333333 | 165.4 | 255 | 37.36666667 | 210.9666667 | 187.7 | 56.33333333 | 176.3666667 | 124.8 | 30.36666667 |
| NO | NO | NO | NO | NO | NO | NO | NO | NO | NO | NO | YES |

| | | | | | | | | | | | |
|---|---|---|---|---|---|---|---|---|---|---|---|
| MB_5420 | MB_5458 | MB_5492 | MB_5407 | MB_5394 | MB_5410 | MB_5435 | MB_5406 | MB_4329 | MB_4005 | MB_6288 | MB_6272 |
| WT | MUT | WT | WT | WT | WT | WT | WT | WT | MUT | MUT | MUT |
| WT | MISS | WT | WT | WT | WT | WT | WT | WT | MISS | NULL | NULL |
| | E285K | | | | | | | | R175H | E198* | T125T |
| HER2p | HER2p | ERpHER2n | ERpHER2n | HER2p | ERpHER2n | ERpHER2n | ERpHER2n | ERpHER2n | ERpHER2n | ERpHER2n | ERnHER2n |
| Living | Living | Died of Disease | Living | Living | Living | Died of Other Causes | Died of Disease | Died of Disease | Living | Died of Disease | Died of Disease |
| 193.9 | 183.4333333 | 23.36666667 | 205.6 | 193.6 | 202.7666667 | 104 | 97.43333333 | 168.2666667 | 188.3666667 | 100.3 | 22.4 |
| NO | YES | NO | NO | NO | NO | NO | NO | NO | NO | NO | YES |

| | | | | | | | | | |
|---|---|---|---|---|---|---|---|---|---|
| MB_6218 | MB_6246 | MB_6217 | MB_6308 | MB_6271 | MB_6245 | MB_6233 | MB_4343 | MB_4281 | MB_4283 |
| WT | MUT | WT | WT | MUT | WT | WT | MUT | MUT | WT |
| WT | NULL | WT | WT | MISS | WT | WT | NULL | MISS | WT |
| | P322Hfs*23 | | | C135F | | | R342* | L194R | |
| ERpHER2n | HER2p | ERpHER2n | ERpHER2n | ERpHER2n | ERnHER2n | ERpHER2n | ERpHER2n | ERpHER2n | ERpHER2n |
| Died of Disease | Died of Other Causes | Died of Other Causes | Died of Other Causes | Living | Living | Living | Died of Other Causes | Died of Other Causes | Living |
| 147.7666667 | 52.9 | 211.5333333 | 151.9333333 | 157.7333333 | 177.6333333 | 201.1666667 | 198.6 | 260.0333333 | 174.8333333 |
| NO | YES | YES | NO | NO | NO | NO | NO | NO | NO |

| | | | | | | | | | |
|---|---|---|---|---|---|---|---|---|---|
| MB_4341 | MB_4303 | MB_4274 | MB_4292 | MB_4278 | MB_5399 | MB_5364 | MB_3526 | MB_6138 | MB_6049 |
| MUT | MUT | WT | WT | WT | WT | WT | MUT | WT | MUT |
| NULL | MISS | WT | WT | WT | WT | WT | MISS | WT | NULL |
| W146* | D281A | | | | | | R282W | | R196* |
| HER2p | ERnHER2n | ERpHER2n | ERpHER2n | ERpHER2n | ERpHER2n | ERpHER2n | ERpHER2n | ERpHER2n | HER2p |
| Died of Disease | Died of Other Causes | Died of Disease | Died of Other Causes | Died of Other Causes | Died of Disease | Died of Disease | Died of Disease | Died of Disease | Living |
| 33.7 | 89.03333333 | 59.93333333 | 335.6 | 244.2 | 106.8 | 125.7666667 | 86.06666667 | 25.63333333 | 264.2333333 |
| YES | NO | NO | NO | NO | NO | NO | NO | NO | NO |

| | | | | | | | | | |
|---|---|---|---|---|---|---|---|---|---|
| MB_6055 | MB_6108 | MB_6044 | MB_6181 | MB_6100 | MB_6085 | MB_6113 | MB_6079 | MB_6103 | MB_6016 |
| MUT | WT | MUT | WT | MUT | WT | MUT | WT | WT | WT |
| MISS | WT | MISS | WT | NULL | WT | MISS | WT | WT | WT |
| R248W | | G334V | | G302Rfs*4 | | D259Y | | | |
| ERnHER2n | ERpHER2n | ERpHER2n | ERpHER2n | HER2p | HER2p | HER2p | ERpHER2n | ERpHER2n | ERpHER2n |
| Living | Living | Died of Other Causes | Died of Other Causes | Died of Disease | Living | Died of Disease | Died of Disease | Living | Died of Other Causes |
| 200.4333333 | 193.9666667 | 219.1 | 54.93333333 | 10.06666667 | 23.9 | 124.5666667 | 239.1666667 | 239.4 | 143.5333333 |
| YES | NO | NO | NO | YES | YES | YES | NO | NO | NO |

| | | | | | | | | | |
|---|---|---|---|---|---|---|---|---|---|
| MB_6149 | MB_6131 | MB_6007 | MB_6147 | MB_6114 | MB_6167 | MB_6146 | MB_6092 | MB_6160 | MB_6075 |
| WT | MUT | MUT | WT | MUT | WT | MUT | WT | MUT | WT |
| WT | NULL | MISS | WT | MISS | WT | NULL | WT | MISS | WT |
| | X224_splice | R175H | | Y234C | | R158Pfs*12 | | R248G | |
| ERpHER2n | HER2p | ERpHER2n | ERpHER2n | HER2p | ERpHER2n | ERpHER2n | ERpHER2n | HER2p | ERpHER2n |
| Died of Other Causes | Living | Died of Other Causes | Died of Other Causes | Living | Died of Other Causes | Died of Disease | Died of Other Causes | Living | Died of Disease |
| 74.93333333 | 257.6666667 | 70.53333333 | 238.0666667 | 66.83333333 | 122.8 | 89.6 | 2.3 | 247.3666667 | 182.3333333 |
| NO | NO | NO | NO | YES | NO | NO | NO | YES | NO |

| | | | | | | | | | | |
|---|---|---|---|---|---|---|---|---|---|---|
| MB_6011 | MB_6071 | MB_6214 | MB_6334 | MB_6346 | MB_6319 | MB_6200 | MB_6336 | MB_6211 | MB_6232 | MB_6234 |
| WT | MUT | WT | MUT | WT | WT | MUT | MUT | WT | WT | WT |
| WT | MISS | WT | NULL | WT | WT | NULL | MISS | WT | WT | WT |
| | R110H | | E198* | | | R213* | R273H | | | |
| ERpHER2n | ERpHER2n | ERpHER2n | HER2p | ERpHER2n | ERpHER2n | ERpHER2n | ERnHER2n | ERpHER2n | ERpHER2n | ERpHER2n |
| Died of Disease | Died of Disease | Died of Other Causes | Living | Living | Living | Died of Disease | Died of Disease | Died of Other Causes | Died of Other Causes | Living |
| 211.1333333 | 57.4 | 174.5 | 128.1 | 281.5 | 154.7 | 30.06666667 | 17.2 | 42.3 | 85 | 213.3666667 |
| NO | NO | NO | NO | NO | YES | NO | YES | NO | NO | NO |

| | | | | | | | | |
|---|---|---|---|---|---|---|---|---|
| MB_6300 | MB_5533 | MB_5543 | MB_4236 | MB_4234 | MB_4270 | MB_4235 | MB_4250 | MB_5467 |
| WT | MUT | MUT | WT | WT | MUT | WT | MUT | WT |
| WT | NULL | NULL | WT | WT | MISS | WT | MISS | WT |
| | E171* | Y107_F109del | | | R175H | | R175H | |
| ERpHER2n | ERpHER2n | ERpHER2n | ERpHER2n | ERpHER2n | HER2p | ERpHER2n | ERpHER2n | ERpHER2n |
| Died of Other Causes | Living | Died of Other Causes | Died of Other Causes | Died of Other Causes | Died of Disease | Died of Disease | Died of Other Causes | Living |
| 63.83333333 | 158.0333333 | 34.43333333 | 203.7 | 253.5333333 | 98.7 | 335.7333333 | 100.1666667 | 186.1 |
| NO | NO | NO | NO | NO | YES | NO | NO | NO |

| | | | | | | | | | |
|---|---|---|---|---|---|---|---|---|---|
| MB_4323 | MB_4318 | MB_4333 | MB_4339 | MB_4322 | MB_4351 | MB_5063 | MB_4360 | MB_4375 | MB_4798 |
| WT | WT | MUT | WT | WT | MUT | MUT | MUT | WT | WT |
| WT | WT | MISS | WT | WT | MISS | MISS | MISS | WT | WT |
| | | R175H | | | V157F | Y163C | G262V | | |
| ERpHER2n | ERpHER2n | ERpHER2n | ERpHER2n | ERpHER2n | ERnHER2n | HER2p | HER2p | ERpHER2n | ERpHER2n |
| Died of Other Causes | Died of Other Causes | Died of Other Causes | Died of Disease | Died of Other Causes | Died of Disease | Living | Died of Disease | Living | Living |
| 126.6333333 | 17.13333333 | 207.9666667 | 173.6 | 196.4666667 | 68.13333333 | 215.1666667 | 11.86666667 | 291.1666667 | 257.1666667 |
| NO | NO | NO | NO | NO | NO | YES | YES | NO | NO |

| | | | | | | | | | | |
|---|---|---|---|---|---|---|---|---|---|---|
| MB_4746 | MB_4838 | MB_4829 | MB_4802 | MB_5426 | MB_5444 | MB_5431 | MB_5447 | MB_5434 | MB_5231 | MB_5339 |
| WT | MUT | WT | WT | MUT | WT | MUT | WT | WT | MUT | WT |
| WT | MISS | WT | WT | NULL | WT | NULL | WT | WT | NULL | WT |
| | C238S | | | X187_splice | | A83Pfs*35 | | | S149Pfs*21 | |
| ERpHER2n | ERpHER2n | ERpHER2n | ERpHER2n | HER2p | ERpHER2n | ERpHER2n | ERpHER2n | ERpHER2n | HER2p | ERpHER2n |
| Living | Died of Other Causes | Died of Other Causes | Died of Disease | Died of Disease | Living | Died of Disease | Died of Disease | Died of Disease | Died of Disease | Living |
| 244.4333333 | 200.7666667 | 176.2666667 | 43.9 | 34.63333333 | 240.4333333 | 15.86666667 | 148.8666667 | 45.16666667 | 16.56666667 | 195.8666667 |
| YES | NO | NO | NO | YES | NO | NO | NO | NO | NO | NO |

| | | | | | | | | | |
|---|---|---|---|---|---|---|---|---|---|
| MB_5208 | MB_5428 | MB_5440 | MB_5425 | MB_5298 | MB_5334 | MB_5326 | MB_4059 | MB_6230 | MB_6150 |
| MUT | WT | MUT | WT | MUT | WT | WT | WT | WT | WT |
| MISS | WT | MISS | WT | MISS | WT | WT | WT | WT | WT |
| S127F | | N239D | | M246R | | | | | |
| ERnHER2n | ERpHER2n | ERnHER2n | ERpHER2n | ERnHER2n | ERpHER2n | ERpHER2n | ERpHER2n | ERpHER2n | ERpHER2n |
| Living | Living | Living | Died of Disease | Died of Disease | Died of Other Causes | Died of Other Causes | Died of Other Causes | Died of Disease | Died of Other Causes |
| 151.9 | 150.4666667 | 196.6333333 | 49.46666667 | 84.9 | 85.4 | 96.2 | 172 | 74.8 | 77.66666667 |
| NO | NO | YES | NO | YES | NO | NO | NO | NO | NO |

| | | | | | | | | | |
|---|---|---|---|---|---|---|---|---|---|
| MB_6014 | MB_6145 | MB_6012 | MB_6024 | MB_6124 | MB_6171 | MB_6060 | MB_6008 | MB_6036 | MB_6144 |
| WT | WT | WT | WT | WT | WT | WT | WT | MUT | MUT |
| WT | WT | WT | WT | WT | WT | WT | WT | MISS | MISS |
| | | | | | | | | R175H | R175H |
| ERpHER2n | ERpHER2n | ERpHER2n | ERpHER2n | ERpHER2n | ERpHER2n | ERpHER2n | ERpHER2n | HER2p | ERnHER2n |
| Died of Other Causes | Died of Disease | Died of Other Causes | Died of Disease | Died of Other Causes | Living | Died of Other Causes | Died of Disease | Living | Living |
| 191.9333333 | 34.33333333 | 144.4333333 | 252.2666667 | 115.6333333 | 220.3 | 38.03333333 | 69.1 | 287.9333333 | 254.9333333 |
| NO | NO | NO | NO | NO | NO | NO | NO | NO | NO |

| | | | | | | | | | |
|---|---|---|---|---|---|---|---|---|---|
| MB_5346 | MB_5373 | MB_6178 | MB_6063 | MB_6062 | MB_6069 | MB_6141 | MB_6118 | MB_6065 | MB_6006 |
| MUT | WT | MUT | WT | MUT | WT | WT | WT | WT | WT |
| MISS | WT | NULL | WT | MISS | WT | WT | WT | WT | WT |
| V173G | | E51Lfs*73 | | E285K | | | | | |
| ERnHER2n | ERpHER2n | ERnHER2n | HER2p | ERnHER2n | ERpHER2n | ERpHER2n | ERpHER2n | ERpHER2n | ERpHER2n |
| Living | Living | Died of Disease | Died of Disease | Living | Died of Other Causes | Died of Other Causes | Living | Died of Other Causes | Died of Other Causes |
| 194.3 | 2.533333333 | 33.13333333 | 15.6 | 282.3666667 | 258.1333333 | 78.86666667 | 264.6 | 188.5333333 | 63.56666667 |
| YES | NO | NO | YES | NO | NO | NO | NO | NO | NO |

| | | | | | | | | | |
|---|---|---|---|---|---|---|---|---|---|
| MB_6048 | MB_6116 | MB_4310 | MB_4300 | MB_4417 | MB_4079 | MB_4317 | MB_4350 | MB_4332 | MB_4324 |
| MUT | MUT | WT | MUT | MUT | MUT | MUT | MUT | MUT | WT |
| NULL | NULL | WT | NULL | MISS | MISS | MISS | NULL | MISS | WT |
| C124Afs*46 | E62* | | R209Kfs*6 | R280T | V274L | A159V | R306* | K132N | |
| HER2p | HER2p | ERpHER2n | ERpHER2n | ERnHER2n | ERpHER2n | HER2p | HER2p | ERnHER2n | ERpHER2n |
| Died of Other Causes | Living | Died of Other Causes | Living | Died of Other Causes | Died of Other Causes | Living | Died of Disease | Living | Died of Disease |
| 168.2 | 193.1333333 | 265.5666667 | 189.0333333 | 16.7 | 351 | 177.9 | 31.06666667 | 307.9333333 | 87.23333333 |
| NO | NO | NO | NO | NO | NO | NO | YES | NO | NO |

| | | | | | | | | | | | |
|---|---|---|---|---|---|---|---|---|---|---|---|
| MB_4306 | MB_7226 | MB_7205 | MB_7160 | MB_7159 | MB_7167 | MB_7207 | MB_7237 | MB_7241 | MB_7236 | MB_7168 | MB_7162 |
| WT | WT | MUT | MUT | MUT | WT | MUT | WT | WT | WT | WT | WT |
| WT | WT | MISS | MISS | NULL | WT | MISS | WT | WT | WT | WT | WT |
| | | R273H | H179D | T102Ifs*46 | | P250L | | | | | |
| ERpHER2n | ERpHER2n | ERnHER2n | ERpHER2n | ERnHER2n | ERpHER2n | HER2p | ERpHER2n | ERpHER2n | ERpHER2n | ERpHER2n | ERpHER2n |
| Died of Other Causes | Died of Disease | Living | Living | Died of Disease | Living | Died of Disease | Died of Disease | Living | Died of Other Causes | Living | Living |
| 234.7 | 53.36666667 | 166.0333333 | 148.0666667 | 21.93333333 | 141.0333333 | 55.73333333 | 23.93333333 | 171.1 | 201.4666667 | 150.5 | 140.2 |
| NO | YES | YES | NO | YES | NO | YES | NO | NO | NO | NO | NO |

| | | | | | | | | |
|---|---|---|---|---|---|---|---|---|
| MB_7235 | MB_4331 | MB_4353 | MB_4354 | MB_4328 | MB_4348 | MB_6030 | MB_6133 | MB_6083 |
| WT | WT | WT | WT | MUT | WT | WT | WT | WT |
| WT | WT | WT | WT | MISS | WT | WT | WT | WT |
| | | | | P151T | | | | |
| ERpHER2n | ERpHER2n | ERpHER2n | ERnHER2n | ERpHER2n | HER2p | ERpHER2n | ERpHER2n | ERpHER2n |
| Living | Died of Other Causes | Died of Disease | Died of Disease | Died of Other Causes | Died of Disease | Died of Other Causes | Died of Disease | Died of Other Causes |
| 170.8 | 235.5666667 | 128.5333333 | 15.36666667 | 199.2666667 | 226.2 | 145.3666667 | 61.8 | 164.0333333 |
| NO | NO | NO | NO | NO | NO | NO | NO | NO |

| | | | | | | | | | | |
|---|---|---|---|---|---|---|---|---|---|---|
| MB_6026 | MB_6168 | MB_6101 | MB_6156 | MB_6051 | MB_6182 | MB_6080 | MB_6143 | MB_6164 | MB_0551 | MB_0391 |
| MUT | WT | WT | MUT | WT | MUT | MUT | MUT | WT | WT | WT |
| MISS | WT | WT | MISS | WT | MISS | MISS | MISS | WT | WT | WT |
| C176S | | | F134C | | C238Y | R175H | R248Q | | | |
| ERpHER2n | ERpHER2n | ERpHER2n | HER2p | ERpHER2n | HER2p | ERpHER2n | ERnHER2n | ERpHER2n | ERpHER2n | HER2p |
| Died of Other Causes | Died of Other Causes | Died of Other Causes | Died of Disease | Died of Disease | Living | Died of Disease | Living | Living | Died of Disease | Living |
| 49.73333333 | 225.7 | 77.5 | 58.93333333 | 186.8333333 | 208.8 | 54.1 | 255.6 | 205.9 | 51.96666667 | 142.4666667 |
| NO | NO | NO | NO | NO | NO | NO | NO | NO | NO | YES |

| | | | | | | | | | | |
|---|---|---|---|---|---|---|---|---|---|---|
| MB_0367 | MB_5171 | MB_5199 | MB_5177 | MB_5178 | MB_5280 | MB_5148 | MB_4342 | MB_5225 | MB_4024 | MB_6314 |
| WT | WT | MUT | WT | WT | WT | WT | WT | MUT | MUT | WT |
| WT | WT | MISS | WT | WT | WT | WT | WT | NULL | MISS | WT |
| | | V173L | | | | | | Q52Pfs*5 | G244C | |
| ERpHER2n | ERpHER2n | HER2p | ERpHER2n | ERpHER2n | ERpHER2n | ERnHER2n | ERpHER2n | ERnHER2n | ERnHER2n | HER2p |
| Living | Died of Disease | Living | Died of Other Causes | Living | Died of Disease | Died of Other Causes | Died of Other Causes | Died of Disease | Living | Living |
| 142.4333333 | 185 | 204.4 | 65.23333333 | 184.3333333 | 77.86666667 | 245.2333333 | 232.7666667 | 43.26666667 | 124.1 | 134.8333333 |
| NO | NO | NO | NO | NO | NO | NO | NO | NO | NO | NO |

| | | | | | | | | | |
|---|---|---|---|---|---|---|---|---|---|
| MB_6257 | MB_6358 | MB_6242 | MB_6237 | MB_6183 | MB_6254 | MB_6231 | MB_6204 | MB_6229 | MB_5154 |
| WT | MUT | MUT | MUT | WT | WT | WT | MUT | WT | MUT |
| WT | NULL | NULL | MISS | WT | WT | WT | MISS | WT | MISS |
| | N345_L348delinsM | X225_splice | R175H | | | | H179R | | G245C |
| ERpHER2n | HER2p | ERnHER2n | ERnHER2n | ERpHER2n | ERpHER2n | ERpHER2n | ERpHER2n | ERpHER2n | HER2p |
| Living | Died of Disease | Living | Died of Other Causes | Died of Other Causes | Died of Disease | Died of Other Causes | Died of Other Causes | Died of Disease | Living |
| 91 | 37.86666667 | 165.4333333 | 105.2 | 79.13333333 | 60.9 | 72.56666667 | 56.76666667 | 0.1 | 170.8333333 |
| NO | NO | YES | NO | NO | NO | NO | NO | NO | NO |

| | | | | | | | | | | |
|---|---|---|---|---|---|---|---|---|---|---|
| MB_5175 | MB_5173 | MB_5162 | MB_5164 | MB_7298 | MB_7299 | MB_7293 | MB_3978 | MB_7288 | MB_2923 | MB_0402 |
| MUT | MUT | WT | WT | WT | WT | MUT | MUT | WT | WT | WT |
| MISS | MISS | WT | WT | WT | WT | MISS | NULL | WT | WT | WT |
| R248Q | R248W | | | | | I255S | N200Ifs*47 | | | |
| ERpHER2n | ERnHER2n | ERnHER2n | HER2p | ERpHER2n | ERpHER2n | ERpHER2n | HER2p | ERpHER2n | HER2p | ERpHER2n |
| Died of Other Causes | Died of Disease | Living | Living | Died of Other Causes | Died of Other Causes | Living | Died of Disease | Died of Disease | Died of Disease | Living |
| 194.7 | 22.13333333 | 164.3333333 | 80.5 | 86.23333333 | 201.9 | 199.2333333 | 100.8666667 | 27.06666667 | 14.1 | 99.73333333 |
| NO | NO | NO | NO | NO | NO | NO | NO | NO | YES | NO |

| | | | | | | | | | | | |
|---|---|---|---|---|---|---|---|---|---|---|---|
| MB_0381 | MB_0592 | MB_5271 | MB_5235 | MB_4931 | MB_5332 | MB_7045 | MB_7049 | MB_7031 | MB_7061 | MB_7050 | MB_7078 |
| MUT | WT | MUT | MUT | MUT | MUT | MUT | WT | MUT | WT | MUT | WT |
| MISS | WT | NULL | MISS | NULL | NULL | NULL | WT | MISS | WT | NULL | WT |
| D208V | | P34Vfs*12 | C141Y | X10_splice | E204* | R306* | | V173L | | Q167* | |
| HER2p | ERpHER2n | ERpHER2n | ERnHER2n | ERnHER2n | HER2p | ERnHER2n | ERnHER2n | ERnHER2n | ERpHER2n | HER2p | ERnHER2n |
| Living | Living | Living | Died of Disease | Died of Disease | Died of Disease | Died of Disease | Living | Living | Died of Other Causes | Died of Disease | Died of Disease |
| 130.2 | 74.56666667 | 167.5 | 204.2 | 30.36666667 | 98.83333333 | 94.93333333 | 102.0333333 | 85.36666667 | 96.76666667 | 65.13333333 | 4.166666667 |
| YES | NO | NO | NO | NO | NO | NO | NO | YES | NO | YES | NO |

| | | | | | | | | | | |
|---|---|---|---|---|---|---|---|---|---|---|
| MB_7054 | MB_7048 | MB_7058 | MB_7056 | MB_7038 | MB_7086 | MB_0177 | MB_0435 | MB_0399 | MB_0388 | MB_0351 |
| MUT | MUT | WT | WT | WT | WT | WT | WT | WT | WT | WT |
| MISS | MISS | WT | WT | WT | WT | WT | WT | WT | WT | WT |
| I195T | R110L | | | | | | | | | |
| ERnHER2n | ERpHER2n | ERpHER2n | ERpHER2n | ERnHER2n | ERpHER2n | ERpHER2n | ERnHER2n | ERnHER2n | HER2p | ERpHER2n |
| Died of Disease | Living | Living | Living | Living | Living | Died of Other Causes | Living | Died of Other Causes | Living | Died of Other Causes |
| 30.43333333 | 88.46666667 | 122.5 | 117 | 81.03333333 | 123.5333333 | 45.6 | 110.5333333 | 135.3333333 | 97.56666667 | 195.3333333 |
| YES | NO | NO | NO | YES | NO | NO | NO | YES | YES | NO |

| | | | | | | | | | | | |
|---|---|---|---|---|---|---|---|---|---|---|---|
| MB_0615 | MB_0265 | MB_0116 | MB_0472 | MB_0191 | MB_0231 | MB_0393 | MB_0553 | MB_0587 | MB_0611 | MB_0640 | MB_0114 |
| MUT | MUT | WT | WT | MUT | WT | WT | WT | MUT | MUT | WT | WT |
| MISS | MISS | WT | WT | MISS | WT | WT | WT | NULL | MISS | WT | WT |
| Y234C | S241Y | | | H193R | | | | R196* | R273C | | |
| HER2p | ERnHER2n | ERpHER2n | ERpHER2n | ERnHER2n | ERpHER2n | ERpHER2n | HER2p | ERpHER2n | ERpHER2n | ERpHER2n | ERpHER2n |
| Living | Living | Living | Living | Died of Other Causes | Living | Died of Disease | Died of Other Causes | Living | Living | Living | Living |
| 34.76666667 | 189.1333333 | 122.2666667 | 27.4 | 15.3 | 19.6 | 20.83333333 | 85.73333333 | 91.26666667 | 104.5333333 | 61.33333333 | 13.4 |
| NO | NO | YES | NO | NO | NO | YES | NO | YES | NO | NO | NO |

| | | | | | | | | | | |
|---|---|---|---|---|---|---|---|---|---|---|
| MB_0222 | MB_0284 | MB_0459 | MB_0128 | MB_0639 | MB_0318 | MB_0417 | MB_6238 | MB_6286 | MB_6328 | MB_6284 |
| WT | MUT | MUT | WT | MUT | WT | WT | WT | MUT | WT | WT |
| WT | MISS | MISS | WT | MISS | WT | WT | WT | MISS | WT | WT |
| | P250L | Y163C | | R248Q | | | | G245C | | |
| ERpHER2n | ERnHER2n | ERpHER2n | ERpHER2n | ERnHER2n | ERnHER2n | ERpHER2n | ERpHER2n | ERpHER2n | ERpHER2n | ERpHER2n |
| Living | Living | Died of Other Causes | Living | Living | Living | Died of Disease | Died of Disease | Died of Other Causes | Living | Died of Other Causes |
| 212.7 | 0 | 35.6 | 137.8 | 75.4 | 168.7 | 6.266666667 | 172.9666667 | 49.53333333 | 126.6666667 | 88.33333333 |
| YES | NO | NO | NO | YES | NO | NO | NO | NO | NO | NO |

| | | | | | | | | | | | |
|---|---|---|---|---|---|---|---|---|---|---|---|
| MB_6363 | MB_6224 | MB_6297 | MB_6239 | MB_6208 | MB_6273 | MB_6344 | MB_6312 | MB_6212 | MB_6302 | MB_6327 | MB_6287 |
| WT | MUT | WT | WT | WT | MUT | MUT | WT | WT | WT | WT | WT |
| WT | NULL | WT | WT | WT | NULL | NULL | WT | WT | WT | WT | WT |
| | A84_S90del | | | | W91* | R306* | | | | | |
| HER2p | HER2p | ERpHER2n | ERpHER2n | ERpHER2n | ERpHER2n | ERpHER2n | ERpHER2n | ERpHER2n | ERpHER2n | ERpHER2n | ERpHER2n |
| Died of Disease | Died of Disease | Living | Living | Living | Living | Died of Disease | Living | Living | Died of Other Causes | Died of Disease | Died of Disease |
| 16.7 | 35.46666667 | 170.8666667 | 200.3333333 | 221.2 | 189.4333333 | 152.9333333 | 162.7666667 | 224.2333333 | 125.8666667 | 18.26666667 | 35.2 |
| YES | YES | NO | YES | NO | YES | NO | NO | NO | NO | NO | YES |

| | | | | | | | | | |
|---|---|---|---|---|---|---|---|---|---|
| MB_6251 | MB_0489 | MB_0342 | MB_7289 | MB_3253 | MB_7291 | MB_7243 | MB_3466 | MB_7286 | MB_7292 |
| MUT | MUT | WT | WT | MUT | MUT | WT | WT | WT | WT |
| NULL | MISS | WT | WT | MISS | MISS | WT | WT | WT | WT |
| R213* | L111P | | | V272M | R273H | | | | |
| ERnHER2n | ERnHER2n | ERpHER2n | ERpHER2n | ERpHER2n | HER2p | ERpHER2n | ERpHER2n | ERpHER2n | ERpHER2n |
| Died of Disease | Living | Died of Disease | Died of Disease | Died of Disease | Died of Disease | Died of Other Causes | Died of Other Causes | Died of Other Causes | Died of Disease |
| 14.7 | 90.56666667 | 125.3333333 | 126.6666667 | 55.83333333 | 6.833333333 | 148.8 | 234.1333333 | 158.6333333 | 78.46666667 |
| YES | YES | NO | NO | NO | NO | NO | NO | NO | NO |

| | | | | | | | | | | | |
|---|---|---|---|---|---|---|---|---|---|---|---|
| MB_7275 | MB_7249 | MB_7295 | MB_7297 | MB_7294 | MB_7296 | MB_3402 | MB_5113 | MB_5450 | MB_7032 | MB_7037 | MB_7040 |
| MUT | WT | WT | WT | MUT | MUT | WT | WT | MUT | WT | WT | WT |
| MISS | WT | WT | WT | NULL | MISS | WT | WT | MISS | WT | WT | WT |
| E171G | | | | X331_splice | M237I | | | C141Y | | | |
| HER2p | ERpHER2n | ERpHER2n | ERpHER2n | ERpHER2n | HER2p | ERpHER2n | ERpHER2n | ERnHER2n | ERpHER2n | ERpHER2n | ERpHER2n |
| Died of Disease | Died of Other Causes | Living | Died of Disease | Died of Disease | Died of Disease | Died of Disease | Living | Living | Living | Living | Living |
| 44.4 | 58.66666667 | 196.8666667 | 175.9666667 | 82.73333333 | 44.73333333 | 50.03333333 | 175.6333333 | 190.2 | 71.46666667 | 86.4 | 71.83333333 |
| YES | NO | NO | NO | NO | NO | YES | NO | NO | NO | NO | NO |

| | | | | | | | | | | |
|---|---|---|---|---|---|---|---|---|---|---|
| MB_7020 | MB_7042 | MB_7043 | MB_7028 | MB_7029 | MB_7027 | MB_7060 | MB_7034 | MB_5191 | MB_5214 | MB_5204 |
| MUT | WT | WT | WT | MUT | MUT | WT | WT | WT | WT | WT |
| NULL | WT | WT | WT | NULL | MISS | WT | WT | WT | WT | WT |
| E349_G356del | | | | Q331* | R342P | | | | | |
| HER2p | ERpHER2n | ERpHER2n | ERpHER2n | ERpHER2n | HER2p | ERpHER2n | ERpHER2n | ERpHER2n | ERpHER2n | ERpHER2n |
| Died of Disease | Living | Living | Living | Living | Died of Disease | Died of Other Causes | Living | Died of Other Causes | Died of Other Causes | Died of Other Causes |
| 71.63333333 | 88.5 | 91.53333333 | 89.8 | 76.86666667 | 36.96666667 | 112.6666667 | 75.1 | 142.1666667 | 107.3666667 | 191.9333333 |
| YES | NO | NO | NO | NO | YES | NO | NO | NO | NO | NO |

| | | | | | | | | | |
|---|---|---|---|---|---|---|---|---|---|
| MB_5155 | MB_5190 | MB_5614 | MB_5592 | MB_0437 | MB_0028 | MB_0309 | MB_0100 | MB_5648 | MB_5572 |
| WT | MUT | MUT | WT | WT | MUT | WT | MUT | WT | WT |
| WT | MISS | NULL | WT | WT | MISS | WT | MISS | WT | WT |
| | R273C | X187_splice | | | C242R | | G245S | | |
| ERnHER2n | HER2p | ERpHER2n | ERpHER2n | ERpHER2n | ERpHER2n | ERpHER2n | ERnHER2n | ERpHER2n | ERnHER2n |
| Living | Living | Died of Disease | Died of Other Causes | Living | Died of Other Causes | Died of Other Causes | Died of Disease | Died of Other Causes | Living |
| 259.9333333 | 242.5666667 | 116.4333333 | 26.33333333 | 90.56666667 | 36.56666667 | 42.33333333 | 8.066666667 | 199.9666667 | 178.6333333 |
| YES | NO | NO | NO | NO | NO | NO | YES | NO | NO |

| | | | | | | | | | |
|---|---|---|---|---|---|---|---|---|---|
| MB_5655 | MB_5577 | MB_5585 | MB_5620 | MB_5652 | MB_0537 | MB_0544 | MB_0468 | MB_0333 | MB_0205 |
| MUT | MUT | WT | WT | MUT | WT | WT | WT | MUT | WT |
| NULL | MISS | WT | WT | MISS | WT | WT | WT | MISS | WT |
| I255del | V216M | | | P190T | | | | p.R273C | |
| ERnHER2n | ERnHER2n | ERpHER2n | ERpHER2n | ERpHER2n | ERpHER2n | ERpHER2n | ERpHER2n | ERnHER2n | ERpHER2n |
| Living | Died of Disease | Died of Disease | Died of Disease | Died of Other Causes | Living | Died of Other Causes | Died of Other Causes | Died of Disease | Living |
| 191.8 | 55.4 | 53.9 | 42.06666667 | 104.6666667 | 113.5666667 | 79.33333333 | 129.4333333 | 26.56666667 | 144.4666667 |
| NO | NO | NO | NO | NO | YES | YES | YES | YES | NO |

| | | | | | | | | | | |
|---|---|---|---|---|---|---|---|---|---|---|
| MB_0233 | MB_0210 | MB_0200 | MB_0244 | MB_5588 | MB_5628 | MB_5633 | MB_0169 | MB_0253 | MB_0226 | MB_0278 |
| MUT | WT | MUT | WT | MUT | MUT | WT | WT | WT | WT | MUT |
| MISS | WT | NULL | WT | MISS | NULL | WT | WT | WT | WT | MISS |
| R280S | | R196* | | R248W | P153Afs*28 | | | | | R248Q |
| ERpHER2n | ERnHER2n | ERnHER2n | ERpHER2n | ERpHER2n | HER2p | ERnHER2n | ERpHER2n | ERpHER2n | ERpHER2n | ERnHER2n |
| Died of Other Causes | Living | Died of Disease | Living | Died of Disease | Died of Disease | Died of Disease | Died of Other Causes | Living | Living | Died of Disease |
| 72.43333333 | 144.9333333 | 128.7 | 149.7333333 | 81.13333333 | 42.6 | 35.03333333 | 145.5 | 152.3 | 57.63333333 | 5.833333333 |
| NO | NO | NO | NO | NO | NO | YES | NO | NO | NO | YES |

| | | | | | | | | | | | |
|---|---|---|---|---|---|---|---|---|---|---|---|
| MB_0266 | MB_0577 | MB_0531 | MB_0245 | MB_5641 | MB_5640 | MB_0534 | MB_0230 | MB_0109 | MB_0304 | MB_0369 | MB_0627 |
| WT | WT | WT | WT | WT | WT | WT | MUT | MUT | MUT | WT | MUT |
| WT | WT | WT | WT | WT | WT | WT | NULL | MISS | MISS | WT | MISS |
| | | | | | | | I255del | C277F | H193L | | R280T |
| ERpHER2n | ERpHER2n | ERpHER2n | ERpHER2n | ERpHER2n | ERpHER2n | ERpHER2n | HER2p | ERpHER2n | ERpHER2n | ERpHER2n | ERnHER2n |
| Died of Other Causes | Living | Living | Living | Living | Living | Living | Living | Died of Other Causes | Died of Disease | Living | Living |
| 90.66666667 | 65.4 | 163.8666667 | 164.7 | 181.8666667 | 172.3 | 124.1 | 200.3333333 | 112.4 | 111.5333333 | 144.3333333 | 0.766666667 |
| NO | NO | YES | NO | NO | NO | YES | NO | NO | NO | YES | NO |

| | | | | | | | | | | |
|---|---|---|---|---|---|---|---|---|---|---|
| MB_0492 | MB_0147 | MB_0442 | MB_0111 | MB_0634 | MB_0280 | MB_0490 | MB_0178 | MB_0242 | MB_5624 | MB_5593 |
| WT | WT | WT | WT | WT | WT | WT | MUT | WT | MUT | MUT |
| WT | WT | WT | WT | WT | WT | WT | NULL | WT | NULL | NULL |
| | | | | | | | X225_splice | | L45Cfs*78 | -125fs |
| ERpHER2n | ERpHER2n | ERpHER2n | ERpHER2n | ERnHER2n | ERpHER2n | ERpHER2n | ERpHER2n | ERpHER2n | ERnHER2n | HER2p |
| Died of Other Causes | Died of Disease | Died of Disease | Living | Living | Died of Disease | Living | Died of Disease | Living | Living | Died of Disease |
| 81.13333333 | 51.76666667 | 103.4666667 | 127.1 | 17.76666667 | 172.9 | 101.9666667 | 104.4666667 | 199.1 | 136 | 92.5 |
| NO | NO | NO | NO | NO | NO | NO | NO | NO | YES | YES |

| | | | | | | | | | | |
|---|---|---|---|---|---|---|---|---|---|---|
| MB_4010 | MB_5605 | MB_5636 | MB_4529 | MB_5566 | MB_0268 | MB_0275 | MB_0127 | MB_0045 | MB_0414 | MB_0259 |
| WT | WT | WT | WT | MUT | MUT | WT | MUT | WT | MUT | MUT |
| WT | WT | WT | WT | NULL | NULL | WT | NULL | WT | NULL | MISS |
| | | | | R282Pfs*62 | R306* | | Q317Sfs*28 | | E171Gfs*3 | R273H |
| ERpHER2n | ERpHER2n | ERpHER2n | ERpHER2n | ERnHER2n | ERpHER2n | ERpHER2n | ERnHER2n | ERnHER2n | ERnHER2n | ERnHER2n |
| Died of Other Causes | Living | Died of Other Causes | Died of Other Causes | Living | Died of Disease | Living | Living | Living | Living | Died of Disease |
| 11.7 | 123.7 | 108.3 | 126.4666667 | 234.4 | 28.5 | 186.1 | 132.0666667 | 164.9 | 76.63333333 | 11.86666667 |
| NO | NO | NO | NO | YES | NO | NO | YES | YES | NO | YES |

| | | | | | | | | | | |
|---|---|---|---|---|---|---|---|---|---|---|
| MB_0396 | MB_0874 | MB_0179 | MB_0526 | MB_0586 | MB_0508 | MB_0288 | MB_0269 | MB_0582 | MB_0901 | MB_0000 |
| MUT | MUT | WT | WT | WT | WT | WT | WT | MUT | MUT | WT |
| MISS | MISS | WT | WT | WT | WT | WT | WT | MISS | NULL | WT |
| H179R | R280T | | | | | | | P151H | N131Cfs*27 | |
| ERnHER2n | ERnHER2n | ERnHER2n | ERpHER2n | ERpHER2n | ERpHER2n | HER2p | ERnHER2n | ERnHER2n | ERnHER2n | ERpHER2n |
| Living | Died of Disease | Died of Disease | Died of Other Causes | Died of Other Causes | Living | Died of Disease | Living | Died of Other Causes | Living | Living |
| 60.66666667 | 16.6 | 17.93333333 | 139.6333333 | 77.23333333 | 114.3333333 | 63.8 | 22.23333333 | 15.53333333 | 136.1666667 | 140.5 |
| YES | YES | YES | NO | NO | NO | YES | YES | NO | YES | NO |

| | | | | | | | | | | |
|---|---|---|---|---|---|---|---|---|---|---|
| MB_0535 | MB_0469 | MB_0478 | MB_0499 | MB_0170 | MB_0464 | MB_0421 | MB_0623 | MB_0241 | MB_0163 | MB_0149 |
| WT | MUT | WT | WT | WT | MUT | MUT | WT | MUT | MUT | MUT |
| WT | MISS | WT | WT | WT | MISS | NULL | WT | MISS | NULL | MISS |
| | R175H | | | | E285K | L194_I195insSIL | | A159V | S106Rfs*41 | R175H |
| ERpHER2n | ERpHER2n | ERpHER2n | ERnHER2n | ERpHER2n | ERnHER2n | HER2p | ERpHER2n | ERpHER2n | ERnHER2n | ERnHER2n |
| Living | Living | Living | Living | Died of Other Causes | Died of Disease | Died of Other Causes | Living | Died of Disease | Died of Other Causes | Died of Other Causes |
| 199.1333333 | 131.2666667 | 132.3 | 131.9 | 93.36666667 | 39.3 | 92.86666667 | 111.0666667 | 73.13333333 | 98.1 | 51.7 |
| NO | YES | NO | NO | NO | YES | NO | NO | NO | NO | NO |

| | | | | | | | | | |
|---|---|---|---|---|---|---|---|---|---|
| MB_0350 | MB_0512 | MB_0020 | MB_0046 | MB_0134 | MB_0495 | MB_0470 | MB_0872 | MB_0661 | MB_0339 |
| MUT | WT | WT | MUT | WT | MUT | WT | WT | WT | WT |
| MISS | WT | WT | MISS | WT | MISS | WT | WT | WT | WT |
| R248G | | | C135R | | G245S | | | | |
| ERnHER2n | ERpHER2n | ERnHER2n | HER2p | ERpHER2n | ERnHER2n | ERnHER2n | ERpHER2n | ERpHER2n | ERpHER2n |
| Died of Disease | Living | Died of Disease | Died of Other Causes | Died of Disease | Died of Disease | Living | Died of Other Causes | Died of Disease | Died of Other Causes |
| 46.06666667 | 121.5333333 | 22.4 | 14.13333333 | 12.93333333 | 71.8 | 88.23333333 | 152.0666667 | 20 | 26.73333333 |
| NO | NO | YES | NO | NO | YES | NO | NO | NO | NO |

| | | | | | | | | | | | |
|---|---|---|---|---|---|---|---|---|---|---|---|
| MB_0264 | MB_0175 | MB_0303 | MB_0172 | MB_0125 | MB_0340 | MB_0234 | MB_0487 | MB_0305 | MB_0502 | MB_0663 | MB_0130 |
| WT | WT | WT | MUT | WT | MUT | WT | WT | WT | WT | MUT | WT |
| WT | WT | WT | MISS | WT | NULL | WT | WT | WT | WT | NULL | WT |
| | | | R110P | | X261_splice | | | | | C238* | |
| ERpHER2n | ERpHER2n | ERnHER2n | ERpHER2n | ERpHER2n | ERnHER2n | ERpHER2n | ERpHER2n | ERpHER2n | ERnHER2n | HER2p | HER2p |
| Died of Disease | Living | Living | Living | Living | Living | Died of Other Causes | Living | Died of Disease | Living | Living | Living |
| 43.1 | 72.36666667 | 60.13333333 | 138.1 | 1.266666667 | 164.7333333 | 94.23333333 | 86.9 | 63.5 | 82.1 | 55.36666667 | 153.5666667 |
| NO | NO | NO | YES | NO | NO | NO | NO | YES | YES | YES | NO |

| | | | | | | | | | | |
|---|---|---|---|---|---|---|---|---|---|---|
| MB_0289 | MB_0129 | MB_0343 | MB_0458 | MB_0570 | MB_0488 | MB_0655 | MB_0235 | MB_0484 | MB_0436 | MB_0877 |
| WT | WT | WT | WT | MUT | WT | WT | WT | MUT | MUT | MUT |
| WT | WT | WT | WT | NULL | WT | WT | WT | MISS | MISS | MISS |
| | | | | Q192* | | | | R248G | R273H | R248Q |
| ERnHER2n | HER2p | ERpHER2n | ERpHER2n | ERpHER2n | ERpHER2n | ERpHER2n | ERpHER2n | HER2p | ERnHER2n | ERpHER2n |
| Died of Other Causes | Died of Other Causes | Died of Disease | Living | Living | Living | Living | Died of Disease | Living | Died of Other Causes | Died of Disease |
| 71.6 | 38.56666667 | 91.13333333 | 134.5 | 272.2 | 112.9333333 | 111.0666667 | 142.5666667 | 82.63333333 | 75.5 | 14.43333333 |
| NO | YES | NO | YES | NO | NO | NO | YES | YES | NO | NO |

| | | | | | | | | | | |
|---|---|---|---|---|---|---|---|---|---|---|
| MB_0097 | MB_0897 | MB_0095 | MB_0227 | MB_0353 | MB_0522 | MB_0543 | MB_0646 | MB_0293 | MB_0552 | MB_0260 |
| WT | WT | WT | WT | MUT | MUT | WT | WT | WT | WT | WT |
| WT | WT | WT | WT | MISS | MISS | WT | WT | WT | WT | WT |
| | | | | Q136H | R248W | | | | | |
| ERpHER2n | ERpHER2n | ERpHER2n | ERpHER2n | ERpHER2n | HER2p | ERpHER2n | ERpHER2n | ERpHER2n | HER2p | ERpHER2n |
| Living | Died of Other Causes | Died of Other Causes | Living | Died of Other Causes | Living | Living | Died of Disease | Living | Died of Disease | Died of Other Causes |
| 98.7 | 30.96666667 | 49.76666667 | 111.8666667 | 41.83333333 | 2.5 | 154.7666667 | 15.16666667 | 64.36666667 | 68.06666667 | 169.8333333 |
| NO | NO | NO | NO | NO | NO | NO | NO | NO | YES | NO |

| | | | | | | | | | | | | |
|---|---|---|---|---|---|---|---|---|---|---|---|---|
| MB_0256 | MB_0626 | MB_0211 | MB_0124 | MB_0510 | MB_0365 | MB_0290 | MB_0113 | MB_0620 | MB_0282 | MB_0228 | MB_0578 | MB_0479 |
| WT | MUT | MUT | WT | WT | MUT | WT | WT | WT | WT | WT | WT | MUT |
| WT | MISS | MISS | WT | WT | NULL | WT | WT | WT | WT | WT | WT | NULL |
| | F134L | R248Q | | | X332_splice | | | | | | | Q136* |
| ERpHER2n | ERpHER2n | ERnHER2n | ERpHER2n | ERpHER2n | HER2p | ERpHER2n | HER2p | ERpHER2n | ERpHER2n | ERpHER2n | ERpHER2n | HER2p |
| Living | Died of Disease | Died of Disease | Living | Living | Died of Disease | Living | Living | Living | Living | Died of Disease | Living | Living |
| 200.7 | 35.53333333 | 44.8 | 118.2 | 128.4 | 87.23333333 | 199.5333333 | 43.16666667 | 112.8 | 194.2333333 | 10.83333333 | 110.1 | 132.7666667 |
| NO | NO | NO | YES | NO | NO | NO | YES | YES | YES | NO | YES | YES |

| | | | | | | | | | | |
|---|---|---|---|---|---|---|---|---|---|---|
| MB_0263 | MB_0509 | MB_0279 | MB_0168 | MB_0588 | MB_0462 | MB_0262 | MB_0554 | MB_0418 | MB_0193 | MB_0652 |
| WT | MUT | WT | WT | WT | MUT | MUT | WT | WT | WT | MUT |
| WT | MISS | WT | WT | WT | MISS | NULL | WT | WT | WT | MISS |
| | R248G | | | | I195T | N131del | | | | R248Q |
| ERpHER2n | ERpHER2n | ERpHER2n | ERpHER2n | ERnHER2n | HER2p | ERpHER2n | ERpHER2n | ERpHER2n | ERpHER2n | HER2p |
| Died of Other Causes | Living | Died of Other Causes | Living | Living | Living | Died of Other Causes | Died of Other Causes | Living | Living | Died of Disease |
| 176.0333333 | 115.3 | 168.3333333 | 122.7666667 | 119.7333333 | 132.7666667 | 145.3 | 111.1666667 | 102.0666667 | 19 | 96.23333333 |
| NO | NO | NO | NO | NO | NO | NO | NO | NO | NO | NO |

| | | | | | | | | | | |
|---|---|---|---|---|---|---|---|---|---|---|
| MB_0638 | MB_0188 | MB_0617 | MB_7225 | MB_7141 | MB_7234 | MB_7089 | MB_7030 | MB_7004 | MB_7119 | MB_7263 |
| WT | WT | WT | MUT | WT | WT | MUT | WT | WT | MUT | MUT |
| WT | WT | WT | MISS | WT | WT | MISS | WT | WT | NULL | MISS |
| | | | P278R | | | P151A | | | R110Pfs*39 | D352Y |
| ERpHER2n | ERnHER2n | ERnHER2n | ERnHER2n | ERpHER2n | ERpHER2n | ERnHER2n | ERnHER2n | ERpHER2n | ERnHER2n | ERpHER2n |
| Died of Other Causes | Died of Disease | Died of Other Causes | Died of Disease | Living | Living | Living | Died of Other Causes | Living | Living | Died of Other Causes |
| 103.6333333 | 31.3 | 92.83333333 | 55 | 138.5666667 | 222.2 | 118.9 | 31.33333333 | 50.46666667 | 128.3666667 | 164.9666667 |
| NO | YES | NO | YES | NO | NO | YES | NO | NO | YES | NO |

| | | | | | | | | | | |
|---|---|---|---|---|---|---|---|---|---|---|
| MB_0174 | MB_7182 | MB_7039 | MB_3797 | MB_7112 | MB_0308 | MB_7230 | MB_5452 | MB_6195 | MB_6317 | MB_5460 |
| MUT | WT | MUT | WT | WT | WT | WT | WT | WT | MUT | WT |
| NULL | WT | MISS | WT | WT | WT | WT | WT | WT | MISS | WT |
| G154Afs*16 | | R342P | | | | | | | Y205N | |
| ERnHER2n | ERpHER2n | ERnHER2n | ERpHER2n | ERpHER2n | ERpHER2n | ERpHER2n | ERpHER2n | ERpHER2n | ERpHER2n | ERpHER2n |
| Living | Died of Disease | Died of Disease | Living | Died of Disease | Living | Living | Died of Other Causes | Living | Died of Other Causes | Living |
| 78.76666667 | 49.46666667 | 49.2 | 228.3333333 | 64.7 | 183.2666667 | 182.6 | 16.16666667 | 202.2333333 | 29.23333333 | 218.6333333 |
| YES | YES | NO | NO | YES | NO | NO | NO | NO | NO | NO |

| | | | | | | | | | | | |
|---|---|---|---|---|---|---|---|---|---|---|---|
| MB_5464 | MB_5552 | MB_5547 | MB_6189 | MB_6122 | MB_6192 | MB_4820 | MB_5527 | MB_5167 | MB_5465 | MB_5453 | MB_5471 |
| WT | WT | MUT | WT | MUT | MUT | WT | WT | WT | MUT | WT | MUT |
| WT | WT | MISS | WT | NULL | NULL | WT | WT | WT | NULL | WT | MISS |
| | | I255T | | R213* | E287Ifs*63 | | | | R213* | | R248Q |
| ERpHER2n | ERpHER2n | ERnHER2n | ERpHER2n | ERpHER2n | ERpHER2n | ERpHER2n | HER2p | ERpHER2n | ERnHER2n | ERnHER2n | ERpHER2n |
| Died of Other Causes | Died of Disease | Living | Living | Living | Died of Other Causes | Died of Disease | Living | Living | Died of Disease | Died of Disease | Living |
| 116.5333333 | 34.7 | 98.56666667 | 240.2 | 260.2 | 42.63333333 | 57.3 | 180.5666667 | 208.4 | 18.8 | 57.3 | 185.7666667 |
| NO | NO | YES | NO | YES | NO | NO | YES | NO | NO | YES | NO |

| | | |
|---|---|---|
| MB_5127 | MB_4313 | MB_4823 |
| WT | WT | MUT |
| WT | WT | NULL |
| | | p.? |
| ERpHER2n | ERpHER2n | HER2p |
| Died of Other Causes | Died of Disease | Died of Other Causes |
| 191.4666667 | 300.7 | 282.3 |
| NO | NO | NO |
