## Supplemental File 4 for "Transcriptional profiling reveals a subset of human breast tumors that retain wt *TP53* but display mutant p53-associated features"

### Mutations in other cancer-related genes identified by sequencing in the PM tumors

| METABRIC.ID | gene | vaf | reads | TP53.LOH |
|---|---|---|---|---|
| MB-0054 | PIK3CA | 0.27 | 37 | NO |
| MB-0054 | ERBB3 | 0.31 | 71 | NO |
| MB-0054 | NCOR2 | 0.55 | 20 | NO |
| MB-0054 | MTAP | 0.5 | 58 | NO |
| MB-0066 | PTEN | 0.225 | 80 | NO |
| MB-0066 | MAP2K4 | 0.196 | 56 | NO |
| MB-0066 | PIK3CA | 0.149 | 94 | NO |
| MB-0066 | CASP8 | 0.109 | 46 | NO |
| MB-0131 | PIK3CA | 0.333 | 105 | YES |
| MB-0131 | KMT2C | 0.349 | 189 | YES |
| MB-0131 | NRAS | 0.271 | 107 | YES |
| MB-0131 | KMT2C | 0.313 | 281 | YES |
| MB-0131 | GATA3 | 0.373 | 161 | YES |
| MB-0131 | CTCF | 0.591 | 181 | YES |
| MB-0143 | KRAS | 0.383 | 162 | NO |
| MB-0143 | SIK2 | 0.328 | 61 | NO |
| MB-0143 | RYR2 | 0.375 | 256 | NO |
| MB-0143 | ASXL1 | 0.38 | 213 | NO |
| MB-0143 | CDH1 | 0.608 | 166 | NO |
| MB-0143 | NCOR2 | 0.375 | 40 | NO |
| MB-0143 | TBX3 | 0.383 | 133 | NO |
| MB-0143 | PIK3CA | 0.336 | 211 | NO |
| MB-0143 | MUC16 | 0.167 | 186 | NO |
| MB-0143 | TBX3 | 0.362 | 130 | NO |
| MB-0147 | CLK3 | 0.631 | 65 | YES |
| MB-0147 | PIK3R1 | 0.31 | 129 | YES |
| MB-0147 | EP300 | 0.25 | 20 | YES |
| MB-0147 | CDH1 | 0.667 | 24 | YES |
| MB-0173 | FOXO3 | 0.386 | 140 | YES |
| MB-0173 | DNAH2 | 0.383 | 227 | YES |
| MB-0173 | UTRN | 0.471 | 257 | YES |
| MB-0173 | RYR2 | 0.408 | 157 | YES |
| MB-0195 | PRPS2 | 0.353 | 85 | NO |
| MB-0195 | PALLD | 0.407 | 81 | NO |
| MB-0195 | TBX3 | 0.398 | 161 | NO |
| MB-0195 | BAP1 | 0.709 | 127 | NO |
| MB-0195 | SGCD | 0.471 | 206 | NO |
| MB-0197 | SIAH1 | 0.19 | 100 | NO |
| MB-0197 | SHANK2 | 0.225 | 173 | NO |
| MB-0197 | STMN2 | 0.128 | 430 | NO |
| MB-0197 | NCOA3 | 0.143 | 28 | NO |
| MB-0197 | ROS1 | 0.169 | 71 | NO |
| MB-0197 | PTPRM | 0.151 | 146 | NO |
| MB-0197 | ROS1 | 0.169 | 71 | NO |
| MB-0311 | MALAT1 | 0.493 | 71 | NO |
| MB-0311 | COL6A3 | 0.543 | 138 | NO |
| MB-0311 | AHNAK | 0.611 | 18 | NO |
| MB-0311 | MUC16 | 0.206 | 155 | NO |
| MB-0311 | AFF2 | 0.122 | 164 | NO |
| MB-0311 | UBR5 | 0.543 | 105 | NO |
| MB-0311 | MUC16 | 0.194 | 170 | NO |

| | | | | |
|---|---|---|---|---|
| MB-0311 | NCOR1 | 0.647 | 17 | NO |
| MB-0311 | MALAT1 | 0.22 | 100 | NO |
| MB-0311 | MAP2K4 | 0.297 | 74 | NO |
| MB-0311 | JAK1 | 0.469 | 81 | NO |
| MB-0311 | PIK3CA | 0.451 | 164 | NO |
| MB-0311 | RYR2 | 0.149 | 208 | NO |
| MB-0311 | MALAT1 | 0.516 | 126 | NO |
| MB-0311 | AFF2 | 0.214 | 192 | NO |
| MB-0311 | AGMO | 0.462 | 80 | NO |
| MB-0311 | SMARCC1 | 0.263 | 19 | NO |
| MB-0311 | STMN2 | 0.122 | 98 | NO |
| MB-0311 | DNAH5 | 0.395 | 43 | NO |
| MB-0311 | ARID2 | 0.158 | 38 | NO |
| MB-0393 | PIK3CA | 0.327 | 104 | YES |
| MB-0393 | SF3B1 | 0.053 | 75 | YES |
| MB-0393 | PIK3CA | 0.254 | 185 | YES |
| MB-0532 | MUC16 | 0.118 | 718 | YES |
| MB-0532 | PIK3CA | 0.254 | 232 | YES |
| MB-0532 | CHD1 | 0.137 | 168 | YES |
| MB-0532 | DNAH5 | 0.5 | 72 | YES |
| MB-0532 | CDKN1B | 0.21 | 157 | YES |
| MB-0532 | TP53 | 0.155 | 155 | YES |
| MB-0532 | TBX3 | 0.108 | 287 | YES |
| MB-0532 | BRCA1 | 0.488 | 254 | YES |
| MB-0532 | MLL2 | 0.462 | 675 | YES |
| MB-0532 | AHNAK2 | 0.409 | 820 | YES |
| MB-0532 | CBFB | 0.267 | 30 | YES |
| MB-0532 | PALLD | 0.524 | 82 | YES |
| MB-0532 | MLL2 | 0.447 | 622 | YES |
| MB-0532 | PPP2R2A | 0.454 | 163 | YES |
| MB-0532 | TP53 | 0.143 | 140 | YES |
| MB-0532 | TP53 | 0.145 | 145 | YES |
| MB-0532 | THSD7A | 0.515 | 270 | YES |
| MB-0542 | MEN1 | 0.433 | 127 | NO |
| MB-0542 | MYO3A | 0.439 | 262 | NO |
| MB-0542 | HERC2 | 0.497 | 191 | NO |
| MB-0542 | TG | 0.505 | 182 | NO |
| MB-0542 | SMARCC2 | 0.425 | 200 | NO |
| MB-0542 | ERBB3 | 0.486 | 140 | NO |
| MB-0542 | TG | 0.471 | 278 | NO |
| MB-0542 | RYR2 | 0.491 | 169 | NO |
| MB-0542 | PRKG1 | 0.56 | 75 | NO |
| MB-0542 | NRG3 | 0.424 | 184 | NO |
| MB-0542 | NPNT | 0.409 | 186 | NO |
| MB-0542 | NPNT | 0.497 | 197 | NO |
| MB-0542 | CDH1 | 0.412 | 51 | NO |
| MB-0542 | PIK3CA | 0.467 | 75 | NO |
| MB-0542 | DTWD2 | 0.5 | 88 | NO |
| MB-0543 | PDE4DIP | 0.2 | 40 | NO |
| MB-0666 | PIK3CA | 0.189 | 37 | NO |
| MB-2634 | SBNO1 | 0.392 | 194 | NO |
| MB-2634 | ARID1A | 0.439 | 41 | NO |

| | | | | |
|---|---|---|---|---|
| MB-2634 | PRR16 | 0.294 | 201 | NO |
| MB-2634 | SYNE1 | 0.613 | 31 | NO |
| MB-2634 | TP53 | 0.789 | 109 | NO |
| MB-2634 | LIFR | 0.417 | 151 | NO |
| MB-2634 | SBNO1 | 0.421 | 57 | NO |
| MB-2634 | MLL2 | 0.566 | 76 | NO |
| MB-2634 | PTPRD | 0.281 | 32 | NO |
| MB-2634 | SYNE1 | 0.539 | 167 | NO |
| MB-3389 | LAMA2 | 0.175 | 80 | NO |
| MB-3389 | CHD1 | 0.404 | 94 | NO |
| MB-3389 | ARID2 | 0.379 | 124 | NO |
| MB-3389 | MYO1A | 0.253 | 75 | NO |
| MB-3389 | GATA3 | 0.579 | 114 | NO |
| MB-3389 | FAM20C | 0.851 | 134 | NO |
| MB-3389 | USP28 | 0.2 | 35 | NO |
| MB-3466 | MLLT4 | 0.108 | 185 | YES |
| MB-3466 | LAMA2 | 0.123 | 195 | YES |
| MB-3466 | AKAP9 | 0.276 | 199 | YES |
| MB-3466 | AFF2 | 0.18 | 100 | YES |
| MB-3466 | DNAH2 | 0.363 | 201 | YES |
| MB-3466 | MALAT1 | 0.242 | 360 | YES |
| MB-3466 | ZFP36L1 | 0.151 | 192 | YES |
| MB-3466 | ERBB3 | 0.254 | 71 | YES |
| MB-3466 | NOTCH1 | 0.146 | 192 | YES |
| MB-3466 | PTPRD | 0.329 | 146 | YES |
| MB-3466 | KDM3A | 0.312 | 77 | YES |
| MB-3466 | MUC16 | 0.11 | 463 | YES |
| MB-3466 | SIAH1 | 0.254 | 59 | YES |
| MB-3466 | NRG3 | 0.306 | 235 | YES |
| MB-3466 | SMAD2 | 0.173 | 208 | YES |
| MB-3466 | MALAT1 | 0.312 | 157 | YES |
| MB-3466 | PTPN22 | 0.185 | 92 | YES |
| MB-3466 | NPNT | 0.181 | 293 | YES |
| MB-3466 | TG | 0.277 | 173 | YES |
| MB-3466 | LAMB3 | 0.255 | 506 | YES |
| MB-3466 | KDM3A | 0.135 | 170 | YES |
| MB-3466 | TBL1XR1 | 0.112 | 251 | YES |
| MB-3466 | HIST1H2BC | 0.244 | 176 | YES |
| MB-3600 | MUC16 | 0.126 | 309 | NO |
| MB-3600 | PTPRD | 0.376 | 165 | NO |
| MB-3600 | USP9X | 0.444 | 169 | NO |
| MB-3600 | CTNNA1 | 0.667 | 21 | NO |
| MB-3600 | BIRC6 | 0.243 | 37 | NO |
| MB-3600 | SETD1A | 0.488 | 82 | NO |
| MB-3600 | ATR | 0.5 | 44 | NO |
| MB-3600 | FOXP1 | 0.529 | 70 | NO |
| MB-3600 | GATA3 | 0.285 | 242 | NO |
| MB-3600 | ROS1 | 0.365 | 104 | NO |
| MB-3600 | MAP3K1 | 0.421 | 164 | NO |
| MB-3600 | RUNX1 | 0.438 | 208 | NO |
| MB-3600 | FANCD2 | 0.187 | 182 | NO |
| MB-3600 | AHNAK | 0.207 | 300 | NO |

| | | | | |
|---|---|---|---|---|
| MB-3600 | KMT2C | 0.208 | 72 | NO |
| MB-3600 | TG | 0.185 | 634 | NO |
| MB-3600 | LAMA2 | 0.257 | 183 | NO |
| MB-4148 | NCOR2 | 0.614 | 153 | NO |
| MB-4148 | NCOR2 | 0.445 | 110 | NO |
| MB-4148 | PTPRD | 0.64 | 150 | NO |
| MB-4148 | SYNE1 | 0.556 | 81 | NO |
| MB-4148 | ALK | 0.261 | 218 | NO |
| MB-4148 | HERC2 | 0.56 | 134 | NO |
| MB-4148 | FOXP1 | 0.432 | 162 | NO |
| MB-4148 | CHD1 | 0.482 | 114 | NO |
| MB-4148 | AKT2 | 0.64 | 175 | NO |
| MB-4148 | SHANK2 | 0.467 | 105 | NO |
| MB-4148 | EP300 | 0.475 | 118 | NO |
| MB-4148 | BIRC6 | 0.667 | 18 | NO |
| MB-4148 | TG | 0.331 | 353 | NO |
| MB-4148 | SYNE1 | 0.533 | 272 | NO |
| MB-4148 | SETDB1 | 0.318 | 274 | NO |
| MB-4148 | GPR124 | 0.194 | 31 | NO |
| MB-4148 | AHNAK | 0.482 | 328 | NO |
| MB-4148 | NF1 | 0.443 | 122 | NO |
| MB-4235 | SMAD4 | 0.458 | 48 | YES |
| MB-4235 | MYH9 | 0.196 | 46 | YES |
| MB-4235 | SYNE1 | 0.437 | 87 | YES |
| MB-4235 | PDE4DIP | 0.213 | 253 | YES |
| MB-4235 | PIK3CA | 0.076 | 158 | YES |
| MB-4235 | MUC16 | 0.43 | 179 | YES |
| MB-4235 | CBFB | 0.154 | 26 | YES |
| MB-4235 | MAP3K1 | 0.108 | 166 | YES |
| MB-4278 | UBR5 | 0.115 | 87 | YES |
| MB-4278 | KMT2C | 0.126 | 111 | YES |
| MB-4278 | AHNAK2 | 0.203 | 153 | YES |
| MB-4278 | ARID5B | 0.486 | 148 | YES |
| MB-4278 | PIK3R1 | 0.13 | 46 | YES |
| MB-4278 | UBR5 | 0.481 | 27 | YES |
| MB-4278 | MAP3K1 | 0.122 | 82 | YES |
| MB-4278 | KDM3A | 0.5 | 88 | YES |
| MB-4278 | KDM6A | 0.127 | 71 | YES |
| MB-4278 | LAMB3 | 0.101 | 159 | YES |
| MB-4278 | GATA3 | 0.083 | 121 | YES |
| MB-4278 | APC | 0.523 | 128 | YES |
| MB-4278 | NCOR1 | 0.159 | 82 | YES |
| MB-4278 | COL22A1 | 0.154 | 78 | YES |
| MB-4278 | AGMO | 0.104 | 67 | YES |
| MB-4278 | PIK3CA | 0.05 | 181 | YES |
| MB-4278 | CACNA2D3 | 0.448 | 125 | YES |
| MB-4278 | BRCA2 | 0.125 | 104 | YES |
| MB-4278 | RYR2 | 0.139 | 72 | YES |
| MB-4278 | MUC16 | 0.117 | 154 | YES |
| MB-4278 | EP300 | 0.133 | 30 | YES |
| MB-4278 | MUC16 | 0.107 | 214 | YES |
| MB-4278 | SYNE1 | 0.252 | 135 | YES |

| | | | | |
|---|---|---|---|---|
| MB-4278 | MALAT1 | 0.139 | 72 | YES |
| MB-4278 | NF2 | 0.444 | 99 | YES |
| MB-4278 | DNAH2 | 0.138 | 29 | YES |
| MB-4278 | NCOR1 | 0.4 | 150 | YES |
| MB-4278 | MYO1A | 0.434 | 136 | YES |
| MB-4278 | FRMD3 | 0.488 | 125 | YES |
| MB-4278 | HERC2 | 0.143 | 112 | YES |
| MB-4767 | PBRM1 | 0.674 | 46 | NO |
| MB-4767 | DNAH2 | 0.723 | 47 | NO |
| MB-4767 | CHEK2 | 0.111 | 90 | NO |
| MB-4767 | NCOR1 | 0.378 | 90 | NO |
| MB-4767 | CACNA2D3 | 0.292 | 65 | NO |
| MB-4767 | MEN1 | 0.474 | 57 | NO |
| MB-4767 | LAMA2 | 0.568 | 74 | NO |
| MB-4767 | COL6A3 | 0.137 | 102 | NO |
| MB-4767 | SETD2 | 0.308 | 117 | NO |
| MB-4767 | PDE4DIP | 0.318 | 245 | NO |
| MB-4767 | NOTCH1 | 0.5 | 40 | NO |
| MB-4767 | BIRC6 | 0.469 | 98 | NO |
| MB-4834 | KMT2C | 0.415 | 82 | YES |
| MB-4834 | ARID1B | 0.362 | 152 | YES |
| MB-4834 | EP300 | 0.768 | 112 | YES |
| MB-4834 | L1CAM | 0.777 | 282 | YES |
| MB-4834 | LIFR | 0.452 | 93 | YES |
| MB-4834 | SHANK2 | 0.646 | 457 | YES |
| MB-4834 | FANCA | 0.446 | 175 | YES |
| MB-4834 | ERBB4 | 0.505 | 182 | YES |
| MB-4834 | THADA | 0.432 | 370 | YES |
| MB-4834 | ROS1 | 0.3 | 20 | YES |
| MB-4834 | RPGR | 0.59 | 78 | YES |
| MB-4834 | COL22A1 | 0.254 | 71 | YES |
| MB-4834 | SYNE1 | 0.621 | 174 | YES |
| MB-4834 | NOTCH1 | 0.494 | 89 | YES |
| MB-4834 | RPGR | 0.528 | 176 | YES |
| MB-4834 | STAB2 | 0.488 | 291 | YES |
| MB-4834 | NCOA3 | 0.52 | 331 | YES |
| MB-4834 | SBNO1 | 0.5 | 54 | YES |
| MB-4834 | SMARCC1 | 0.445 | 229 | YES |
| MB-4834 | MLLT4 | 0.172 | 87 | YES |
| MB-4834 | SMAD2 | 0.15 | 266 | YES |
| MB-4834 | MUC16 | 0.202 | 382 | YES |
| MB-4834 | MAGEA8 | 0.203 | 182 | YES |
| MB-4834 | GPR124 | 0.474 | 135 | YES |
| MB-4834 | DNAH5 | 0.405 | 311 | YES |
| MB-4834 | ASXL1 | 0.505 | 412 | YES |
| MB-4834 | PIK3R1 | 0.462 | 264 | YES |
| MB-4834 | UBR5 | 0.363 | 91 | YES |
| MB-4834 | LAMB3 | 0.446 | 352 | YES |
| MB-4834 | PDE4DIP | 0.191 | 787 | YES |
| MB-4834 | RYR2 | 0.48 | 325 | YES |
| MB-4834 | MAP3K1 | 0.475 | 158 | YES |
| MB-4834 | PALLD | 0.643 | 14 | YES |

| | | | | |
|---|---|---|---|---|
| MB-4834 | RYR2 | 0.169 | 267 | YES |
| MB-4834 | TAF1 | 0.218 | 78 | YES |
| MB-4834 | COL6A3 | 0.536 | 84 | YES |
| MB-4876 | FOXO3 | 0.423 | 26 | YES |
| MB-4876 | PBRM1 | 0.529 | 68 | YES |
| MB-4876 | HERC2 | 0.528 | 286 | YES |
| MB-4876 | SETD2 | 0.585 | 135 | YES |
| MB-4876 | CACNA2D3 | 0.511 | 139 | YES |
| MB-4876 | ARID1B | 0.524 | 254 | YES |
| MB-4876 | RYR2 | 0.595 | 79 | YES |
| MB-4876 | DNAH5 | 0.484 | 289 | YES |
| MB-4876 | KDM6A | 0.333 | 24 | YES |
| MB-4876 | EP300 | 0.487 | 197 | YES |
| MB-4876 | SETD2 | 0.5 | 146 | YES |
| MB-4937 | DNAH2 | 0.326 | 181 | NO |
| MB-4937 | TAF4B | 0.463 | 339 | NO |
| MB-4937 | USP28 | 0.735 | 68 | NO |
| MB-4937 | ASXL2 | 0.268 | 254 | NO |
| MB-4937 | NDFIP1 | 0.508 | 120 | NO |
| MB-4937 | COL6A3 | 0.114 | 272 | NO |
| MB-4937 | FOXO3 | 0.357 | 56 | NO |
| MB-4937 | CHEK2 | 0.324 | 173 | NO |
| MB-4937 | SMAD2 | 0.783 | 46 | NO |
| MB-4937 | EP300 | 0.661 | 56 | NO |
| MB-4937 | AHNAK | 0.609 | 338 | NO |
| MB-4937 | CDH1 | 0.368 | 182 | NO |
| MB-4937 | AHNAK | 0.609 | 340 | NO |
| MB-4937 | NRAS | 0.133 | 60 | NO |
| MB-4937 | APC | 0.572 | 138 | NO |
| MB-4937 | PTPRD | 0.409 | 127 | NO |
| MB-5001 | SYNE1 | 0.244 | 45 | NO |
| MB-5001 | SETD2 | 0.276 | 145 | NO |
| MB-5001 | GATA3 | 0.263 | 137 | NO |
| MB-5001 | SGCD | 0.327 | 147 | NO |
| MB-5001 | TBL1XR1 | 0.292 | 48 | NO |
| MB-5001 | HRAS | 0.406 | 202 | NO |
| MB-5001 | AFF2 | 0.492 | 189 | NO |
| MB-5001 | COL6A3 | 0.328 | 137 | NO |
| MB-5001 | PTEN | 0.333 | 75 | NO |
| MB-5001 | LIFR | 0.231 | 238 | NO |
| MB-5001 | PPP2R2A | 0.384 | 73 | NO |
| MB-5001 | CLK3 | 0.412 | 68 | NO |
| MB-5001 | ARID5B | 0.338 | 133 | NO |
| MB-5001 | CACNA2D3 | 0.744 | 43 | NO |
| MB-5001 | FAM20C | 0.529 | 104 | NO |
| MB-5001 | DCAF4L2 | 0.19 | 311 | NO |
| MB-5001 | KDM3A | 0.321 | 109 | NO |
| MB-5001 | MUC16 | 0.512 | 217 | NO |
| MB-5001 | ARID1A | 0.472 | 36 | NO |
| MB-5001 | AHNAK | 0.672 | 189 | NO |
| MB-5001 | PALLD | 0.476 | 103 | NO |
| MB-5001 | NRAS | 0.447 | 161 | NO |

| | | | | |
|---|---|---|---|---|
| MB-5001 | HERC2 | 0.217 | 46 | NO |
| MB-5188 | TAF4B | 0.529 | 272 | NO |
| MB-5188 | FLT3 | 0.174 | 23 | NO |
| MB-5188 | TAF1 | 0.259 | 54 | NO |
| MB-5188 | ROS1 | 0.32 | 75 | NO |
| MB-5188 | SIK2 | 0.486 | 173 | NO |
| MB-5188 | SYNE1 | 0.447 | 141 | NO |
| MB-5188 | ARID1A | 0.519 | 241 | NO |
| MB-5188 | MALAT1 | 0.122 | 197 | NO |
| MB-5188 | EP300 | 0.446 | 130 | NO |
| MB-5188 | ROS1 | 0.466 | 73 | NO |
| MB-5188 | LAMA2 | 0.269 | 119 | NO |
| MB-5188 | EP300 | 0.485 | 268 | NO |
| MB-5350 | AHNAK2 | 0.574 | 197 | NO |
| MB-5350 | ALK | 0.661 | 112 | NO |
| MB-5350 | AHNAK2 | 0.596 | 161 | NO |
| MB-5350 | AHNAK2 | 0.545 | 66 | NO |
| MB-5350 | ARID5B | 0.555 | 128 | NO |
| MB-5350 | THSD7A | 0.366 | 41 | NO |
| MB-5350 | PTEN | 0.445 | 119 | NO |
| MB-5350 | MUC16 | 0.471 | 174 | NO |
| MB-5350 | AHNAK2 | 0.639 | 191 | NO |
| MB-5350 | STAB2 | 0.492 | 122 | NO |
| MB-5350 | TBX3 | 0.153 | 85 | NO |
| MB-5350 | USH2A | 0.114 | 123 | NO |
| MB-5350 | CDH1 | 0.092 | 119 | NO |
| MB-5467 | NA | NA | NA | NO |
| MB-5521 | RYR2 | 0.391 | 115 | YES |
| MB-5521 | FOXP1 | 0.121 | 91 | YES |
| MB-5521 | DCAF4L2 | 0.466 | 174 | YES |
| MB-5521 | CASP8 | 0.429 | 35 | YES |
| MB-5521 | LDLRAP1 | 0.587 | 150 | YES |
| MB-5521 | DNAH11 | 0.106 | 104 | YES |
| MB-5521 | PTPRM | 0.444 | 207 | YES |
| MB-5521 | AHNAK | 0.428 | 243 | YES |
| MB-5521 | NF1 | 0.048 | 145 | YES |
| MB-5521 | FAM20C | 0.188 | 16 | YES |
| MB-5521 | SYNE1 | 0.153 | 85 | YES |
| MB-5521 | MEN1 | 0.124 | 145 | YES |
| MB-5521 | GLDC | 0.127 | 79 | YES |
| MB-5521 | PIK3CA | 0.06 | 84 | YES |
| MB-5521 | DNAH11 | 0.112 | 152 | YES |
| MB-5521 | MALAT1 | 0.493 | 211 | YES |
| MB-5521 | SBNO1 | 0.425 | 214 | YES |
| MB-5521 | NDFIP1 | 0.135 | 52 | YES |
| MB-5521 | STK11 | 0.571 | 182 | YES |
| MB-5521 | NEK1 | 0.628 | 43 | YES |
| MB-5521 | GATA3 | 0.083 | 132 | YES |
| MB-5521 | MLL2 | 0.504 | 129 | YES |
| MB-5521 | PRKG1 | 0.459 | 122 | YES |
| MB-5521 | ACVRL1 | 0.594 | 32 | YES |
| MB-5521 | PIK3R1 | 0.538 | 130 | YES |

| | | | | |
|---|---|---|---|---|
| MB-5521 | SYNE1 | 0.593 | 54 | YES |
| MB-5552 | TP53 | 0.129 | 85 | NO |
| MB-5552 | ASXL1 | 0.486 | 313 | NO |
| MB-5552 | TP53 | 0.165 | 109 | NO |
| MB-5552 | SF3B1 | 0.123 | 81 | NO |
| MB-5552 | COL12A1 | 0.335 | 194 | NO |
| MB-5552 | ALK | 0.421 | 216 | NO |
| MB-5552 | PDE4DIP | 0.106 | 492 | NO |
| MB-5552 | CLRN2 | 0.41 | 173 | NO |
| MB-5552 | USH2A | 0.162 | 191 | NO |
| MB-5552 | HERC2 | 0.432 | 139 | NO |
| MB-5552 | NT5E | 0.367 | 166 | NO |
| MB-5552 | PTEN | 0.152 | 66 | NO |
| MB-5552 | EGFR | 0.477 | 128 | NO |
| MB-5552 | THSD7A | 0.432 | 111 | NO |
| MB-5552 | DNAH11 | 0.536 | 330 | NO |
| MB-5552 | SBNO1 | 0.25 | 20 | NO |
| MB-5575 | PTPN22 | 0.4 | 90 | NO |
| MB-5575 | COL12A1 | 0.726 | 124 | NO |
| MB-5575 | ASXL1 | 0.643 | 252 | NO |
| MB-5575 | DNAH2 | 0.171 | 82 | NO |
| MB-5575 | USP28 | 0.3 | 10 | NO |
| MB-5575 | PIK3R1 | 0.329 | 82 | NO |
| MB-5575 | CACNA2D3 | 0.328 | 116 | NO |
| MB-5575 | COL22A1 | 0.122 | 131 | NO |
| MB-5575 | COL22A1 | 0.122 | 131 | NO |
| MB-5575 | NPNT | 0.333 | 96 | NO |
| MB-5575 | COL22A1 | 0.464 | 181 | NO |
| MB-5575 | TP53 | 0.162 | 74 | NO |
| MB-5653 | HERC2 | 0.237 | 38 | NO |
| MB-5653 | ARID1A | 0.218 | 307 | NO |
| MB-5653 | MAP3K1 | 0.589 | 95 | NO |
| MB-5653 | SMAD4 | 0.471 | 191 | NO |
| MB-5653 | AHNAK2 | 0.506 | 433 | NO |
| MB-5653 | LAMA2 | 0.197 | 147 | NO |
| MB-5653 | NOTCH1 | 0.484 | 153 | NO |
| MB-5653 | SBNO1 | 0.491 | 224 | NO |
| MB-5653 | PALLD | 0.543 | 243 | NO |
| MB-5653 | MYH9 | 0.533 | 45 | NO |
| MB-5653 | ARID1A | 0.223 | 305 | NO |
| MB-5653 | PIK3CA | 0.189 | 106 | NO |
| MB-5653 | RYR2 | 0.549 | 133 | NO |
| MB-5653 | NF1 | 0.456 | 250 | NO |
| MB-5653 | SYNE1 | 0.435 | 177 | NO |
| MB-5653 | SYNE1 | 0.218 | 293 | NO |
| MB-6060 | GPS2 | 0.696 | 135 | NO |
| MB-6060 | BRCA2 | 0.53 | 83 | NO |
| MB-6060 | ATR | 0.122 | 98 | NO |
| MB-6060 | FOXP1 | 0.543 | 35 | NO |
| MB-6060 | USH2A | 0.678 | 115 | NO |
| MB-6060 | DNAH5 | 0.37 | 92 | NO |
| MB-6060 | BRCA2 | 0.53 | 83 | NO |

| | | | | |
|---|---|---|---|---|
| MB-6060 | NF1 | 0.138 | 29 | NO |
| MB-6060 | PIK3CA | 0.481 | 189 | NO |
| MB-6060 | EP300 | 0.45 | 60 | NO |
| MB-6060 | FRMD3 | 0.482 | 56 | NO |
| MB-6060 | AHNAK2 | 0.177 | 379 | NO |
| MB-6060 | SYNE1 | 0.629 | 175 | NO |
| MB-6060 | MUC16 | 0.121 | 199 | NO |
| MB-6060 | COL6A3 | 0.609 | 174 | NO |
| MB-6060 | MLL2 | 0.444 | 90 | NO |
| MB-6060 | KMT2C | 0.553 | 38 | NO |
| MB-6060 | PDE4DIP | 0.301 | 103 | NO |
| MB-6060 | DNAH2 | 0.122 | 41 | NO |
| MB-6060 | HERC2 | 0.167 | 48 | NO |
| MB-6060 | USP9X | 0.131 | 61 | NO |
| MB-6060 | BRCA1 | 0.416 | 101 | NO |
| MB-6060 | L1CAM | 0.486 | 109 | NO |
| MB-6060 | UBR5 | 0.744 | 156 | NO |
| MB-6060 | SGCD | 0.67 | 97 | NO |
| MB-6060 | PDE4DIP | 0.34 | 241 | NO |
| MB-6060 | TP53 | 0.464 | 112 | NO |
| MB-6060 | SGCD | 0.173 | 52 | NO |
| MB-6060 | LIFR | 0.494 | 168 | NO |
| MB-6060 | CACNA2D3 | 0.722 | 18 | NO |
| MB-6060 | USH2A | 0.781 | 32 | NO |
| MB-6060 | MAP3K10 | 0.417 | 36 | NO |
| MB-6060 | SHANK2 | 0.652 | 267 | NO |
| MB-6060 | THSD7A | 0.438 | 16 | NO |
| MB-6077 | KMT2C | 0.112 | 260 | YES |
| MB-6077 | AHNAK | 0.509 | 273 | YES |
| MB-6077 | L1CAM | 0.536 | 84 | YES |
| MB-6077 | FANCA | 0.571 | 191 | YES |
| MB-6077 | PIK3CA | 0.204 | 206 | YES |
| MB-6077 | PDE4DIP | 0.233 | 511 | YES |
| MB-6077 | PTEN | 0.148 | 264 | YES |
| MB-6077 | RUNX1 | 0.444 | 160 | YES |
| MB-6077 | KMT2C | 0.387 | 181 | YES |
| MB-6077 | HERC2 | 0.489 | 225 | YES |
| MB-6077 | KDM3A | 0.135 | 37 | YES |
| MB-6077 | PTEN | 0.164 | 165 | YES |
| MB-6077 | MYH9 | 0.342 | 231 | YES |
| MB-6208 | AHNAK2 | 0.124 | 129 | YES |
| MB-6208 | PDE4DIP | 0.149 | 221 | YES |
| MB-6208 | AHNAK | 0.493 | 144 | YES |
| MB-6208 | FANCD2 | 0.125 | 64 | YES |
| MB-6208 | AHNAK2 | 0.41 | 205 | YES |
| MB-6208 | AHNAK2 | 0.128 | 125 | YES |
| MB-6208 | USH2A | 0.368 | 19 | YES |
| MB-6208 | CDKN2A | 0.48 | 25 | YES |
| MB-6208 | AHNAK2 | 0.15 | 273 | YES |
| MB-6208 | CDKN1B | 0.605 | 43 | YES |
| MB-6208 | FANCD2 | 0.337 | 190 | YES |
| MB-6208 | AHNAK | 0.42 | 69 | YES |

| | | | | |
|---|---|---|---|---|
| MB-6208 | SYNE1 | 0.607 | 168 | YES |
| MB-6208 | NCOA3 | 0.7 | 110 | YES |
| MB-6208 | LAMA2 | 0.277 | 94 | YES |
| MB-6208 | PIK3CA | 0.263 | 209 | YES |
| MB-6208 | AHNAK2 | 0.124 | 129 | YES |
| MB-6208 | AHNAK2 | 0.477 | 222 | YES |
| MB-6208 | USH2A | 0.652 | 92 | YES |
| MB-6208 | AHNAK2 | 0.184 | 141 | YES |
| MB-6208 | ASXL2 | 0.478 | 138 | YES |
| MB-6284 | USP9X | 0.28 | 157 | NO |
| MB-6284 | SIK1 | 0.542 | 190 | NO |
| MB-6284 | FOXP1 | 0.264 | 106 | NO |
| MB-6284 | BIRC6 | 0.406 | 101 | NO |
| MB-6284 | USP28 | 0.691 | 110 | NO |
| MB-6284 | UTRN | 0.338 | 74 | NO |
| MB-6284 | PALLD | 0.5 | 58 | NO |
| MB-6284 | TAF1 | 0.4 | 45 | NO |
| MB-6284 | PIK3R1 | 0.286 | 119 | NO |
| MB-6284 | COL12A1 | 0.5 | 80 | NO |
| MB-6284 | BIRC6 | 0.495 | 99 | NO |
| MB-6284 | MLLT4 | 0.571 | 70 | NO |
| MB-6284 | SMARCC2 | 0.522 | 67 | NO |
| MB-6287 | NCOR2 | 0.503 | 183 | PROBABLY |
| MB-6287 | CHD1 | 0.435 | 115 | PROBABLY |
| MB-6287 | TBX3 | 0.261 | 46 | PROBABLY |
| MB-6287 | ALK | 0.155 | 110 | PROBABLY |
| MB-6287 | NF1 | 0.407 | 59 | PROBABLY |
| MB-6287 | KDM3A | 0.543 | 92 | PROBABLY |
| MB-6287 | AKAP9 | 0.41 | 178 | PROBABLY |
| MB-6287 | CDH1 | 0.357 | 56 | PROBABLY |
| MB-6287 | NCOR2 | 0.358 | 81 | PROBABLY |
| MB-6287 | PIK3CA | 0.266 | 214 | PROBABLY |
| MB-6287 | MALAT1 | 0.413 | 167 | PROBABLY |
| MB-6287 | SEPT7P2 | 0.333 | 45 | PROBABLY |
| MB-6287 | LAMB3 | 0.8 | 60 | PROBABLY |
| MB-6287 | NDFIP1 | 0.491 | 159 | PROBABLY |
| MB-6287 | MYO3A | 0.354 | 127 | PROBABLY |
| MB-6287 | RASGEF1B | 0.549 | 71 | PROBABLY |
| MB-6287 | ERBB3 | 0.446 | 168 | PROBABLY |
| MB-7011 | COL6A3 | 0.405 | 205 | NO |
| MB-7011 | MYH9 | 0.21 | 62 | NO |
| MB-7011 | RYR2 | 0.8 | 65 | NO |
| MB-7011 | DNAH11 | 0.189 | 106 | NO |
| MB-7011 | AHNAK2 | 0.102 | 255 | NO |
| MB-7011 | DNAH5 | 0.667 | 30 | NO |
| MB-7011 | LAMA2 | 0.169 | 89 | NO |
| MB-7011 | AFF2 | 0.2 | 225 | NO |
| MB-7011 | RB1 | 0.294 | 17 | NO |
| MB-7011 | LAMA2 | 0.279 | 154 | NO |
| MB-7011 | ROS1 | 0.472 | 72 | NO |
| MB-7011 | AFF2 | 0.199 | 226 | NO |
| MB-7011 | ARID2 | 0.857 | 63 | NO |

| | | | | |
|---|---|---|---|---|
| MB-7043 | COL6A3 | 0.49 | 192 | NO |
| MB-7043 | SYNE1 | 0.485 | 99 | NO |
| MB-7043 | ERBB4 | 0.167 | 108 | NO |
| MB-7043 | GLDC | 0.455 | 101 | NO |
| MB-7043 | DNAH11 | 0.478 | 134 | NO |
| MB-7043 | AKT1 | 0.19 | 137 | NO |
| MB-7043 | ERBB2 | 0.629 | 213 | NO |
| MB-7043 | USH2A | 0.207 | 29 | NO |
| MB-7043 | USP28 | 0.368 | 38 | NO |
| MB-7043 | SF3B1 | 0.063 | 63 | NO |
| MB-7043 | LAMB3 | 0.517 | 118 | NO |
| MB-7043 | TG | 0.578 | 102 | NO |
| MB-7043 | USH2A | 0.117 | 180 | NO |
| MB-7043 | BIRC6 | 0.103 | 116 | NO |
| MB-7092 | FRMD3 | 0.692 | 52 | NO |
| MB-7092 | ATR | 0.521 | 328 | NO |
| MB-7092 | SHANK2 | 0.12 | 251 | NO |
| MB-7092 | ARID1A | 0.129 | 70 | NO |
| MB-7092 | AHNAK | 0.393 | 247 | NO |
| MB-7092 | SBNO1 | 0.227 | 203 | NO |
| MB-7092 | MALAT1 | 0.105 | 171 | NO |
| MB-7092 | TG | 0.477 | 199 | NO |
| MB-7092 | AKT1 | 0.472 | 322 | NO |
| MB-7092 | ERBB4 | 0.384 | 305 | NO |
| MB-7092 | DNAH5 | 0.511 | 184 | NO |
| MB-7092 | TBL1XR1 | 0.452 | 155 | NO |
| MB-7092 | PRKCZ | 0.48 | 298 | NO |
| MB-7092 | UTRN | 0.447 | 94 | NO |
| MB-7092 | APC | 0.44 | 25 | NO |
| MB-7092 | GATA3 | 0.22 | 218 | NO |
| MB-7092 | ERBB4 | 0.51 | 104 | NO |
| MB-7133 | DNAH2 | 0.686 | 51 | YES |
| MB-7133 | TG | 0.517 | 176 | YES |
| MB-7133 | NOTCH1 | 0.556 | 153 | YES |
| MB-7133 | TG | 0.296 | 115 | YES |
| MB-7133 | ARID5B | 0.543 | 70 | YES |
| MB-7133 | GLDC | 0.747 | 79 | YES |
| MB-7133 | GLDC | 0.31 | 116 | YES |
| MB-7133 | PTPN22 | 0.397 | 58 | YES |
| MB-7133 | GATA3 | 0.218 | 262 | YES |
| MB-7133 | PTPRM | 0.174 | 46 | YES |
| MB-7133 | TAF1 | 0.19 | 184 | YES |
| MB-7133 | ASXL2 | 0.629 | 62 | YES |
| MB-7133 | FOXO1 | 0.375 | 152 | YES |
| MB-7133 | DNAH11 | 0.571 | 84 | YES |
| MB-7292 | MYO1A | 0.541 | 37 | NO |
| MB-7292 | ERBB4 | 0.553 | 76 | NO |
| MB-7292 | MUC16 | 0.523 | 130 | NO |
| MB-7292 | PRKACG | 0.474 | 114 | NO |
| MB-7292 | STAB2 | 0.496 | 135 | NO |
| MB-7292 | NCOR2 | 0.561 | 66 | NO |
| MB-7292 | AKAP9 | 0.512 | 121 | NO |

| | | | | |
|---|---|---|---|---|
| MB-7292 | MALAT1 | 0.463 | 149 | NO |
| MB-7292 | FANCA | 0.4 | 70 | NO |
| MB-7299 | DNAH5 | 0.357 | 56 | YES |
| MB-7299 | PRKG1 | 0.429 | 63 | YES |
| MB-7299 | STAB2 | 0.169 | 65 | YES |
| MB-7299 | MUC16 | 0.55 | 149 | YES |
| MB-7299 | NOTCH1 | 0.663 | 101 | YES |
| MB-7299 | CHD1 | 0.143 | 28 | YES |
| MB-7299 | USH2A | 0.621 | 87 | YES |
| MB-7299 | FOXO3 | 0.41 | 61 | YES |
| MB-7299 | PDE4DIP | 0.111 | 459 | YES |
| MB-7299 | EP300 | 0.205 | 88 | YES |
| MB-7299 | EP300 | 0.266 | 154 | YES |
| MB-7299 | RYR2 | 0.658 | 38 | YES |
| MB-7299 | FANCA | 0.481 | 185 | YES |
| MB-7299 | PTPRM | 0.615 | 78 | YES |
| MB-7299 | BRIP1 | 0.361 | 83 | YES |
